# RasGRP1 agonism stimulates P-TEFb biogenesis via MEK-ERK-mTORC1 signaling to reverse HIV latency with minimal CD4 downregulation

**DOI:** 10.64898/2026.01.14.699500

**Authors:** Uri Mbonye, Ana Bellomo, Eleonora Elhalem, Lucia Gandolfi Donadio, Muda Yang, Ata Abbas, Amie Donner, Maria Julieta Comin, Jonathan Karn

## Abstract

Reactivating latent HIV to eliminate persisting infected cells requires non-toxic latency-reversing agents (LRAs) that activate P-TEFb, a cellular transcription factor essential for efficient HIV RNA synthesis. Diacylglycerol (DAG)-mimicking PKC agonists induce P-TEFb to reverse HIV latency mainly through a PKC-independent RasGRP1-Ras-Raf-MEK-ERK1/2 pathway, but also elicit global T-cell activation and a drastic downregulation of CD4 receptors. Here, we demonstrate that synthetic DAG-indololactones, which preferentially bind RasGRP1 over PKC by up to 60-fold, reverse HIV latency and robustly induce posttranscriptional P-TEFb expression in primary T cells via MEK-ERK1/2-mTORC1 signaling without triggering T-cell activation markers and with minimal CD4 loss. Elevation of T-cell activation markers by natural and synthetic PKC agonists proceeds through MEK-ERK1/2 but is independent of mTORC1 activity. Combining DAG-indololactones with HDAC inhibitors synergistically reactivates latent HIV in a primary T-cell model and reverses latency in CD4^+^ T cells from treated individuals at levels comparable to a potent PKC agonist. These findings suggest that a combination LRA approach targeting P-TEFb production through RasGRP1-ERK1/2-mTORC1 signaling and the epigenetic activation of proviral HIV can efficiently and safely reverse HIV latency.

## INTRODUCTION

In treated people with HIV (PWH), HIV-1 persists within tissue reservoirs and a small subset of resting memory CD4^+^ T cells (∼60-100 per 10^6^ cells) that harbor transcriptionally latent proviruses ^1–3^. Latently infected memory T cells evade host antiviral immune surveillance by producing minimal viral RNA and proteins. Mechanistic studies using ex vivo primary CD4^+^ T- cell models of HIV latency have shown that restrictive epigenetic structures at the proviral HIV promoter, along with very low expression of the host transcription elongation factor P-TEFb, are the main barriers to reactivating transcriptionally latent HIV in memory CD4^+^ T cells ^4–8^. Despite these insights, deliberate reactivation of latent HIV to facilitate clearance of these persistent infected memory T cells – the “Shock and Kill” strategy – has achieved limited success so far, partly because non-toxic latency-reversing agents (LRAs) capable of effectively mobilizing P-TEFb to proviral HIV remain to be identified.

P-TEFb is a nuclear heterodimer complex consisting of the CDK9 serine/threonine kinase and one of three cyclin T subunits. Although cyclin T1 (CycT1) is the primary regulatory partner for CDK9 and the only cyclin able to interact with HIV Tat to recruit P-TEFb to proviral HIV, the alternative cyclin T splice variants T2a and T2b might also support the transcription of cellular genes ^9–11^. The reduced levels of P-TEFb in resting CD4^+^ T cells is due to a posttranscriptional block of CycT1 mRNA translation caused by microRNAs, along with the proteasomal degradation of CycT1 protein ^12, 13^. The subcellular localization and molecular mechanisms that stabilize CycT1 mRNA in memory CD4^+^ T cells remain to be determined. One possibility is that existing CycT1 transcripts are confined to cytoplasmic ribonucleoprotein condensates, such as P bodies or stress granules, which are known to contain translationally repressed mRNAs ^14, 15^.

Restricted CycT1 expression in memory CD4^+^ T cells causes the CDK9 subunit, which is usually present, to be trapped in an inactive state in the cytoplasm by the kinase-specific chaperone complex Hsp90/Cdc37 ^6, 16^. Activation of memory T cells via the T-cell receptor (TCR) prompts rapid posttranscriptional production of CycT1, which is closely linked with the phosphorylation of CDK9 at Thr186 (pThr186 CDK9). This modification is crucial for stabilizing the heterodimeric formation of P-TEFb and enabling its kinase activity ^6, 8^. The coordination of the heterodimer by pThr186 CDK9 is also essential for incorporating P-TEFb into 7SK snRNP. This nucleoplasmic complex inhibits P-TEFb kinase activity and permits the enzyme to be delivered to genes that have begun transcription ^17^. In contrast, inducible phosphorylation of CDK9 by CDK7 at another conserved activation loop residue, Ser175 (pSer175 CDK9), was only observed in a subset of P-TEFb that has dissociated from 7SK snRNP and is unlikely to be in complex with BRD4, a bromodomain-containing protein thought to be the major recruiter of P-TEFb to cellular genes ^18^. Further functional analysis of pSer175 showed that, although unmodified Ser175 is crucial for enabling a key interaction between P-TEFb and BRD4, HIV Tat manipulates pSer175 to interact more effectively with the CDK9 activation loop, thereby promoting the competitive recruitment of P-TEFb to proviral HIV ^18^. Based on these observations, we recently developed a sensitive flow cytometry assay (P-TEFb immuno-flow) to detect transcriptionally active P-TEFb by identifying the co-expression of translated CycT1 and pSer175 CDK9 ^7, 19^.

By dissecting the complex T-cell receptor (TCR) signaling cascade in a primary T-cell model of HIV latency, alongside healthy donor-derived memory CD4^+^ T cells, we have previously defined the signaling pathways that are essential for P-TEFb biogenesis and enabling P-TEFb-dependent proviral reactivation. An unexpected finding was that, in response to TCR co-stimulation, the RasGRP1-Ras-Raf-MEK-ERK1/2 and PI3K-mTORC2-AKT-mTORC1 pathways can both induce the posttranscriptional synthesis of CycT1 and the generation of transcriptionally active P-TEFb ^7^. The central role of ERK signaling in reversing HIV latency through P-TEFb regulation was confirmed by the observation that protein kinase C (PKC) agonists, such as prostratin and ingenol, stimulate CycT1 protein synthesis and reactivate latent HIV primarily via the RasGRP1-Ras-Raf-MEK-ERK1/2 pathway rather than through PKC enzymes. These natural PKC agonists are functional mimics of the lipid second messenger diacylglycerol (DAG), which broadly bind to proteins containing a C1 domain, leading to their membrane recruitment and activation of their enzymatic functions. The activation of conventional and novel PKC enzymes through their respective C1 domains is also known to activate NF-κB via the canonical pathway and can cause rapid downregulation of the CD4 receptor ^20–23^. Our findings therefore indicate that bypassing PKC activation by specifically targeting the RasGRP1 C1 domain can elevate P-TEFb expression through ERK1/2 signaling, in a way that is sufficient to reverse HIV latency with minimal unintended global T-cell activation responses.

Activation of mTORC1 through the PI3K-mTORC2-AKT pathway is widely regarded as the main mechanism for the initiation of protein synthesis in most cell types, including T cells. Phosphorylation of ribosomal protein S6 kinase (p70S6K) by mTORC1 initiates multiple phosphorylation events on components of the translation machinery, including the ribosomal protein S6 (rpS6) and translation initiation and elongation factors ^24^. Since DAG-mimicking C1 domain agonists are not known to signal through PI3K or AKT, it is unclear how they are able to reverse the translational repression of CycT1 mRNA via the MAPK ERK signaling pathway.

Here, we demonstrate that a class of synthetic DAG-mimetic compounds called DAG-indololactones, which preferentially bind the C1 domain of RasGRP1 over the PKC enzymes, can stimulate CycT1 mRNA translation and produce transcriptionally active P-TEFb in memory CD4^+^ T cells with minimal induction of the T-cell activation markers CD25 and CD69, as well as minimal loss of CD4. Whereas PKC agonists were found to reactivate latent HIV through ERK-dependent, complementary actions of mTORC1 and RSK, DAG-indololactones reactivated latent HIV via ERK-mTORC1 signaling. Combinations of the DAG-indololactone 2A127 and HDAC inhibition (HDACi) demonstrated synergy in reactivating latent HIV in an ex vivo primary T-cell model and effectively reversed HIV latency in memory CD4^+^ T cells from treated PWH. While natural and synthetic PKC agonists caused rapid, selective, and significant downregulation of the CD4 receptor on uninfected memory CD4^+^ T cells, through a process deemed to be independent of RasGRP1-MEK-ERK1/2 signaling, the combination of 2A127 and HDACi minimally affected the expression of CD4 or other surface receptors tested. Our findings highlight the specific process through which RasGRP1-directed signaling can reactivate latent HIV in primary T cells and support the idea that a dual LRA strategy targeting P-TEFb biogenesis and the epigenetic activation of the HIV promoter can effectively and safely reverse HIV latency.

## RESULTS

### HIV latency reversal requires CycT1 translation and assembly of nucleoplasmic P-TEFb

There is strong evidence from both our ex vivo primary cell model of HIV latency and latently infected Jurkat T cells that P-TEFb is essential for reactivating latent HIV and initiating the production of the viral trans-activator Tat. In both models, proviral reactivation in response to a wide variety of latency-reversing agents (LRAs) or stimuli, including HDAC inhibitors, is abrogated by the P-TEFb kinase inhibitor flavopiridol (**Supplementary Figs. 1A-E**). Therefore, defining the cellular mechanisms that govern P-TEFb expression and activity in primary T cells is central to developing an effective latency-reversal strategy.

Although P-TEFb is constitutively expressed in actively dividing cells and cell line models, its abundance is severely limited in quiescent primary T cells due to translational repression (**Fig. 1A**) and proteolytic turnover of the CycT1 subunit ^12, 13, 25^. Activation of resting memory CD4^+^ T cells triggers rapid protein induction, accompanied by the nuclear localization of P-TEFb and nucleoplasmic assembly of 7SK snRNP (**Figs. 1A, 1B, 1F and Supplementary Fig. 1F**). To determine the primary regulatory checkpoint governing CycT1 restriction during cellular quiescence, we examined CycT1 mRNA levels, protein expression, and protein turnover in resting versus T-cell receptor (TCR)-activated memory CD4^+^ T cells. RNA FISH revealed that CycT1 transcript levels increase only modestly – by 1.44-and 1.33-fold at 4 h and 24 h after TCR co-stimulation, respectively (**Fig. 2A and Supplementary Fig. 1F**). This observation aligns with our previous bulk RNA-seq measurements across primary memory CD4^+^ T cells and polarized effector subsets ^7^. In contrast, immunofluorescence microscopy of these cells showed a 3-fold and 14.7-fold increase in CycT1 protein expression at 4 h and 24 h after TCR activation, respectively (**Fig. 1B, Supplementary Fig. 1F and Fig. 2A**). Similarly, using immunofluorescence flow cytometry, we observed an 8.5-fold and 16.7-fold increase in the percentage of CycT1+ cells at 4 h and 24 h after TCR activation, respectively (**Fig. 1F**). To test whether constitutive proteasomal degradation accounts for the low abundance of CycT1 in resting cells, we treated unstimulated memory CD4^+^ T cells with the proteasomal inhibitor MG132. While MG132 treatment over 8 h yielded a modest accumulation of both CycT1 and the rapidly turned over cell-cycle control Cyclin D3 (CycD3), it failed to recapitulate the robust expression driven by TCR co-stimulation (**Supplementary Fig. 2**). Conversely, blocking de novo protein synthesis with cycloheximide (CHX) during TCR activation completely abolished CycT1 induction (**Supplementary Figs. 2B and 2C**). Furthermore, CHX chase assays performed following TCR activation revealed that once synthesized, CycT1 exhibits negligible decay over a 2-hour chase window, in sharp contrast to CycD3, which lost over 50% of its abundance in a proteasomal-dependent manner (**Supplementary Figs. 3A and 3B**). Together, these findings establish that relief of translational repression, rather than transcriptional induction or baseline protein turnover is the primary rate-limiting checkpoint governing CycT1 expression in primary T cells. Because resting memory CD4^+^ T cells maintain this posttranscriptional block on P-TEFb biogenesis, they provide an intracellular environment conducive to maintaining latent HIV reservoirs in treated PWH.

**Figure 1.**
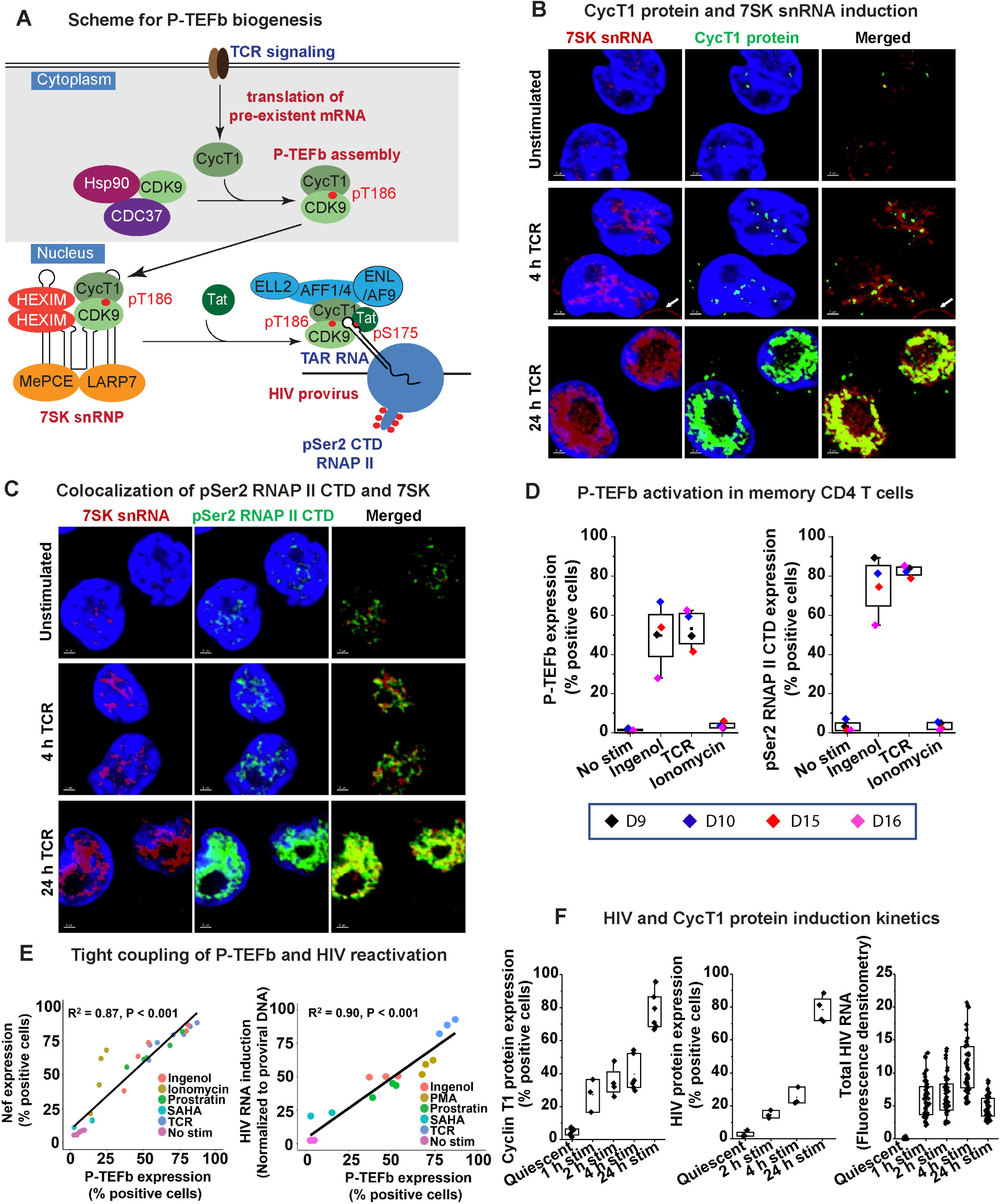
Emergence of HIV from latency is tightly coupled to the biogenesis of P-TEFb in primary T cells. (A) Scheme showing the signal-dependent translation of cyclin T1 (CycT1), its nuclear import following assembly with CDK9 to form P-TEFb and the incorporation of P-TEFb into 7SK snRNP. (B) RNA FISH and immunofluorescence staining of donor D9 memory CD4^+^ T cells to examine the subcellular co-expression of 7SK snRNA and the CycT1 subunit of P-TEFb following TCR activation for 4 h or 24 h. The white arrow points to a non-specific signal detected in the Cy5 (Red) channel emanating from an activator bead coated with anti-CD3 and anti-CD28 antibodies. (C) RNA FISH and immunofluorescence staining of donor D16 memory CD4^+^ T cells to examine the subcellular co-expression of 7SK snRNA and pSer2 RNAP II CTD following TCR activation for 4 h or 24 h. (D) *Left graph*, Assessment of transcriptionally active P-TEFb expression as measured by monitoring the dual expression of CycT1 and pSer175 CDK9. *Right graph*, Measurement of the phosphorylation of RNA polymerase II (RNAP II) at Ser2 of the heptad repeats of its C-terminal domain (pSer2 RNAP II CTD). Plotted experiments were conducted with memory CD4^+^ T cells from four different healthy donors as shown. Representative flow plots of these graphed data are shown in Supplementary Fig. 2A. The concentrations of stimuli used were as follows: 50 nM ingenol-3-angelate (ingenol); 1:1 bead-to-cell ratio anti-CD3/CD28 Dynabeads; and 1 µg/ml ionomycin. (E) Correlative expression profiles of P-TEFb and HIV Nef (*Left plot*) or poly-A unspliced HIV RNA (*Right plot*) showing that the extent of proviral reactivation in primary CD4^+^ T cells strongly correlates with the extent of inducible P-TEFb expression. Latently infected primary Th17 cells prepared ex vivo with the QUECEL procedure using naïve CD4^+^ T cells and derived from multiple donors (D14, D16, D17 and D18) were challenged for 24 h with the treatments shown. Correlation coefficients (R^2^) and statistical significance based on a Student’s t-test are shown. Plotted flow data are from two or more biological replicates where each experimental treatment was conducted with cells derived from at least two donors. The HIV RNA RT-qPCR data plotted are from three technical replicates conducted using cells from the same donor (D23). The concentrations of stimuli used were as follows: 50 nM ingenol; 1 µg/ml ionomycin; 1 µM prostratin; 500 nM SAHA; 1:1 bead-to-cell ratio anti-CD3/CD28 Dynabeads; 50 ng/ml PMA. (F) Comparison of expression profiles of CycT1 protein, HIV proteins and HIV RNA following primary T cell activation. HIV RNA was examined by RNA FISH microscopy. CycT1 protein was measured by immunofluorescence flow cytometry. Plotted flow cytometry data for CycT1 expression are compiled from seven independent experiments, including two complete time-course experiments, two biological replicates using primary memory CD4+ T cells from healthy donors (D14 and D16), and one experiment using *ex vivo*-generated QUECEL primary Th17 cells. Plotted flow cytometry data for HIV protein expression are compiled from four independent experiments: two measuring Tat and two measuring Nef. Plotted data for HIV RNA FISH are from the quantification of fluorescence signal from 50 imaged QUECEL primary Th17 cells which were prepared using naïve T cells from donor D27. Representative images and histogram flow plots of these graphed data are shown in Supplementary Figs. 4B-E.

**Figure 2.**
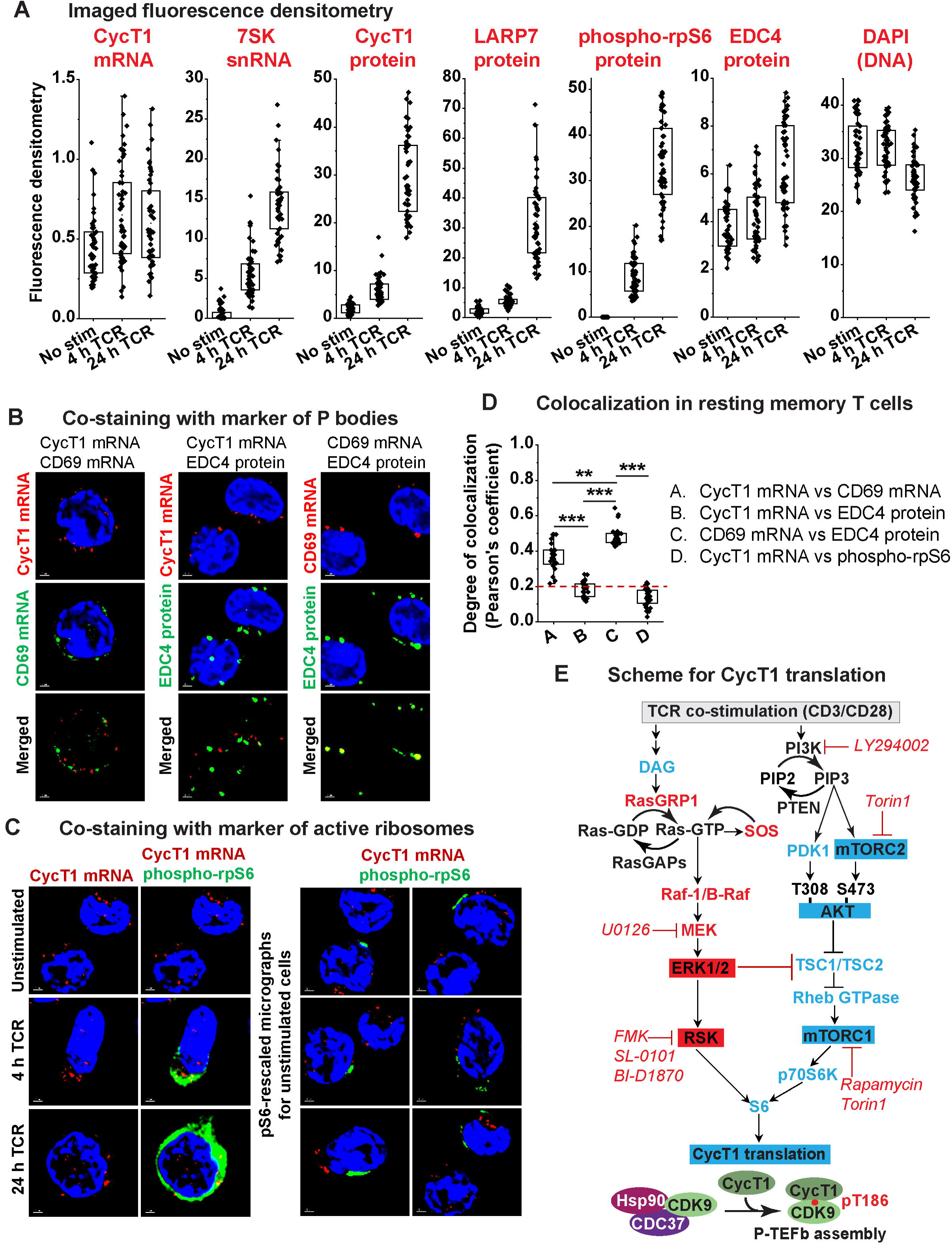
CycT1 mRNA is translationally repressed in unstimulated memory CD4^+^ T cells, but not associated with P bodies. (A) Quantification of the imaged fluorescence signal captured by 100x microscopy in unstimulated memory CD4^+^ T cells and following TCR activation for 4 h and 24 h. Representative plotted data are all from imaging experiments conducted using memory CD4^+^ T cells belonging to donor D16. Each graph shows the quantification of fluorescence signal from 50 imaged cells. (B) CycT1 mRNA exhibits little colocalization with EDC4, a marker of P bodies. *Left panel*, RNA FISH co-staining of CycT1 (red) and CD69 (green) mRNA. *Middle panel*, Combined RNA FISH and immunofluorescence staining of CycT1 mRNA (red) and EDC4 (green) protein. *Right panel*, Combined RNA FISH and immunofluorescence staining of CD69 mRNA (red) and EDC4 (green) protein. All three imaging experiments were conducted using unstimulated memory CD4^+^ T cells from donor D23. (C) *Left panel*, RNA FISH and immunofluorescence co-staining of CycT1 mRNA (red) and phospho-rpS6 (green) in unstimulated D16 memory CD4^+^ T cells and following their TCR activation for 4 h or 24 h. *Right panel*, Micrographs of unstimulated memory CD4^+^ T cells with less stringent scaling of the phospho-rpS6 signal to identify the subcellular localization of active ribosomes relative to that of CycT1 mRNA. (D) Utilization of the Pearson’s correlation coefficient (PCC) to determine the degree of colocalization between the factors shown in sets A-D in unstimulated D16 and D23 memory CD4^+^ T cells. Statistical significance was calculated using a two-tailed Student’s t test. Two asterisks indicate a *p* value < 10^−5^. Three asterisks indicate a *p* value < 10^−10^. The red dashed line identifies the PCC = 0.2 threshold which demarcates poor versus meaningful correlation. (E) Scheme showing proposed signaling mechanisms that regulate the ERK-dependent translation of CycT1 and P-TEFb assembly in memory CD4^+^ T cells. Inhibitors used in the current study are shown in red italics.

**Figure 3.**
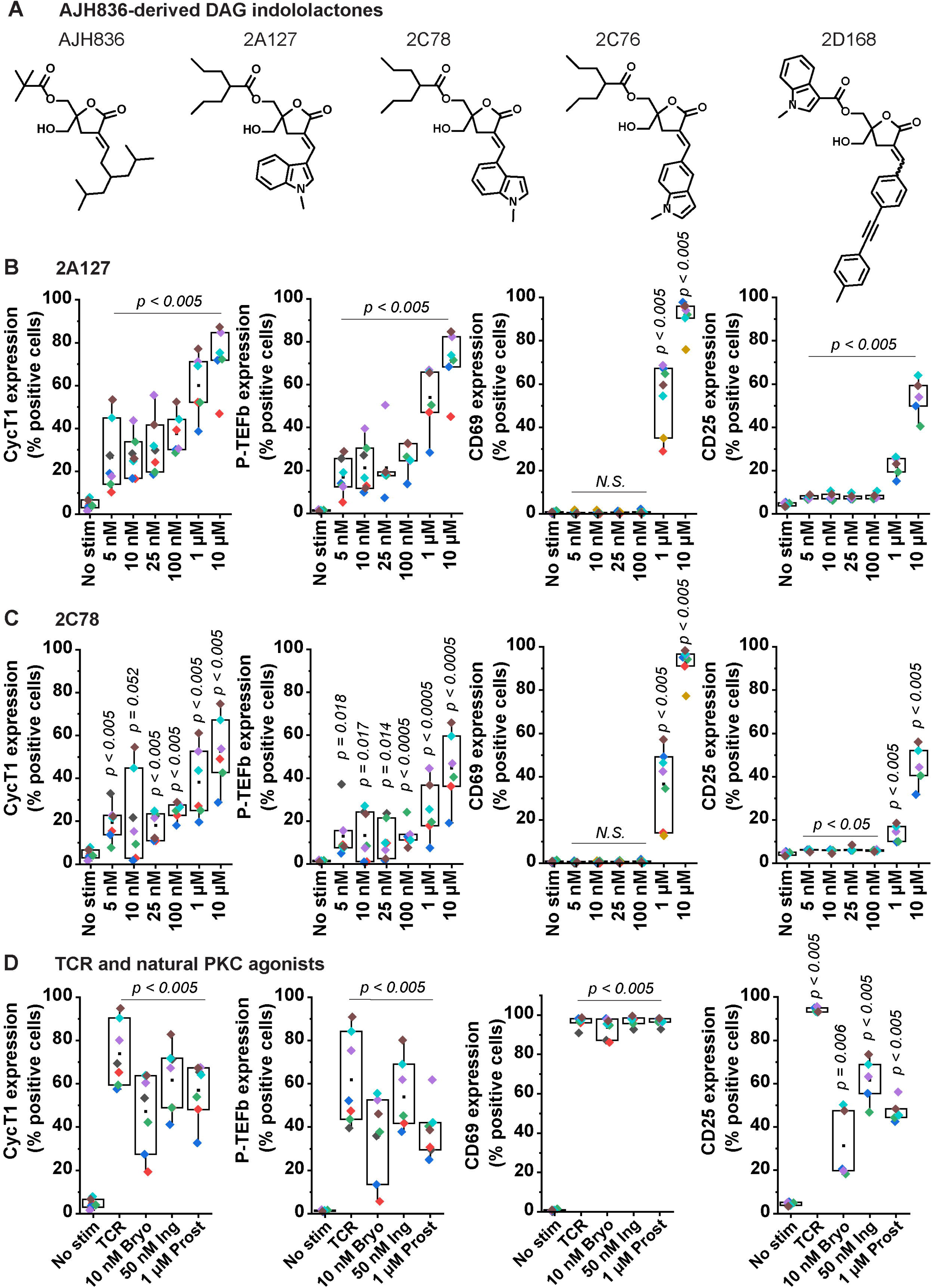
DAG-indololactones stimulate CycT1 translation and induce active P-TEFb with minimal elevation of T-cell activation markers. (A) Structures of the parent DAG-lactone AJH836 and DAG-indololactone derivatives. (B-D) The DAG-indololactones 2A127 and 2C78 stimulate CycT1 translation and induce active P-TEFb with minimal elevation of CD69 and CD25. Memory CD4^+^ T cells derived from two donors, an adult male (D23) and adult female (D24), were treated with varying concentrations of 2A127 or 2C78, or stimulated through the TCR or with natural PKC agonists for 24 h prior to immunostaining for transcriptionally active P-TEFb (dual staining for CycT1 and pSer175 CDK9), CD69 and CD25. Fluorescently stained cells were then analyzed by flow cytometry. Plotted data are from at least five independent experiments. Cyan and brown data points are from two independent experiments conducted with donor D24 cells; the other data points are from independent experiments carried out with donor D23 cells. Statistical significance was calculated for each treatment relative to the untreated condition (No stim) using a two-tailed Student’s t test. *N.S.* (not statistically significant) indicates a *p* value > 0.05. Representative 2D dot plots of these graphed flow data are shown in Supplementary Fig. 9D.

Following its signal-dependent translation, CycT1 rapidly localizes to the nucleus, assembling with CDK9 to form P-TEFb and incorporating into the 7SK snRNP complex (**Figs. 1A, 1B, 1F and Supplementary Fig. 1F**). Using our P-TEFb immuno-flow assay – which monitors transcriptionally active P-TEFb via the coordinated expression of CycT1 and phosphorylation of CDK9 at Ser175 (pSer175 CDK9) ^7, 19^ – we demonstrated that inducible P-TEFb biogenesis in memory CD4^+^ T cells from four healthy donors closely parallels the phosphorylation of RNA polymerase II (RNAP II) at Ser2 of its C-terminal domain heptad repeats (pSer2 RNAP II CTD) (**Fig. 1D and Supplementary Fig. 4A**). Because this pSer2 RNAP II CTD modification, primarily catalyzed by P-TEFb kinase ^26, 27^, serves as a marker of processive transcription elongation, it can be used to identify nucleoplasmic regions occupied by actively transcribing genes (**Fig. 1C**). Although 7SK snRNP sequesters P-TEFb in a catalytically inactive state ^28–31^, it also stabilizes P-TEFb retention within the nucleus to facilitate a convenient exchange of the enzyme to poised promoters ^32^. Upon TCR stimulation of memory CD4^+^ T cells, 7SK snRNA and its integral partner LARP7 adopt a doughnut-shaped nucleoplasmic distribution that is strongly coincident with elongation-competent pSer2 RNAP II CTD (**Fig. 1C and Supplementary Figs. 1G and 1H**). Therefore, 7SK snRNP is localized to a subnuclear region or phase-separated condensate that is rich in actively transcribing class II genes, facilitating the exchange of P-TEFb onto transcriptionally poised genes.

Consistent with the essential role of P-TEFb in proviral transcription elongation, LRA-induced P-TEFb activation strongly correlated with latent viral reactivation, as measured by immunofluorescence detection of Nef protein and RT-qPCR quantification of unspliced viral transcripts in a well-characterized ex vivo primary cell model known as the QUECEL (**Fig. 1E**). Kinetic profiling between 1 h and 4 h after T-cell activation further revealed that the onset of CycT1 protein expression mirrors the rapid induction of HIV RNA synthesis **Fig. 1F and Supplementary Fig. 4B)**. Inducible viral transcripts peaked at 4 h post-stimulation and noticeably declined by 24 h – likely reflecting an equilibrium between viral RNA synthesis and turnover – yet remained elevated 3- to 5-fold above unstimulated controls at 24 h (**Fig. 1F and Supplementary Fig. 1C**). Importantly, this sustained transcript pool supported continuous viral translation: time-course flow cytometry analysis in reactivated QUECEL primary Th17 cells demonstrated that viral proteins become detectable within 2 h of stimulation and progressively accumulate to peak at 24 h (**Fig. 1F and Supplementary Figs. 4C-4H**). Notably, among the representative cellular transcripts examined in bulk RNA-seq datasets from healthy donor memory CD4^+^ T cells and QUECEL effector subsets (SRA accessions SRP026389 and SRP145508), the expression profile of HIV RNA (measured by RNA FISH and RT-qPCR) most closely mirrored that of the Super Elongation Complex (SEC) component ELL2 (**Fig. 1 and Supplementary Figs. 5A and 5B**).

**Figure 4.**
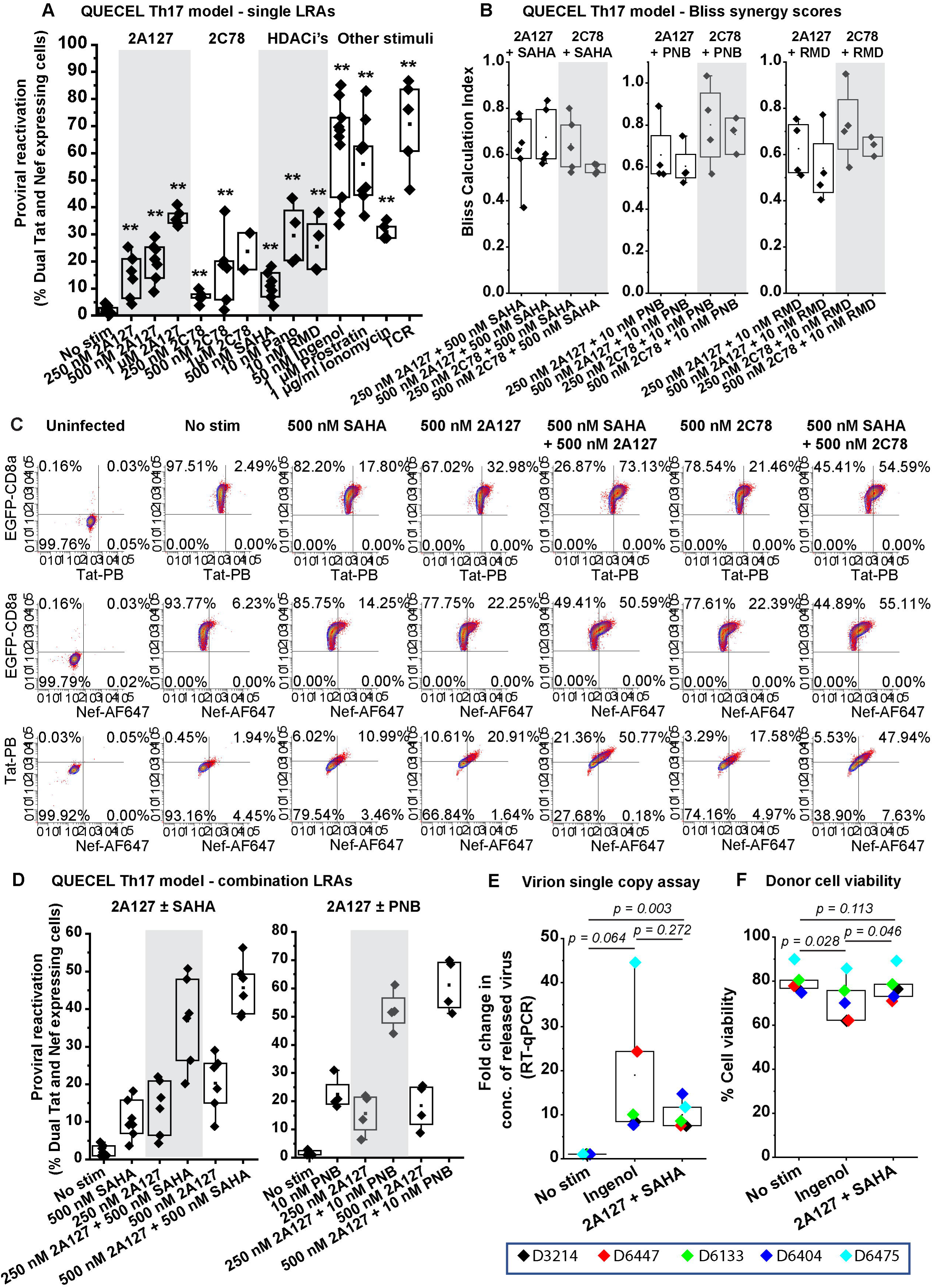
DAG-indololactones synergize with HDAC inhibitors to reactivate latent HIV in primary CD4^+^ T cells. (A) Viral reactivation measurements of individual LRA treatments in the QUECEL primary Th17 cell model. Latently infected primary Th17 cells were challenged for 24 h with varying concentrations of the DAG indololactones 2A127 or 2C78 or the HDAC inhibitors SAHA, panobinostat (PNB), or romidepsin (RMD). For comparison, cells were also activated through the TCR or treated with ingenol or ionomycin. Proviral reactivation was examined by measuring the dual expression of Tat and Nef using immunofluorescence flow cytometry. Plotted flow data are from two or more biological replicates where each experimental treatment was conducted with QUECEL primary Th17 cells generated using naïve T cells from at least two donors. Statistical significance was calculated relative to the untreated condition (No stim) using a two-tailed Student’s t test. Two asterisks indicate a *p* value < 0.0005. (B) Combination treatments with SAHA (*Left graph*), PNB (*Middle graph*) or RMD (*Right graph*) elicit effective synergy in viral reactivation in the QUECEL primary Th17 cell model. Plots of the calculated Bliss scores for the evaluation of synergy between DAG indololactones and HDAC inhibitors are shown with calculation indices of <1 deemed to be evidence of synergy. Plotted Bliss synergy data are from at least four independent experiments conducted with QUECEL primary Th17 cells generated using naïve T cells derived from two donors (D23 and D24). (C) Two-dimensional plots showing latent HIV reactivation profiles of individual or combination treatments as assessed by dual Tat and Nef immunofluorescence flow cytometry. (D) Combination of 2A127 and SAHA (*Left graph*) or 2A127 and PNB (*Right graph*) elicit effective synergy in viral reactivation in the QUECEL primary Th17 cell model. Plotted flow data are from four or more independent experiments conducted with QUECEL primary Th17 cells generated using naïve T cells derived from two donors (D23 and D24). (E and F) Combination of 2A127 and SAHA reactivates latent HIV in memory CD4^+^ T cells obtained from five treated PWH with minimal effects on cell viability. Following 24-h treatments as shown, latent HIV reactivation was examined by measuring virus release using an RT-qPCR RNA single copy assay targeting the integrase region of HIV. At least three qPCR repeats were performed for each donor; averages of these technical repeats are plotted in (E). Cell viability measurements based on propidium iodide staining are shown in (F). Statistical significance values shown were calculated using a two-tailed Student’s t test.

**Figure 5.**
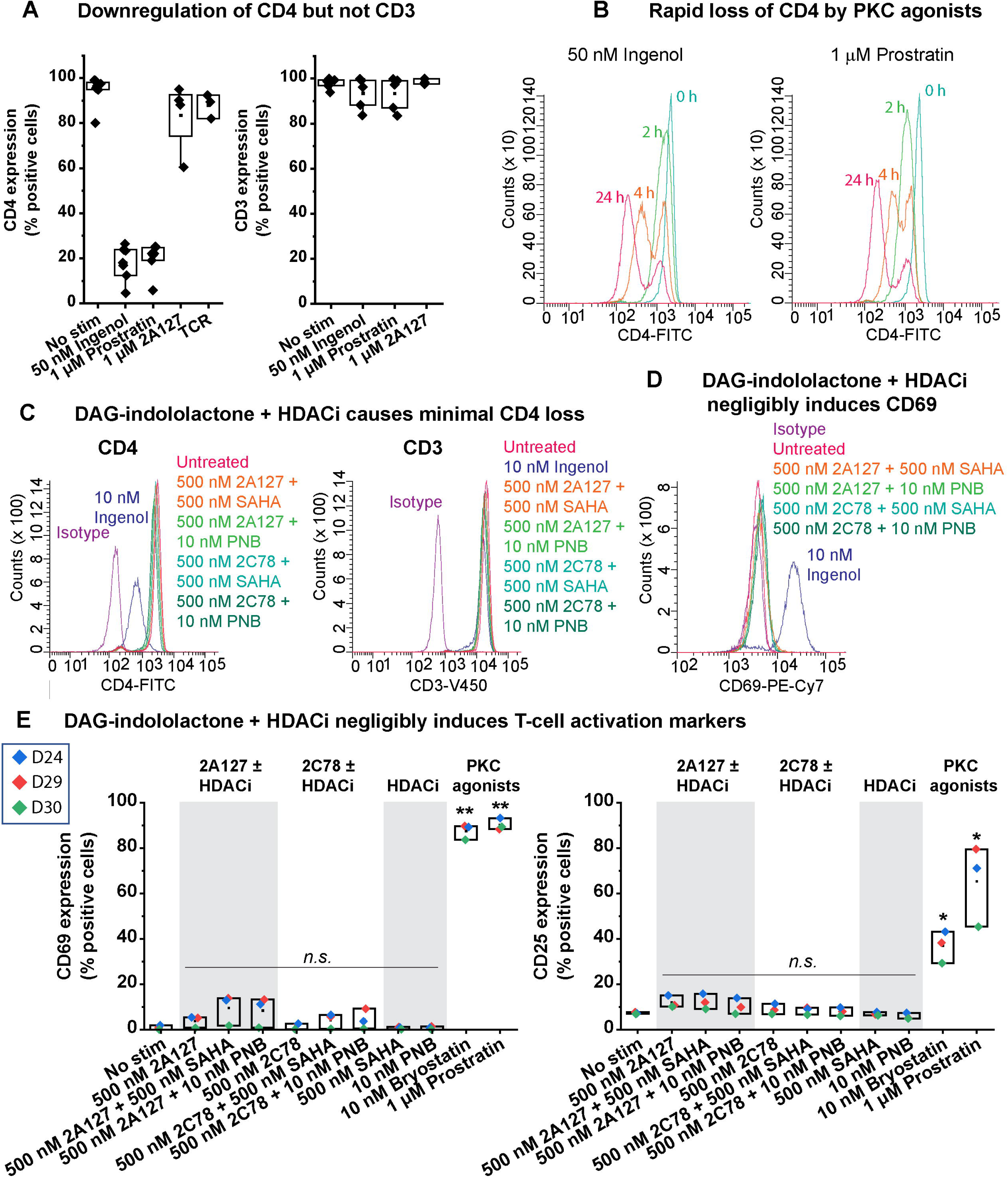
Selective downregulation of CD4 on memory T cells by PKC agonists, but not DAG-indololactones. (A) Ingenol and prostratin downregulate CD4 without adversely affecting CD3 expression. Memory CD4^+^ T cells were treated with 50 nM ingenol, 1 µM prostratin or activated through the TCR for 24 h prior to examination of CD4 and CD3 expression by immunofluorescence flow cytometry. Plotted flow data are from three or more independent experiments conducted with cells derived from at least two healthy donors. A total of five healthy donors (D23, D24, D25, D27, and D30) were sampled. (B) Rapid loss of CD4 expression following treatment of D25 memory CD4^+^ T cells with 50 nM ingenol or 1 µM prostratin. (C and D) Combination treatments of DAG-indololactones and HDAC inhibitors minimally impact CD4 and CD69 expression in D30 memory CD4^+^ T cells. (E) Combination treatments of DAG-indololactones and HDAC inhibitors minimally impact CD69 and CD25 expression in memory CD4^+^ T cells. Plotted flow data are from three independent experiments performed with cells derived from donors D24, D29 and D30. Statistical significance was calculated for each treatment relative to the untreated condition (No stim) using a two-tailed Student’s t test. Two asterisks indicate a *p* value < 0.0005, one asterisk indicates a *p* value < 0.05, and *n.s.* (not statistically significant) indicates a *p* value > 0.05.

### Translationally repressed CycT1 transcripts are not associated with P bodies

To further dissect posttranscriptional regulation in primary T cells, we compared CycT1 with the early T-cell activation marker CD69. Even though CD69 mRNA is rapidly and transiently induced following T-cell activation (based on an examination of a publicly available bulk RNA-seq dataset generated using unstimulated and TCR-activated memory CD4^+^ T cells ^33^; **Supplementary Fig. 5A**), like CycT1, CD69 mRNA is well established to be translationally repressed in resting T cells and rapidly mobilized upon activation ^25, 34, 35^. For instance, it has previously been shown that early CD69 protein induction following T cell activation is insensitive to transcriptional blockade with the RNAP II inhibitor actinomycin D ^34^, a finding that we can confirm in the current study: Treating memory CD4^+^ T cells with the covalent CDK7 inhibitor THZ1 significantly suppressed the 24-h transcriptional induction of the IL-2 receptor alpha subunit CD25, but had minimal effect on CD69 protein elevation (**Supplementary Figs. 6A–C**). Even the rapid (2 h) induction of CD69 protein was refractory to either THZ1 and actinomycin D treatment, whereas it was strongly suppressed by combined inhibition of MEK and PKC-θ (**Supplementary Fig. 6D**). Notably, while 100 nM FVP abrogated P-TEFb-dependent latent HIV reactivation (**Supplementary Figs. 1A–E**), the same concentration of FVP did not impair CD25 and CD69 expression (**Supplementary Fig. 6C**), confirming pathway-specific sensitivity rather than global transcriptional arrest and highlighting the heightened dependence of HIV on P-TEFb activity. Collectively, these data demonstrate that while CD25 inducible expression requires de novo transcription, the signal-dependent induction of both CycT1 and CD69 in memory CD4^+^ T cells relies on the translation of pre-existing mRNA pools.

We next performed combined RNA FISH and immunofluorescence imaging in unstimulated memory CD4^+^ T cells to determine whether translationally repressed CycT1 and CD69 transcripts reside within processing (P) bodies, which are cytoplasmic ribonucleoprotein condensates associated with mRNA storage and decay ^15^. Both CycT1 and CD69 mRNA were readily detectable by RNA FISH predominantly in the cytoplasm of unstimulated memory CD4^+^ T cells (**Figs. 2B, 2C and Supplementary Fig. 1I**), consistent with their detection by RNA-seq (baseline average quantitation of approximately 20 transcripts per million for either mRNA; **Supplementary Fig. 5**) . CD69 transcripts showed a bipartite distribution pattern in unstimulated memory T cells. One subset of CD69 mRNA appeared to colocalize with CycT1 mRNA, with a modest Pearson’s correlation coefficient (PCC) of 0.37 (**Figs. 2B, 2D and Supplementary Fig. 1I**). By contrast, a second group showed a distinct colocalization with EDC4 protein (PCC = 0.49), which is considered a reliable marker of P bodies ^36–39^ (**Figs. 2B, 2D and Supplementary Fig. 1I**). In unstimulated memory T cells, CycT1 mRNA did not significantly colocalize with EDC4 (PCC = 0.19) (**Figs. 2B and 2D**) or with phospho-rpS6 (PCC = 0.14), a marker of active ribosomes ^40^ (**Figs. 2C and 2D**); a PCC measurement of ≤ 0.2 is well established to be indicative of a poor or negligible degree of colocalization ^41, 42^. Thus, CD69 mRNA can be found in P bodies, whereas CycT1 transcripts are translationally repressed in cytoplasmic regions of primary T cells that are distinct from P bodies.

### Synthetic RasGRP1 agonists stimulate CycT1 translation and formation of active P-TEFb

As summarized in the scheme shown in **Fig. 2E**, by dissecting the complex array of TCR signaling pathways using inhibitor-based and transcriptomic approaches, we previously showed that two signaling arms, the RasGRP1-Ras-Raf-MEK-ERK1/2 pathway, activated by phospholipase C-γ mobilization of the lipid second messenger diacylglycerol (DAG), and the PI3K-mTORC2-AKT-mTORC1 pathway, complement each other in mediating P-TEFb expression in primary CD4^+^ T cells ^7^. In contrast, activation of PKC enzymes and stimulation of the MEKK1-MKK4/7-JNK-c-Jun MAPK pathway were found to be unnecessary for producing P-TEFb or even reactivating latent HIV in primary T cells. Naturally occurring phorbol esters, specifically ingenol-3-angelate (ingenol), prostratin, and bryostatin 1 (bryostatin), structurally mimic DAG and strongly bind the C1 domains of PKC and RasGRP, leading to their recruitment to the inner leaflet of the plasma membrane. By utilizing a MEK inhibitor (U0126), an inhibitor of PKC-θ (Sotrastaurin), which is considered the functionally predominant PKC isoform in T cells ^43–45^, and a pan-PKC inhibitor (Ro-31-8220), we were able to affirm that both ingenol and prostratin stimulate the posttranscriptional expression of CycT1 primarily via the MEK-ERK1/2 pathway rather than through PKC enzymes. CycT1 protein synthesis in primary T cells induced by ingenol or prostratin was significantly inhibited by U0126 and was minimally affected by treatment with PKC inhibitors. (**Supplementary Figs. 7A and 7B**). In contrast, U0126 modestly suppressed CycT1 protein synthesis induced by TCR co-stimulation (**Supplementary Fig. 7A**). This aligns with our previous report that TCR-activated biogenesis of P-TEFb is mediated by the combined actions of the MEK-ERK1/2 and PI3K-mTORC2-AKT-mTORC1 pathways ^7^. Posttranscriptional synthesis of CD69 and the transcriptional induction of CD25 by prostratin and bryostatin were also found to be largely dependent on ERK1/2 and much less dependent on PKC-θ activity (**Supplementary Figs. 7C-G**). While the induction of CD25 by ingenol was entirely MEK-dependent, the induction of CD69 by ingenol was partially inhibited by the MEK inhibitor but nearly eliminated by the combined inhibition of MEK and PKC-θ (**Supplementary Figs. 7C, 7D and 7G**).

**Figure 6.**
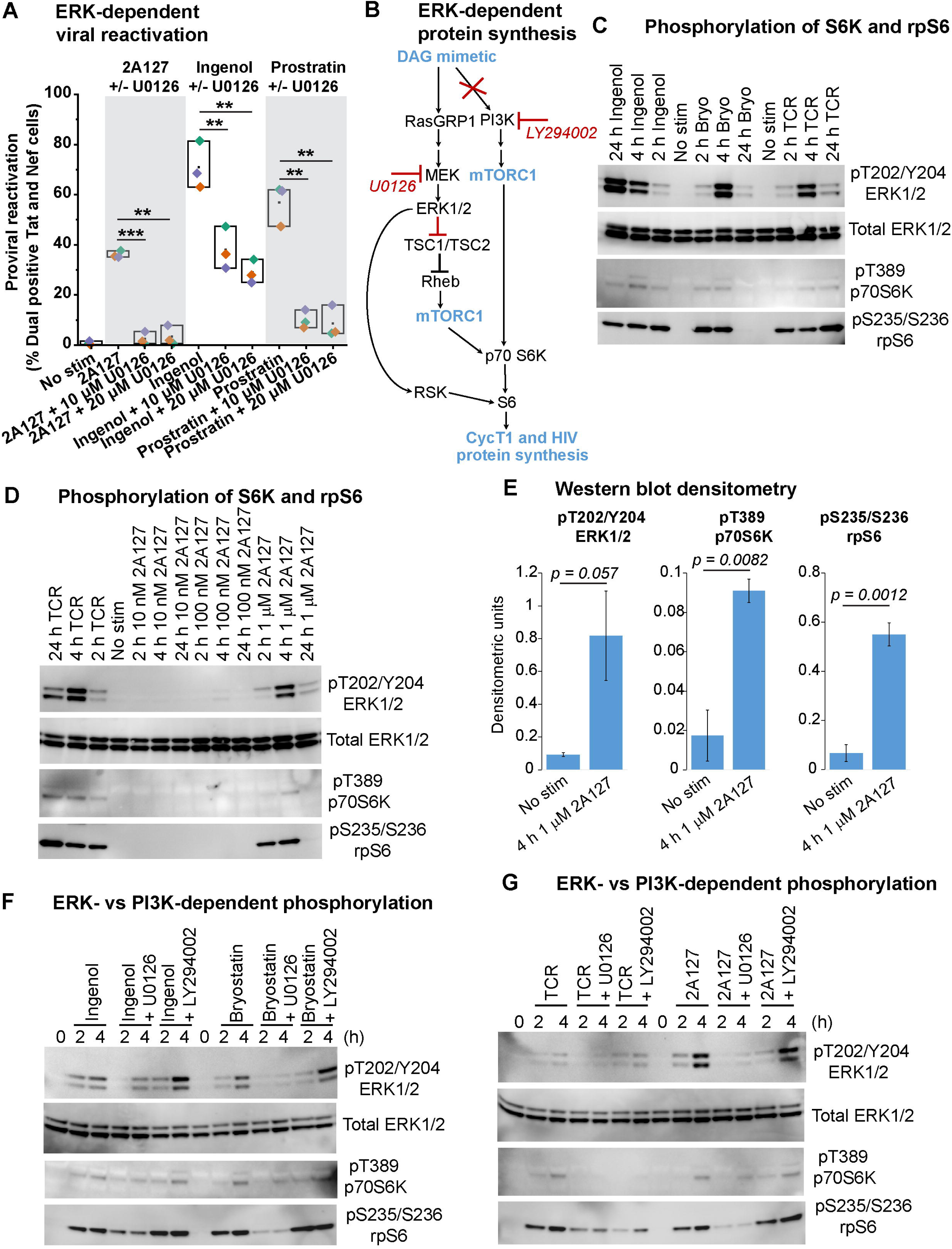
DAG-mimicking C1 domain agonists bypass PI3K to activate mTORC1 kinase through MAPK ERK signaling. (A) Latent HIV reactivation by 2A127 is abrogated by the MEK inhibitor U0126. Latently infected QUECEL primary Th17 cells were pre-treated or not with U0126 at the concentrations shown for 30 min prior to challenge for 24 h with 1 µM 2A127, 50 nM ingenol or 1 µM prostratin. Proviral reactivation was examined by measuring the dual expression of Tat and Nef using immunofluorescence flow cytometry. Plotted flow data are from three independent experiments conducted with QUECEL primary Th17 cells generated using naïve T cells derived from two donors (D23 and D27). Statistical significance was calculated using a two-tailed Student’s t test. Three asterisks indicate a *p* value < 0.005. Two asterisks indicate a *p* value < 0.01. (B) Scheme showing the regulation of CycT1 translation by diacylglycerol (DAG)-dependent stimulation of RasGRP1 signaling rather than through the stimulation of PI3K. (C) Phosphorylation of p70S6K and rpS6 following PKC agonist stimulation. Healthy donor-derived memory CD4^+^ T cells were challenged for the times shown with 50 nM ingenol, 10 nM bryostatin or with anti-CD3 and anti-CD28 antibodies. (D) 2A127 induces the phosphorylation of p70S6K and rpS6 in memory CD4^+^ T cells. Shown in (C) and (D) are Western blotting data generated using donor D24 cells. (E) Graphs showing the densitometry quantification of the effect of a 4-h 1 µM 2A127 treatment on the phosphorylation of ERK1/2, p70S6K and rpS6 in memory CD4+ T cells. Levels of pT202/Y204 ERK1/2, pT389 p70S6K and pS235/S236 rpS6 were normalized to the total levels of ERK1/2. The plotted average densitometry values are from at least three Western blotting experiments performed using memory CD4^+^ T cells belonging to donors D23, D24 and D27. Error bars denote +/-S.E. of the mean. Statistical significance values shown were calculated using a two-tailed Student’s t test. (F) Ingenol and prostratin stimulate the phosphorylation of p70S6K and rpS6 through MAPK ERK signaling rather than through PI3K. (G) 2A127 stimulates the phosphorylation of p70S6K and rpS6 through MAPK ERK signaling rather than through PI3K. In (F) and (G), healthy memory CD4^+^ T cells from donor D23 were pre-treated or not with 10 µM U0126 (MEK inhibitor) or 5 µM LY294002 (PI3K inhibitor) prior to being challenged with 50 nM ingenol, 10 nM bryostatin, 1 µM 2A127 or stimulated through the TCR for the times shown.

**Figure 7.**
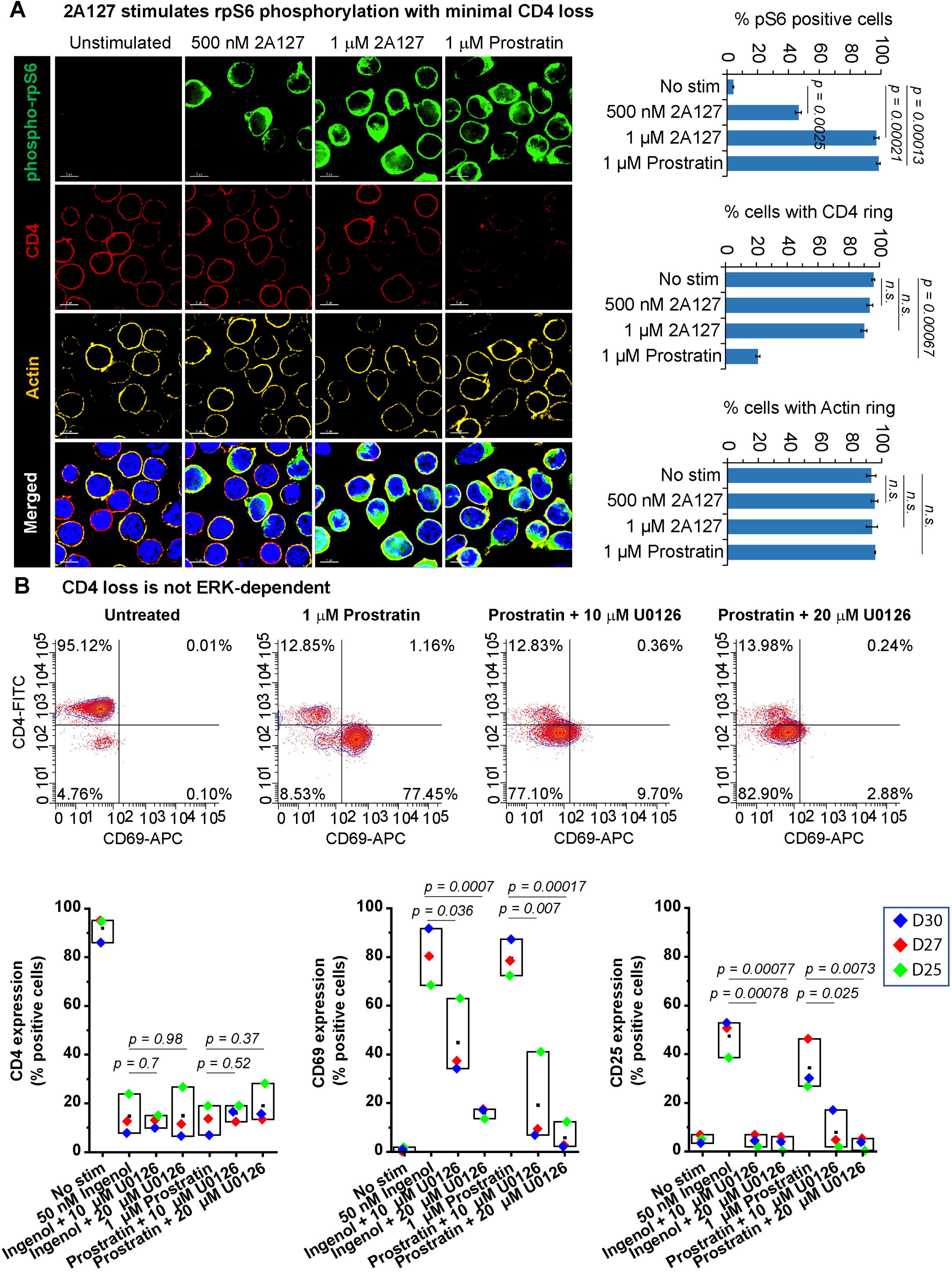
PKC agonist-induced loss of CD4 receptors occurs without altering actin cytoskeletal arrangement and is independent of MEK-ERK1/2 signaling. (A) 2A127 stimulates rpS6 phosphorylation with minimal reduction of surface CD4. Memory CD4^+^ T cells derived from two donors (D29 and D30) were treated or not for 2 h with 500 nM 2A127, 1 µM 2A127 or 1 µM prostratin, followed by co-immunostaining for phospho-rpS6 (pS6), CD4 and actin. At least 100 imaged cells from three independent experiments were used to quantify the percentage of cells expressing phospho-rpS6 (pS6) or maintaining cortical ring patterns of CD4 and actin. Data represent mean ± SEM (*n = 3*). Statistical significance was calculated using a two-tailed Student’s t test and *n.s.* (not statistically significant) indicates a *p* value > 0.05. (B) The MEK inhibitor U0126 blocks the elevation of CD69 expression induced by PKC agonists without rescuing the loss of CD4. *Top panel*, Representative flow experiment demonstrating the effect of MEK inhibition on the dual expression of CD4 and CD69 in memory CD4^+^ T cells that were subjected to 24 h challenge with 1 µM prostratin. *Bottom panel*, Graphed compilation of three independent experiments conducted using memory CD4^+^ T cells from three separate donors (D25, D27, and D30) showing the effect of MEK inhibition on the expression of CD4, CD69 and CD25 following 24 h challenge with 50 nM ingenol or 1 µM prostratin. Statistical significance was calculated using a two-tailed Student’s t test.

Based on the finding that DAG mimetic-induced CycT1 translation is mediated by MEK-ERK1/2, we hypothesized that specifically targeting the C1 domain of RasGRP1 to bypass PKC activation and stimulate P-TEFb via MAPK ERK1/2 could be an optimal strategy for reversing HIV latency. Small-molecule DAG-lactones are a unique class of synthetic DAG mimetics. Their core lactone ring structure, formed by cyclizing the glycerol backbone of DAG, provides stability and high affinity for binding to C1 domain-containing proteins. Unlike complex natural DAG analogs, DAG-lactones can be more easily modified at their sn-1 and sn-2 positions to facilitate the study of C1 domain binding selectivity ^46, 47^. We previously reported that discrimination between the PKC and RasGRP1 C1 domains can be achieved by replacing the alkadiene chain with an indole ring at the sn-2 position of the parent DAG-lactone AJH836. This substitution generates a new series of DAG-indololactones that exhibit 5- to 64-fold in vitro binding selectivity for RasGRP1 (**Fig. 3A and Supplementary Table 1**) ^48, 49^.

We first tested whether these DAG-indololactones reactivate latent HIV in Jurkat cell line models while evaluating their effects on cell viability across a concentration gradient (10 nM to 10 µM). Jurkat T cells constantly express P-TEFb, which is mainly regulated by sequestration into 7SK snRNP ^50^. Measurable latent HIV reactivation in the Jurkat 3C9 clone was observed at a starting concentration of 500 nM for all four tested DAG-lactones (**Supplementary Figs. 8A and 8B**). Latency reversal by DAG-lactones was abolished by GGG to CTC mutations of both NF-κB/NFAT binding elements at the proviral LTR in the Jurkat 2B5 clone (**Supplementary Fig. 8C**). Since the first NF-κB/NFAT element overlaps with a putative HSF1 binding site, the GGG to CTC mutation could also potentially interfere with the recruitment of P-TEFb by HSF1. Recruitment of P-TEFb by HSF1 is thought to be essential for kick-starting HIV transcription elongation to synthesize Tat ^51–53^. Two of the three DAG-indololactones tested – 2A127 and 2C78 – were well-tolerated by Jurkat cells (>90% viability relative to untreated cells) up to a concentration of 1 µM (**Supplementary Fig. 8D**). By contrast, the parent DAG-lactone AJH836 elicited higher levels of Jurkat cell toxicity at both 500 nM and 1 µM (83% and 77% relative cell viability, respectively). Based on these results, we proceeded to test these compounds at a concentration range of 5 nM to 10 µM in primary T cells to evaluate their ability to stimulate P-TEFb expression and reverse HIV latency.

**Figure 8.**
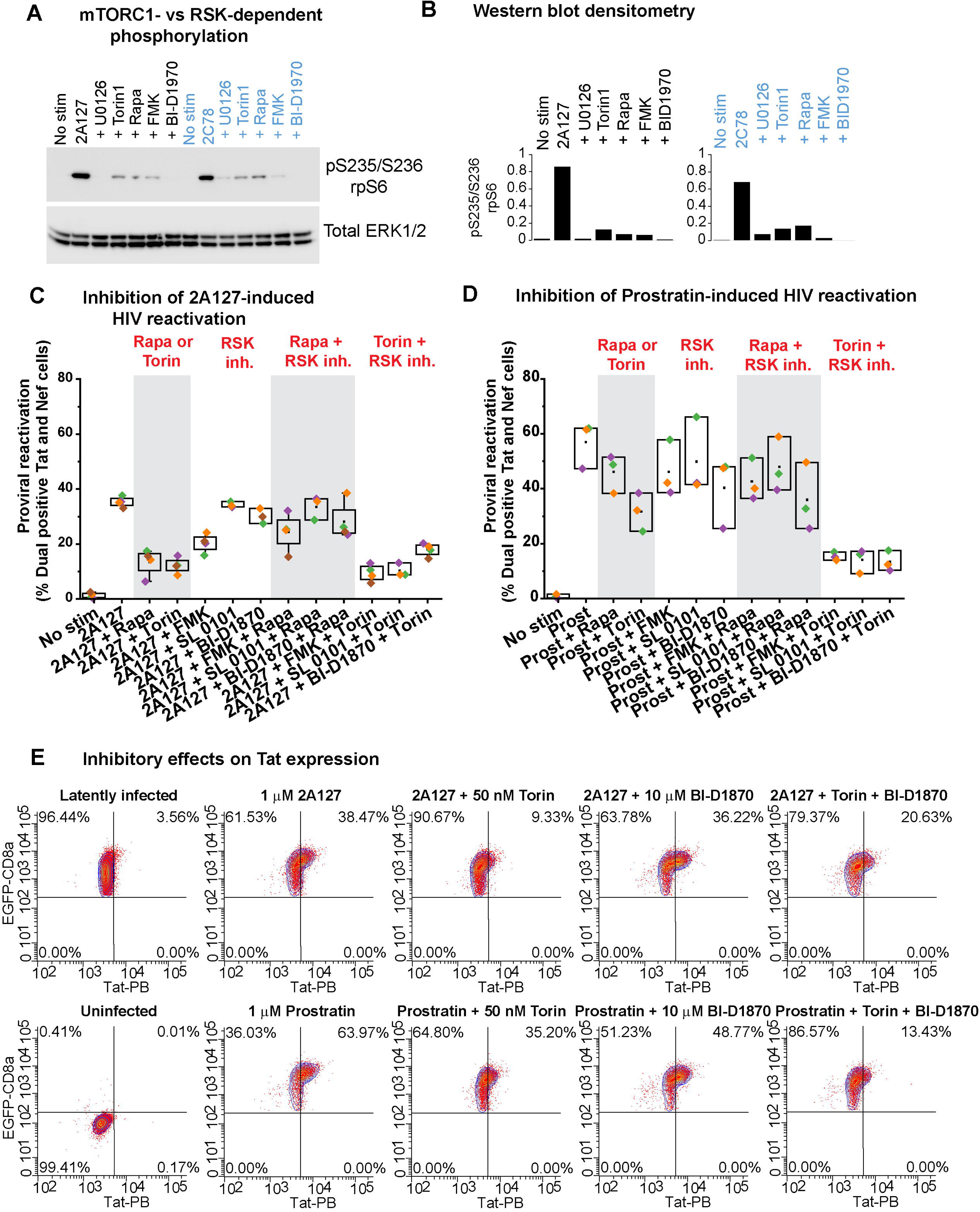
Inhibition of mTORC1 but not RSK effectively suppresses the reactivation of latent HIV by 2A127. (A) Effects of the inhibition of MEK, mTORC1 and RSK on the phosphorylation of rpS6 in QUECEL primary Th17 cells. QUECEL primary Th17 cells were pre-treated or not with the following inhibitors for 30 min: 10 µM U0126, 50 nM Torin1, 500 nM rapamycin, 10 µM FMK and 10 µM BID1970. Thereafter, the cells were treated with 1 µM 2A127 or 1 µM 2C78 for 24 h prior to preparation of whole cell extracts for Western blotting to examine the phosphorylation of rpS6. Total ERK1/2 was also analyzed as a normalization control. (B) Densitometry quantification of the Western blots shown in (A) with levels of pT202/Y204 ERK1/2 and pS235/S236 rpS6 normalized to the total levels of ERK1/2. (C) 2A127-induced HIV reactivation in QUECEL primary Th17 cells is effectively suppressed by rapamycin or Torin1 but not by the RSK inhibitors FMK, SL 0101 or BI-D1870. (D) Prostratin-induced HIV reactivation in QUECEL primary Th17 cells is effectively suppressed by the combined inhibition of mTORC1 and RSK. The results shown in (C) and (D) are from at least three independent experiments conducted with QUECEL primary Th17 cells that were prepared using naïve T cells from two donors (D23 and D27). (E) Representative flow experiment showing the effects of mTOR and RSK inhibition on 2A127- or prostratin-induced latent HIV reactivation in QUECEL primary Th17 cells. In (C), (D) and (E), latently infected cells were pre-treated or not with the following inhibitors individually or in combination for 30 min: 50 nM Torin1, 500 nM rapamycin, 10 µM FMK, 10 µM SL 0101, and 10 µM BID1970. Thereafter, the cells were treated with 1 µM 2A127 or 1 µM prostratin for 24 h.

Healthy donor-derived memory CD4^+^ T cells were treated for 24 h with DAG-lactones, then subjected to P-TEFb immuno-flow analysis to measure the posttranscriptional induction of CycT1 and the biogenesis of transcriptionally active P-TEFb. We also evaluated how these compounds affect T-cell activation by measuring CD25 and CD69 induction. Effective induction of CycT1 and active P-TEFb was observed with three of the four tested DAG-indololactones (2A127, 2C78, and 2C76) as well as the parent DAG-lactone AJH836 at a concentration range of 5-100 nM (**Figs. 3B, 3C, and Supplementary Figs. 9A-D**). Within this concentration range we did not observe any significant increase in the expression of the surface T-cell activation markers CD69 and CD25 (**Figs. 3B, 3C, and Supplementary Figs. 9A-D**). However, at higher concentrations of the DAG-lactones (1 µM and 10 µM), we observed an increase in CD69 and CD25, as well as a more pronounced activation of P-TEFb. Within the tested concentration range, all compounds were well-tolerated, and cell viability at the highest dose (10 µM) was comparable to that seen with the functional doses of the natural PKC agonists (**Supplementary Fig. 10A**). Notably, 1 µM AJH836 induced greater upregulation of CD69 and CD25 than equivalent DAG-indololactones – particularly 2A127, 2C78 and 2D168 – reaching levels similar to natural PKC agonists (**Figs. 3C-D, and Supplementary Fig. 9A-C**). Although the sample sizes were limited, we did not observe any clear differences in how male and female donor memory T cells responded to the DAG-lactones. Therefore, unlike the naturally occurring C1 domain agonists, synthetic DAG-indololactones that preferentially bind to RasGRP1 can trigger CycT1 translation and P-TEFb expression in memory CD4^+^ T cells at doses below those needed for T-cell activation.

### Synthetic RasGRP1 agonists synergize with HDAC inhibitors to reverse HIV latency

A well-characterized ex vivo primary Th17 cell model of latent HIV, the Quiescent Effector Cell Latency (QUECEL) model developed by the Karn laboratory ^54, 55^, was used to characterize the DAG-lactones further. Reactivation of latent HIV in the QUECEL model can be reliably monitored by the dual immunofluorescence detection of Tat and Nef (**Supplementary Figs. 11A and 11B**) ^19^. As reported previously ^7^, 50 nM ingenol was just as effective as TCR co-stimulation at reactivating latent HIV in the QUECEL model (with an average of 62% versus 71% Tat/Nef dual positive cells or 32.4-fold versus 37.3-fold reactivation relative to the unstimulated condition) (**Fig. 4A and Supplementary Fig. 11A**). The lowest tested concentration of the DAG-indololactones 2A127 and 2C78 that resulted in measurable HIV reactivation after 24 h was 250 nM; at this concentration, these compounds elicited a 7.3-fold and 3.4-fold viral induction, respectively (**Fig. 4A**). In comparison, HDAC inhibition with either SAHA (at 500 nM), panobinostat (at 10 nM), or Romidepsin (at 10 nM) caused a 5.5-fold, 13.9-fold, or 12.2-fold reactivation of latent HIV, respectively (**Fig. 4A**). Notably, using the Tat/Nef immunofluorescence assay, we did not observe measurable latency reversal activity in the QUECEL model with any of the other candidate LRA agents tested, namely, the SMAC mimetic AZD5582 (at 100 nM), the bromodomain inhibitor JQ1 (at 1 µM), the BRD4 PROTAC ZXH-3-26 (at 50 nM), and IL-15 (at 50 ng/ml). Moreover, in combined treatments with the DAG-indololactones, none of these four candidate LRAs was able to appreciably enhance the reactivation of latent HIV (**Data not shown**).

We then tested whether combination treatments of a DAG-indololactone and an HDACi can produce synergistic effects in viral reactivation using the Bliss Independence approach to formally evaluate synergy ^56, 57^. All tested drug combinations, including 2A127 or 2C78 with an HDACi (SAHA, panobinostat, or romidepsin), induced effective synergy in viral reactivation in the QUECEL model, with each combination achieving a Bliss score below 1 (**Figs. 4B-D, and Supplementary Figs. 12A and 12B**). Importantly, these combinatorial treatments elicited latent viral reactivation comparable to the potent PKC agonists (ingenol and prostratin) as assessed by dual measurement of Tat and Nef protein expression (**Figs. 4A, 4D, and Supplementary Figs. 12A and 12B**). Cell viability data for these combination treatments are shown in **Supplementary Fig. 10B**.

### DAG-indololactone/HDACi combination reverses latency in clinical reservoirs

Having exhaustively established the synergistic reactivation of latent HIV by combinations of DAG-indololactones and HDAC inhibitors in our ex vivo QUECEL model, we next sought to evaluate the efficacy of the combination treatment in clinical samples. Memory CD4^+^ T cell recovery from phlebotomy-derived PBMCs limited our experimental design to three conditions (no treatment, ingenol positive control, and combination treatment). Thus, the LRA experiments utilizing cells isolated from treated PWH were not designed to test for synergy, but rather to address a specific translational question: Can the combination treatment – at modest, non-toxic concentrations that synergize in the ex vivo QUECEL model while sparing global T-cell activation – achieve sufficient potency to reactivate latent reservoirs in clinical samples? To test this, virus release was measured using an RNA single-copy assay targeting a highly conserved region of the HIV integrase gene ^58^. A 24-hour cellular treatment with 2A127 plus SAHA (at 500 nM each) caused a 10-fold increase in genomic HIV RNA detected in the culture supernatant, compared to a 19-fold increase following treatment with 50 nM ingenol (**Fig. 4E**). Excluding donor D6475, the average fold HIV induction in the four remaining donors was 9.6 after the combination treatment, versus 12.6 with ingenol. We also found that combining 500 nM 2A127 with an HDACi had minimal effects on cell viability in memory CD4^+^ T cells and negligibly induced the expression of CD69 and CD25 (**Figs. 4F**, **5E**, **Supplementary Figs. 10B, 12C and 12D**). Overall, while synergy was formally defined in the QUECEL primary cell model, these PWH-derived studies demonstrate that the DAG-indololactone/HDACi combination achieves potency comparable to a strong PKC agonist in reversing latency in clinical reservoirs, with minimal toxicity and little indication of widespread T-cell activation.

### Bypassing PKC to target RasGRP1 obviates CD4 receptor loss

Among all the LRAs tested in ex vivo latently infected primary T cells and in vivo animal models, PKC-targeting DAG mimetics like those from the phorbol ester family have proven to be the most potent and effective at reversing HIV latency ^59–62^. However, besides causing global T-cell activation, natural PKC agonists have also been reported to lead to the rapid loss of essential immune surface receptors, most notably CD4 in uninfected T cell lines ^22, 23^. Therefore, as part of our investigation into the cellular and viral responses triggered by the DAG-indololactones, we assessed their ability to modulate the expression of CD4 and other surface receptors on primary T cells using flow cytometry. Consistent with previous reports, the natural PKC agonists ingenol (at 10 nM and 50 nM), prostratin (at 500 nM and 1 µM), and bryostatin (at 10 nM), as well as the synthetic DAG-lactone AJH836 (at 1 μM), all caused a significant loss of total CD4 over a 24-hour period, as observed in permeabilized memory CD4^+^ T cells (**Fig. 5A, Supplementary Figs. 13A and 14**). PKC agonist-induced loss of CD4 was rapid and selective, as the expression of CD3, CD28, CD45-Ro, and CCR5 remained largely unaffected (**Figs. 5A, 5B, Supplementary Figs. 13B, 13C and 14**). In contrast, 24 h treatment with the DAG-indololactones 2A127 and 2C78 at a concentration of 500 nM barely caused a shift in CD4 immunofluorescence signal; treatment with these compounds at 1 µM resulted in a modest decrease in overall CD4 expression (**Fig. 5A and Supplementary Fig. 14**). Combination treatments with 2A127 or 2C78 (at 500 nM) plus HDAC inhibitors also had minimal impact on CD4 expression in primary T cells (**Fig. 5C and Supplementary Fig. 13D**), while reactivating latent HIV in QUECEL primary Th17 cells to a level comparable to that observed with PKC agonists or a combination of ingenol plus ionomycin (**Figs. 4A, 4C, 4D, Supplementary Figs. 12A and 15**,). Therefore, we conclude that while potent PKC-targeting DAG mimetics cause a rapid, selective, and significant downregulation of the CD4 receptor in primary T cells, DAG-indololactones or their combination with HDACi can reverse HIV latency with minimal changes in CD4 expression.

### Latency reversal by DAG-indololactones involves stimulation of ribosomal activity through MEK-ERK1/2-mTORC1 signaling

Our demonstration that the PI3K-mTORC2-AKT-mTORC1 pathway acts as a complementary TCR-activated route for promoting P-TEFb expression ^7^, suggested that mTORC1 can participate in P-TEFb biogenesis by facilitating the translation of CycT1. While the activation of mTORC1 kinase primarily by PI3K-mTORC2-AKT signaling has long been linked to the initiation of protein synthesis in most cell types, including T cells ^63, 64^, DAG-mimicking agonists that target the C1 domains of RasGRP1 or PKC enzymes are not known to signal through this pathway. However, activated GTP-bound Ras can bind PI3K at the membrane to positively influence its lipid kinase activity ^65–68^. Using the DAG-indololactone 2A127 as an experimental tool to investigate the signaling mechanism of HIV latency reversal by RasGRP1 agonism, we found that the MEK inhibitor U0126 completely suppressed the reactivation of latent HIV in the QUECEL model induced by 1 μM 2A127, as measured by dual immunofluorescence detection of Tat and Nef expression (**Fig. 6A and Supplementary Fig. 16A**). U0126 also blocked the reactivation of latent HIV induced by 1 μM prostratin. Similarly, induction of active P-TEFb by either 2A127 or prostratin was substantially suppressed by U0126 treatment (**Supplementary Figs. 16B and 16C**). Overall, these results show that C1 domain agonists, especially those that preferentially target RasGRP1, promote P-TEFb production and the subsequent synthesis of viral proteins in reactivated primary T cells through MEK-ERK1/2 signaling.

How could the RasGRP1-Ras-Raf-MEK-ERK1/2 pathway trigger the translation of CycT1 to reverse HIV latency? The kinase activity of mTORC1 is considered essential for the rapid initiation of protein synthesis through the sequential phosphorylation and activation of components of the translation machinery, which include the ribosomal S6 kinase (S6K), its downstream substrate ribosomal protein S6 (rpS6), as well as initiation and elongation factors^24^. We therefore tested whether ERK stimulation by C1 domain agonists (PKC agonists or synthetic DAG-lactones) can activate mTORC1, as indicated by phosphorylation of both S6K and rpS6 (**Fig. 6B**). Treating healthy donor-derived memory CD4^+^ T cells with 50 nM ingenol or 10 nM bryostatin caused a rapid (within 2 h) phosphorylation of p70S6K and rpS6, which coincided with ERK1/2 phosphorylation and was comparable to levels seen after TCR co-stimulation (**Fig. 6C and Supplementary Figs. 17A-D**). While ingenol caused a sustained phosphorylation of p70S6K and rpS6 throughout the 24 h treatment, bryostatin’s effect was temporary and matched the duration of ERK1/2 phosphorylation (**Fig. 6C and Supplementary Fig. 17A**). Western blotting of memory T cells treated with 2A127 at concentrations of 10 nM, 100 nM, and 1 μM also showed phosphorylation of p70S6K and rpS6 at 1 μM, coinciding with phosphorylation of ERK1/2 (**Figs. 6D and 6E**). Fluorescence microscopy revealed that a 2 h treatment of memory T cells with 500 nM 2A127 induced rpS6 phosphorylation in 46% of the cells examined without affecting the surface expression of CD4, unlike treatment with 1 µM prostratin (**Figure 7A**). Notably, the drastic CD4 downregulation caused by ingenol or prostratin could not be rescued by inhibition of MEK (**Figure 7B**), suggesting that this process does not proceed through RasGRP1-MEK-ERK signaling. Furthermore, PKC agonist-induced CD4 loss occurred without altering the actin cytoskeleton, bolstering the evidence for the selectivity of this mechanism **(Figure 7A**).

Inhibitor studies were used to confirm that the stimulation of p70S6K and rpS6 phosphorylation by C1 domain agonists was not due to a crosstalk between RasGRP1-Ras and PI3K signaling and was instead mediated by MEK-ERK1/2. p70S6K and rpS6 phosphorylation following cellular treatment with ingenol, bryostatin or 2A127 were effectively repressed by MEK inhibition with U0126 (**Figs. 6F, 6G, Supplementary Figs. 17E and 17F**). In contrast, inhibiting PI3K with LY294002 did not alter the levels of phosphorylated p70S6K and rpS6 produced in response to either of these agents (**Figs. 6F, 6G, Supplementary Figs. 17E, and 17F**). U0126 and LY294002 each blocked TCR-induced phosphorylation of p70S6K and partially inhibited the formation of phospho-rpS6 (**Fig. 6G and Supplementary Fig. 17F**), consistent with the stimulation of ribosomal activity by both the PI3K-mTORC2-AKT and MEK-ERK1/2 signaling pathways during TCR co-stimulation. These immunoblotting results therefore confirm that the DAG-mimicking C1 domain agonists signal through ERK1/2 rather than PI3K to elicit the phosphorylation of p70S6K and rpS6.

ERK1/2 has been reported to positively regulate mTORC1 activity in the context of cellular and tissue models of carcinogenesis by directly phosphorylating and inactivating the TSC1/TSC2 GTPase-activating protein (GAP) complex, which is crucial for suppressing mTORC1 signaling activation by the GTPase Rheb (**Fig. 6B**) ^69^. Additionally, ERK phosphorylates RSK (p90S6K), which, like the mTORC1-activated p70S6K, can phosphorylate rpS6 ^70–72^. Upon activation by ERK, RSK also phosphorylates TSC1/TSC2, leading to increased mTORC1 signaling to p70S6K ^73^. We therefore tested whether ERK-induced phosphorylation of rpS6 and ERK-mediated reactivation of latent HIV in primary T cells are dependent on mTORC1, RSK, or both kinases.

Inhibition of mTORC1 activity with Torin1 or rapamycin effectively suppressed the phosphorylation of rpS6 induced in primary T cells by the DAG-lactones (2A127, 2C78, or AJH836) or ingenol to a comparable extent as U0126 treatment (**Figs. 8A, 8B, and Supplementary Figs. 18A-D**). Similarly, inhibition of RSK with FMK or BID1970 also effectively suppressed rpS6 phosphorylation induced by either 2A127, 2C78, AJH836 or ingenol (**Figs. 8A, 7B, and Supplementary Figs. 18A-D**). In contrast to these results, we observed different effects of the mTORC1 and RSK inhibitors on HIV latency reversal in the QUECEL model depending on the C1 domain agonist used as a stimulus. Reactivation of latent HIV by the DAG-indololactones 2A127 and 2C78 was effectively blocked by the mTORC1 inhibitors and not as well suppressed by RSK inhibitors (**Figs. 8C, 8E, Supplementary Figs. 19A and 19B**). Moreover, combined inhibition of mTORC1 and RSK did not enhance viral suppression beyond that observed with the individual mTORC1 inhibitors (**Fig. 8C, Supplementary Figs. 19B and 19C**). For viral reactivation triggered by ingenol, prostratin, or the DAG-lactone AJH836, inhibiting either mTORC1 or RSK was ineffective or only slightly reduced viral reactivation (**Figs. 8D, 8E and Supplementary Figs. 19A, 19D and 19E**). By contrast, combined inhibition of RSK and mTORC1 caused a blockade in viral reactivation triggered by ingenol, or prostratin that was similar to or nearly as effective as the blockade seen with the MEK inhibitor U0126.

We also examined the effects of mTORC1 and RSK inhibition on CD69 and CD25 expression in memory CD4^+^ T cells and in quiescent QUECEL primary Th17 cells. While the extent of the posttranscriptional expression of CD69 induced by 24 h challenge with ingenol or prostratin was comparable in both cell types, CD25 elevation induced by either agent was observed to be substantially lower in memory CD4^+^ T cells (**Fig. 9A**). Surprisingly, we found that inhibition of mTORC1, RSK, or even their combined inhibition was ineffective at suppressing CD69 induction in both cell types by any of the C1 domain agonists tested (1 μM prostratin, 50 nM ingenol, or 1 μM AJH836) (**Figs. 9B-D and Supplementary Figs. 20A and 20B**). CD25 expression in both cell types was also refractory to inhibition of mTORC1, RSK or their combined inhibition with the odd exception that BI-D1870, which is known to inhibit all the four known isoforms of RSK, effectively blocked CD25 elevation in memory T cells but modestly inhibited its expression in primary Th17 cells (**Figs. 9E, 9F, Supplementary Figs. 20C and 20D**). In contrast to observations made with the mTORC1 and RSK inhibitors, elevation of CD69 and CD25 by any of the C1 domain agonists tested was found to be sensitive to the inhibition of MEK in both cell types (**Figs. 9B-F and Supplementary Figs. 20A-D**). The impact of these agonist and inhibitor treatments on the viability of memory CD4^+^ T cells and primary QUECEL Th17 cells are shown in **Supplementary Fig. 21**.

**Figure 9.**
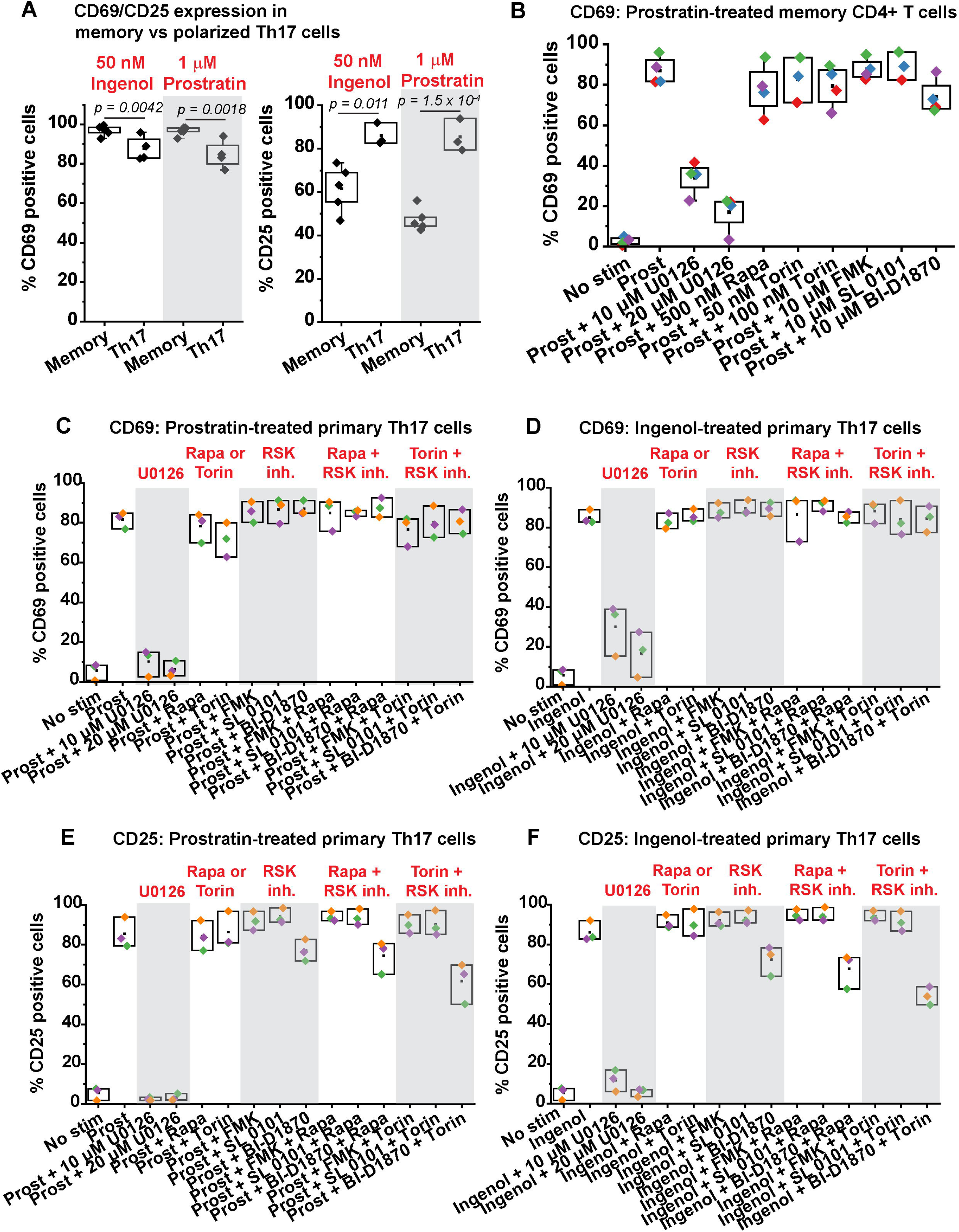
PKC agonist-induced elevation of CD69 and CD25 is refractory to the inhibition of mTOR in both memory CD4^+^ and QUECEL primary Th17 cells. (A) Comparison between memory CD4^+^ and in vitro polarized QUECEL primary Th17 cells in CD69 and CD25 expression induced by 24 h treatment with 50 nM ingenol or 1 μM prostratin. Statistical significance was calculated using a two-tailed Student’s t test. (B-F) Effect of individual or combination inhibitor treatments on the inducible expression of CD69 or CD25 in memory CD4^+^ or primary Th17 cells. Cells were pre-treated with inhibitors for 30 min prior to a 24-h challenge with 50 nM ingenol or 1 µM prostratin. Inhibitor concentrations were as follows: 10 µM or 20 µM U0126, 50 nM Torin1, 500 nM Rapamycin, 10 µM FMK, 10 µM SL 0101 and 10 µM BID1970. Fixed and permeabilized cells were then subjected to immunofluorescent staining for CD69 and CD25. The results shown in (B) from four independent experiments conducted with memory CD4^+^ T cells from four donors (D9, D10, D24 and D25). The results shown in (C-F) are from three independent experiments conducted with QUECEL primary Th17 cells that were prepared using naïve T cells from two donors (D23 and D27).

Overall, these findings demonstrate that the posttranscriptional induction of CD69 by C1 domain agonists in primary T cells is sensitive to MEK-mediated blockade of ERK1/2, yet resistant to the inhibition of mTORC1, RSK, or both kinases. Similarly, agonist-induced CD25 expression is mediated by MEK-ERK signaling but does not involve mTORC1 activity. Conversely, the downregulation of CD4 induced by C1 domain agonists is independent of MEK-ERK signaling. Furthermore, while agonist-induced rpS6 phosphorylation is significantly blunted by inhibiting either mTORC1 or RSK, these pathways affect HIV latency reversal differently depending on the specific agonist used. Specifically, HIV latency reversal by DAG-indololactones in the QUECEL model is hindered by mTORC1 inhibition, but not by RSK inhibitors. Therefore, while the RasGRP1-MEK-ERK1/2 pathway is essential for mediating latent HIV reactivation by stimulating ribosomal activity in response to various C1 domain agonists, DAG-indololactones rely exclusively on mTORC1 to promote proviral gene expression.

## DISCUSSION

### Induction of P-TEFb by DAG-indololactones leads to HIV latency reversal

HIV latency in infected primary T cells is enforced by the phenotypic transition to a quiescent memory state which enables an intracellular environment that is non-permissive to host-initiated viral transcription ^74^. Emergence of HIV from latency requires viral promoter activation by host transcription initiation factors, leading to relief of restrictive chromatin, and stimulation of transcription elongation by P-TEFb. Quiescent primary T cells possess a nuclear absence of host initiation factors and are deficient in P-TEFb expression due to translational repression of its CycT1 subunit ^8, 12^. In contrast, P-TEFb is constitutively expressed in actively dividing T cells where it is primarily sequestered and inhibited by 7SK snRNP ^31, 50^. Using both primary cell models and cells derived from PWH we have confirmed the essential role of P-TEFb in the reversal of HIV latency and defined the cellular signaling pathways needed to induce P-TEFb in resting memory T cells.

PKC agonists are the most potent single-molecule agents capable of reversing HIV latency in ex vivo primary T cells ^59–62^, partly because, unlike all other tested small-molecule compounds, they can stimulate viral transcription initiation as well as potently induce the expression of P-TEFb. However, their usefulness as LRAs is limited by adverse side effects, which include widespread coagulopathy, triggering global T-cell activation and inducing a significant loss of CD4 receptor expression ^22, 23, 75–77^. Synthetic DAC mimics such as bryostatin 1 analogs (bryologs), prostratin analogs, and DAG-lactones have already shown promise as effective LRAs in cell line and primary cell models because they are designed to target the C1 domains of PKC enzymes with high affinity ^78–83^. It is therefore likely that their latency-reversal effects are also triggered through PKC-dependent canonical NF-κB signaling and other pathways known to be linked to overall T-cell activation. Interestingly, PKC signaling also promotes the activation and nuclear translocation of HSF1 ^84–86^, a transcription factor whose putative binding sequence overlaps with the NF-κB/NFAT element at the HIV promoter ^51^. HSF1 is thought to be essential for driving early, Tat-independent viral transcription by recruiting P-TEFb ^52, 53^. Thus, identification of PKC agonists that retain the potency to stimulate viral transcription via P-TEFb and possibly HSF1, while bypassing canonical NF-κB activation, is a critical objective. Recent in vivo studies show that a synthetic, PKC-ε-selective agonist and an African plant extract are well-tolerated at doses that successfully reverse HIV latency ^77, 87^. These findings revitalize confidence in the utilization of PKC agonists in HIV cure studies.

Building on our prior findings that natural PKC agonists activate P-TEFb expression to reactivate latent HIV via a PKC-independent RasGRP1-Ras-Raf-MEK-ERK1/2 pathway ^7^, we tested a series of synthetic DAG mimics with known varying affinities for PKC and RasGRP1 to assess their effectiveness as non-toxic LRAs. Discrimination between the PKC and RasGRP1 C1 domains can be achieved by adding an indole ring structure at the sn-2 position of the parent DAG-lactone AJH836, which changes the compound from a strong PKC agonist to a series of DAG-indololactones that show 5- to 64-fold in vitro binding preference for RasGRP1 ^48,49^. Finding synthetic RasGRP1-selective C1 domain agonists that can effectively promote P- TEFb expression and activity in primary T cells is a promising LRA approach that could avoid undesirable cellular and systemic inflammatory responses. We showed that synthetic DAG-indololactones induce active P-TEFb (defined as the coordinated posttranscriptional expression of CycT1 and phosphorylation of CDK9 at Ser175), at a concentration range that is well-tolerated in memory CD4^+^ T cells with minimal increases in the T-cell activation markers CD69 and CD25.

To the best of our knowledge, there are no available small molecule compounds specifically intended to target the biogenesis of P-TEFb or stimulation of its kinase activity in primary T cells. Our current study sets a precedent in revealing a signaling mechanism for inducing P-TEFb synthesis in primary T cells and identifying RasGRP1 as a key molecular target for initiating this process. Crucially by demonstrating a clear mechanistic separation between P-TEFb production (with subsequent HIV latency reversal) and unwanted cellular side effects (CD4 downregulation and global T-cell activation), we establish the RasGRP1-ERK1/2-mTORC1 axis as a precise, non-toxic pathway for P-TEFb activation. These insights directly support the utility of DAG-indololactones in dual-targeted latency-reversing strategies for HIV cure efforts.

### Synergy between DAG-indololactones and HDAC inhibitors in HIV latency reversal

Combination treatments of a DAG-indololactone (2A127 or 2C78) and an HDACi (SAHA, panobinostat, or romidepsin) showed effective synergy in reactivating latent HIV in the QUECEL primary Th17 cell model, with reactivation levels comparable to those seen with ingenol treatment. While natural (ingenol, prostratin, and bryostatin) and synthetic (AJH836) PKC agonists caused rapid, selective, and significant downregulation of the CD4 receptor on memory CD4^+^ T cells, the representative DAG-indololactones 2A127 and 2C78, alone or combined with HDACi, minimally affected CD4 expression.

By measuring virus release using a single-copy RNA assay targeting the HIV integrase region, we further demonstrated that the combination of 2A127 and SAHA could reverse latency in memory CD4^+^ T cells from five treated PWH. Inhibition of HDAC activity is well known to promote transcription initiation by reversing epigenetic chromatin restrictions on promoter regions, thus enabling the recruitment of RNAP II to the promoter without the need for transcription initiation factors. The LRA candidates JQ1, BRD4 PROTAC ZXH-3-26, SMAC mimetic AZD5582, and IL-15 failed to reverse HIV latency in the QUECEL primary cell model. Furthermore, unlike HDAC inhibitors, none of these agents showed significant synergistic reactivation in combination with DAG-indololactones. These findings are consistent with previous reports showing that when used alone, JQ1, ZXH-3-26, AZD5582 or IL-15 elicit weak, ineffective, or variable latency reversal responses in a monoculture of primary T cells ^60, 88–90^. For instance, AZD5582, a potent activator of the non-canonical NF-κB pathway ^91^, robustly induces latency reversal in cell lines and shows measurable responses in in vivo animal models ^92^. However, recent ex vivo CRISPR and transcriptomic evaluations show that AZD5582 alone fails to produce full-length, viral particle-producing transcripts in CD4^+^ T cells from ART-suppressed individuals ^93^. Despite being capable of potently reactivating latent HIV in cell line models by inducing the release of P-TEFb from the inhibitory 7SK snRNP complex or by degrading BRD4 to eliminate the competition for P-TEFb ^88, 94^, JQ1 and ZXH-3-26 exhibit poor LRA activity in primary T cells ^60, 88^, likely because they do not stimulate the biogenesis of P-TEFb. Results from our combination treatments also suggest that HDAC inhibitors may be substantially more potent at driving HIV transcription initiation in primary T cells than the activation of the JAK/STAT5 pathway with IL-15 or activation of non-canonical NF-κB.

Although P-TEFb mobilization is required for HIV to emerge from latency, it is not sufficient for efficient reactivation. At the concentrations used in the current study for combination treatments (250 nM and 500 nM), the DAG-indololactones would be ineffective initiators of HIV transcription because, unlike PKC agonists, they are unable to activate transcription initiation factors such as canonical NF-κB. Consequently, reversing latency with these compounds necessitates a two-pronged strategy: combining a RasGRP1 agonist to induce posttranscriptional P-TEFb expression with a HDAC inhibitor to promote RNAP II recruitment to the HIV promoter. Together, these agents provide the synergy needed to reverse HIV latency while minimizing global T-cell activation.

### DAG-indololactones do not downregulate CD4 on primary T cells

Loss of CD4 from the cell surface following PKC agonist treatment was found to be sustained over a 24 h period and is unlikely to be transient because it involves internalization and degradation of the receptor. Although we did not test the effect of withdrawing the agonist, the results of others suggest that internalized CD4 is usually routed for lysosomal degradation ^95^; therefore, receptor recovery would require de novo synthesis . Other surface receptors examined namely CD3, CD28, CD45-Ro and CCR5 were not impacted by PKC agonist treatment, clearly implying a selective CD4 downregulatory effect. Internalization and subsequent degradation of CD4 after treatment with PKC-targeting, C1 domain-binding DAG mimetics depend on the transient phosphorylation of its cytoplasmic domain. CD4 phosphorylation, likely by activated PKC, causes the receptor to dissociate from p56LCK and associate with clathrin-coated pits ^23, 96^. The finding that the DAG-indololactones 2A127 and 2C78 cause minimal CD4 loss compared to their parent DAG-lactone AJH836, a potent PKC agonist, clearly indicates that selectively targeting RasGRP1 over PKC effectively prevents the mechanism that leads to CD4 downregulation. Moreover, we demonstrate that the PKC agonist-induced loss of CD4 is independent of RasGRP1-MEK-ERK1/2 signaling since the MEK inhibitor U0126 is unable to prevent this process. We consider the CD4 downregulation to be an undesirable side effect because immune activation of both naive and memory CD4^+^ T cells during antigen presentation critically depends on the recognition and binding of CD4 to the antigen-presenting MHC Class II molecule ^97, 98^. Therefore, DAG mimetics that can bypass PKC activation to signal through RasGRP1 selectively are the more suitable candidate compounds for safely reversing HIV latency.

### Regulation of CycT1 synthesis in primary T cells

Unstimulated CD4^+^ T cells are mainly quiescent but are poised to rapidly transition to an activated state by accumulating a large number of idling ribosomes and keeping a pool of approximately 250 essential mRNA species in a translationally repressed state ^25, 99^. Whereas prior studies have implicated both microRNA-mediated translational inhibition and proteolytic turnover in restricting P-TEFb availability during quiescence ^12, 13, 25^, our kinetic and pharmacological dissection establishes that relief of translational repression – rather than de novo transcription or constitutive proteasomal degradation – is the primary rate-limiting checkpoint governing CycT1 expression in primary T cells. Specifically, while TCR co-stimulation produced only minimal changes in CycT1 transcript abundance (as examined by RNA FISH and consistent RNA-seq measurements), it drove a rapid, massive surge in CycT1 protein synthesis that was completely abolished by cycloheximide (CHX) yet could not be recapitulated by proteasomal inhibition with MG132 alone. Moreover, CHX chase assays demonstrated that once synthesized upon activation, CycT1 is stable, in sharp contrast to the rapidly turned-over cell-cycle regulator Cyclin D3 (CycD3). By enforcing this stringent posttranscriptional block on P-TEFb biogenesis, resting memory CD4^+^ T cells maintain an intracellular environment uniquely conducive to preserving transcriptionally latent HIV proviruses in ART-treated PWH.

Comparing CycT1 with CD69 further illuminated the signaling and spatial logic of posttranscriptional control in primary T cells. By decoupling transcription from translation using the covalent CDK7 inhibitor THZ1 and actinomycin D, we confirmed that whereas inducible CD25 expression strictly requires de novo transcription, rapid CD69 and CycT1 protein induction following TCR co-stimulation or C1-domain agonist challenge relies on the mobilization of pre-existing mRNA pools. Consistent with this observation, the CycT1 promoter is known to be TATA-less, constitutively active and modestly responsive to cellular activating stimuli such as PKC agonists ^100, 101^. Despite its highly inducible transcriptional expression, CD69 mRNA has also been identified as one of the most robust targets of the AU-rich element-binding protein ZFP36 in a HITS-CLIP (high-throughput sequencing of RNA isolated by crosslinking immunoprecipitation) analysis in primary T cells ^35^. ZFP36 binds the CD69 3’ untranslated region to restrain its expression, and TCR-induced phosphorylation or genetic depletion of ZFP36 rapidly releases these transcripts for immediate translation ^35, 102, 103^. Notably, flavopiridol (FVP) at 100 nM selectively abrogated P-TEFb-dependent HIV-1 reactivation without impairing CD25 or CD69 induction, underscoring both pathway specificity and the heightened reliance of latent proviruses on threshold P-TEFb activity. High-resolution RNA FISH and immunofluorescence imaging further revealed that translationally silenced CycT1 and CD69 transcripts occupy distinct cytoplasmic territories in resting memory T cells. While a prominent fraction of CD69 mRNA resides within EDC4-positive P bodies, CycT1 transcripts are excluded from both P bodies and phospho-rpS6–marked active ribosomes. Pre-existing CycT1 mRNA may therefore be sequestered within alternative phase-separated RNP assemblies, such as stress granules, which harbor translationally silent transcripts alongside stalled initiation factors and poised ribosomes ^14^.

Following its posttranscriptional induction, CycT1 protein immediately establishes a nuclear localization. High resolution imaging revealed that P-TEFb, 7SK snRNA, and the hyperphosphorylated form of RNAP II (pSer2 RNAP II CTD) share the same subnuclear space in activated primary T cells. We have previously observed a similar doughnut-shaped strong nuclear colocalization for HEXIM1/7SK and Tat/7SK following T-cell activation ^7, 104^. These findings suggest that the 7SK snRNP inhibitory complex localizes to the same nuclear condensate as actively transcribing genes, as marked by pSer2 RNAP II CTD. This close placement likely allows for the rapid and convenient transfer of P-TEFb to transcriptionally poised genes.

### RasGRP1-mediated ERK signaling pathway activates P-TEFb through mTORC1

Our previous RNA-seq analysis has shown that RasGRP1 is the overwhelmingly predominant DAG-dependent RasGRP isoform in both resting and activated memory CD4^+^ T cells ^7^, implying that it is the primary initiator of Ras-Raf-MEK-ERK1/2 signaling. Stimulation of RasGRP1 by DAG in T cells is necessary for the allosteric activation of membrane-bound SOS, a second GDP-GTP exchange factor for Ras ^105^. Guanine nucleotide exchange activity of SOS is triggered by the initial activation of Ras by RasGRP1 through a positive feedback mechanism that involves the allosteric binding of GTP-bound Ras to SOS. Ras activation by RasGRP1 and SOS causes a conformational change in Ras, enabling the stable formation of Ras:Raf complexes at the membrane that eventually leads to the activation of MEK and its substrates, the MAP kinase isoforms ERK1 and ERK2 ^106^. When targeting the RasGRP1-Ras-Raf-MEK-ERK1/2 pathway for HIV latency reversal, it is important to note that although dysregulated Ras signaling has been linked to many types of human cancer, Ras’s oncogenic activity is solely attributed to genetic mutations that transform it from a proto-oncogene into an entity with constitutive activity ^107^. DAG or DAG-mimicking C1 domain agonists that reversibly interact with RasGRP1 cannot induce oncogenic activity because they trigger only transient Ras signaling that remains under physiological control.

In our mechanistic study of how CycT1 translation is regulated in memory CD4^+^ T cells, C1 domain agonists, including DAG-indololactones, were used as experimental tools to explore the cellular mechanism by which RasGRP1 signaling can break the translational repression of CycT1 and reverse HIV latency. In these experiments, we also performed a parallel analysis of the expression of the T cell activation markers CD69 and CD25. ERK has been shown to stimulate mRNA translation by inactivating the TSC1/TSC2 GTPase activating protein complex, which then leads to mTORC1 activation ^73^. In addition, ERK may also effect protein synthesis through two other mechanisms: a) by activating RSK which, like the mTORC-activated S6K, can phosphorylate rpS6 ^70, 71^; and b) by phosphorylating MNK which in turn phosphorylates the translation initiation factor eIF4E ^108^. We initially observed that MEK inhibition abrogated the reactivation of latent HIV by DAG-indololactones in the QUECEL primary cell model and substantially suppressed P-TEFb induction by these agents. We also found that the ability of DAG-indololactones to reverse HIV latency in the QUECEL primary cell model is effectively suppressed by inhibition of mTORC1, but not by RSK inhibitors. Conversely, although both the induction of CycT1 and HIV latency reversal following treatment with PKC agonists were effectively blocked by MEK inhibition, a combined inhibition of mTORC1 and RSK was necessary to achieve a blockade in viral reactivation similar to that seen with the MEK inhibitor. Based on these findings, we conclude that activation of the RasGRP1-MEK-ERK1/2 pathway by a wide range of C1 domain agonists is crucial for inducing latent HIV reactivation in primary T cells. This occurs through the stimulation of ribosomal activity in a manner dependent on mTORC1 and RSK. Specifically, our studies also show that DAG-indololactones signal exclusively through mTORC1 to promote proviral gene expression. Interestingly, inhibiting mTORC1, RSK, or even combining their inhibition was found to be largely ineffective at reducing the expression of CD69 and CD25 induced by ingenol, prostratin or the synthetic DAG-lactone AJH836. Since the C1 domain agonist-driven expression of these activation markers appears to be ERK-dependent, it is likely that their expression might be mediated by ERK-phosphorylated MNK, which was not examined in this study. These results uncover for the first time the mRNA translation mechanisms that RasGRP1-mediated ERK signaling use to stimulate P-TEFb biogenesis and facilitate the emergence of HIV from latency in primary T cells.

### Signaling crosstalk and off-target effects

Beyond directly activating RasGRP1 via its C1 domain, PKC agonists likely modulate its activity through PKC-dependent phosphorylation at Thr184 ^109^. Exploring this signaling crosstalk and its downstream impact on the MEK-ERK-mTORC1 axis will be an important next step once the phospho-Thr184 RasGRP1 antibody becomes available. Off-target signaling effects of PKC agonists and DAG-indololactones may arise from binding to other diacylglycerol-activated, C1 domain-containing proteins. These include protein kinase D, diacylglycerol kinases, α and β chimaerins, and Munc13 proteins ^110^. To further establish RasGRP1 as a key driver of P-TEFb biogenesis and HIV latency reversal, we attempted to knock down the expression of this guanine nucleotide exchange factor using a set of validated siRNAs. This approach failed, likely because the short in vitro lifespan of unstimulated memory T cells is insufficient to deplete constitutively expressed RasGRP1. Because PROTAC chemical degraders can rapidly clear stable proteins in primary T cells – as recently demonstrated with BRD4 ^88^ – we intend to leverage this technology by designing a RasGRP1-targeting PROTAC that covalently links a DAG-indololactone to an E3 ligase recognition moiety.

## METHODS

### Ethics statement

De-identified, cryopreserved peripheral blood mononuclear cells (PBMCs) from individuals with HIV on antiretroviral therapy were obtained from the Rustbelt CFAR Clinical Core repository. The study protocol (UH# 12-11-33) was approved by the Institutional Review Board of University Hospitals Cleveland Medical Center. All participants provided written informed consent, and all specimens and data were coded to maintain confidentiality.

### Ex vivo generation of uninfected or latently infected primary Th17 cells

The optimized procedure for preparing quiescent, latently infected primary Th17 cells, known as the Quiescent Effector Cell Latency (QUECEL) model has been previously described in extensive detail ^19, 54, 55^. De-identified, cryopreserved and healthy donor-derived peripheral blood mononuclear cells (PBMCs) were commercially obtained from AllCells and StemCell Technologies (**Supplementary Table 2**). Briefly, naïve CD4^+^ T cells were isolated from these PBMCs by negative bead selection using the EasySep Naïve CD4^+^ T-cell isolation kit (StemCell) and activated with 2 μg/ml Concanavalin A (Con A) while being polarized to the Th17 effector phenotype using the following cocktail of cytokines and antibodies: IL-1β (10 ng/ml), IL-2 (60 IU/ml), IL-6 (30 ng/ml), IL-23 (50 ng/ml), TGF-β (5 ng/ml), anti-IL-4 (500 ng/ml), and anti-IFN-γ (10 ng/ml). Following a 7-day period of Con A-induced T-cell expansion, polarized Th17 cells were infected with a VSVG-pseudotyped pHR’-CD8a-EGFP-Nef+ single-round HIV construct, then subjected to a positive selection of acutely infected cells using the EasySep Mouse CD8a Positive Selection Kit II (StemCell) at Day 4 post-infection. Viral latency was established by enforcement of a quiescent phenotype through a 2-week culture of the positively selected cells in medium supplemented with IL-8 (50 ng/ml), IL-10 (10 ng/ml), TGF-β (10 ng/ml) and reduced concentrations of IL-2 (15 IU/ml) and IL-23 (31.25 ng/ml). Achievement of quiescence in both the uninfected and infected cells were examined in the context of T-cell activation by monitoring the expression of CD25, CD69, and P-TEFb using immunofluorescence flow cytometry. Entry of HIV into latency and its reactivation were also measured by immunofluorescence flow cytometry analysis of Tat and Nef.

### Isolation of memory CD4^+^ T cells

Approximately 10 to 15 million memory CD4^+^ T cells were routinely isolated by negative bead selection from 100 million de-identified, cryopreserved and healthy donor-derived PBMCs using the EasySep memory CD4^+^ T-cell enrichment kit (StemCell). Designation of the donor PBMCs used in the current study is shown in **Supplementary Table 2**. Isolated cells were allowed to recover overnight in complete RPMI medium lacking IL-2 prior to treating them with inhibitors and/or stimuli described in the current study. Purity and quiescent status of the memory CD4^+^ T cells were confirmed by immunofluorescence flow cytometry analysis of CD45-RO, CD69, CD25 and P-TEFb expression.

### Primary cell culture and compounds

Polarized primary Th17 or freshly isolated memory CD4^+^ T cells were cultured in complete RPMI medium containing 10% fetal bovine serum, 100 IU/ml penicillin, 100 µg/ml streptomycin, and 25 mM Hepes, pH 7.2. To examine P-TEFb or latent HIV reactivation, cells were equally divided into 1-ml 24-well cultures at 3.0 X 10^5^/ml then challenged with the agonists or stimuli under investigation for 24 h. To investigate the signaling pathways involved, cells were pre-treated for 30 min with the experimental inhibitors prior to the 24-h challenge with the agonists or stimuli under investigation. The effect of these treatments on cell viability was assessed by either propidium iodide staining or eFluor 450 fixable viability dye staining. A series of structurally defined DAG-lactones bearing a 1-methyl-1H-indole at the sn-2 position were chemically synthesized using the parent DAG-lactone compound AJH836 as previously described ^48, 49^.

### Bliss Independence for evaluation of LRA synergy

Combination treatments of a DAG indololactone and an HDACi were evaluated for drug synergy using the Bliss Independence approach with all of the tested compounds satisfying the following Bliss criteria: 1) There is existing knowledge on the drugs’ mechanisms of action; 2) DAG-indololactones and HDAC inhibitors act independently by targeting different sites of action and without interfering with each other’s activity; and 3) Both drugs contribute to the observed effect of latent HIV reactivation. The Calculation Index (CI) for synergy measurement was determined by the equation **(E_A_ + E_B_ – E_A_E_B_)/E_AB_**, where **E_A_** is the observed % effect of Drug A, **E_B_** is the % effect of Drug B and **E_AB_** is the % effect of the combined treatment ^56, 57^. Synergy, antagonism or additivity was determined by CI under, above or near equal to 1, respectively.

### Virus release single copy assay

De-identified, cryopreserved PBMCs belonging to people with HIV that were on antiretroviral therapy were obtained through the Rustbelt CFAR Clinical Core Repository. Approximately 3 million memory CD4^+^ T cells were isolated from 30 million PBMCs and equally divided into 3 24-wells at 1 million cells per ml. Following a 24 h challenge with 50 nM ingenol-3-angelate (ingenol) or a combination of 500 nM 2A127 and 500 nM SAHA, cells were centrifuged at 500 xg for 5 min in sterile 1.5-ml screw cap microcentrifuge tubes. Cell culture supernatants were carefully harvested without touching the cell pellets and passed through a low protein-binding cellulose acetate 0.45 μm filter by centrifugation at 13,000 rpm for 1 min. For the subsequent RT-qPCR measurements, cell media supernatants belonging to uninfected memory CD4^+^ T cells and a CEMx174 5.25 reporter cell line infected with the NL-AD8 HIV isolate were used as negative and positive controls, respectively. A modified version of the RT-qPCR assay with single-copy sensitivity targeting the integrase region of HIV (integrase single-copy assay v2.0) developed by Tosiano et al. ^58^, was employed to measure the extent of virus release following LRA treatment. Briefly, the filtered cell culture supernatants were subjected to centrifugation at 21,000 xg for 2 h in a refrigerated Eppendorf high speed microcentrifuge. Following centrifugation, media was removed by pipetting from the top leaving a uniform residual volume of 50 μl for each sample. The Allprep RNA/DNA mini kit was then used to lyse the pelleted virions and isolate RNA. Reverse transcription was carried out using an integrase-specific reverse primer (5’-CCTGCCATCTGTTTTCCA-3’) with the QuantiTect Reverse Transcription kit. Real time PCR reactions were set up in 96-wells in triplicate or sextuplicate using the BlasTaq Probe 2X qPCR Mastermix (Applied Biological Materials). Reactions were run on a QuantStudio 3 Real-Time PCR System with the following thermal cycling conditions: an initial denaturation at 95°C for 3 min, followed by 40 cycles of 95°C for 15 sec and 60°C for 60 sec. Triplicates of two-fold serial dilutions of a cDNA sample prepared from the NL-AD8 HIV isolate were also included to generate the standard curve. The primer set and probe used for qPCR were as follows: (Forward: 5’-TTTGGAAAGGACCAGCAAA-3’; Reverse: 5’-CCTGCCATCTGTTTTCCA-3’; Probe: FAM –AAAGGTGAAGGGGCAGT– MGB NFQ). Data analysis was performed using the Design and Analysis 2 software (ThermoFisher).

### Measurement of cell-associated HIV RNA by RT qPCR

Latently infected primary Th17 cells generated using the QUECEL protocol were placed in 24-wells at 1 million cells per well and treated or not for 24 h with various candidate LRAs as follows: 50 nM ingenol, 50 ng/ml PMA, 1 μM prostratin, 500 nM SAHA, or activation through the T-cell receptor with bead-bound anti-CD3 and anti-CD28 antibodies. After harvesting the cells in 1.5 ml screw cap tubes by microcentrifugation at 500 xg for 5 min, total cellular RNA and genomic DNA were isolated using the Allprep RNA/DNA Mini kit (Qiagen). For each sample, 200 ng of RNA was used to prepare a cDNA library using the QuantiTect Reverse Transcription kit (Qiagen) by following the manufacturer’s recommendations for the reaction setup and RT conditions. For qPCR, the cDNA products were diluted 10-fold with nuclease-free water and a total volume of 20 μl of the PCR reaction was prepared by mixing 10 μl of FastStart Universal SYBR Green Master (Millipore Sigma), 4 μl of nuclease-free water, 5 μl of sample cDNA , and 1 μl of HIV-specific primer mix designed to detect unspliced transcripts (Forward: 5’-GGGTGCGAGAGCGTCGGTATTAAGC-3’; Reverse: 5’-TCCTGTCTGAAGGGATGGTTGTAGC-3’) or multiply spliced transcripts (Forward: 5’-CTTAGGCATCTCCTATGGCAGGAA-3’; Reverse: 5’-TCAAGCGGTGGTAGCTGAAGAGG-3’). The qPCR reactions were set up in triplicate for each sample and carried out in the QuantStudio 3 Real-Time PCR system (Thermofisher) using the following reaction conditions: Initial phase – 50°C for 2 min then 95°C for 10 min; PCR phase – 40 cycles of 95°C for 10 min then 60°C for 1 min; Melting curve – 95°C for 15 min then 60°C for 1 min and 95°C for 1 sec. Total proviral DNA measurements were also performed by qPCR using the following primer set targeting the EGFP reporter gene (Forward: 5’-AGCAGAAGAACGGCATCAAG-3’; Reverse: 5’-CTCCAGCAGGACCATGTGAT-3’). For normalization, HIV RNA/DNA ratio for each sample was calculated using the following equation: 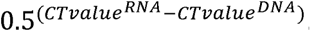.

### Immunofluorescence flow cytometry

Experimentally treated primary Th17 or memory CD4^+^ T cells were placed in polystyrene flow tubes, harvested by centrifugation at 1500 rpm for 3 min and washed with 1X PBS prior to fixation in 4% formaldehyde for 15 min at room temperature (RT). Fixed cells were then permeabilized by a sequential treatment with 0.2% Triton-X-100 for 10 min followed by incubation with 1X BD Perm/Wash buffer (BD Biosciences) for 15 min at room temperature. Following a 15-min blocking step with 10 µg of a non-specific IgG, permeabilized cells were immunostained for 45 min in the dark and at room temperature with fluorophore-conjugated antibodies directed against CD25, CD69, Cyclin T1, pSer175 CDK9, pSer2 RNAP II CTD, Nef, or Tat. For antibodies that were unconjugated, Alexa Fluor antibody labeling kits (ThermoFisher Scientific) were used for fluorophore conjugation as previously described ^19^. The sources of these reagents are listed in **Supplementary Table 2**. Tat monoclonal 2D1.1 and pSer175 CDK9 antibodies were custom-synthesized; their generation, purification and validation have been described previously ^18, 19, 111^. Immunostained cells were rinsed with 1X PBS then subjected to flow cytometry analysis using the BD LSR Fortessa instrument and BD FACSDiva software (BD Biosciences) with the appropriate compensation settings. At least 10,000 events were captured for each fluorescently stained cell sample. The resulting raw flow data in FCS file format was analyzed using Winlist 3D version 9.0.1 and the processed data with the appropriate gating were presented as either 2D scatterplots or 1D histograms. The extent of target protein expression was reported as a percentage of cells that were determined to be fluorescently positive relative to the appropriate gating control (isotype control, or a fluorescently stained unstimulated or uninfected control). Flow gating strategies employed in the current study are shown in **Supplementary Fig. 22**.

### RNA FISH staining of primary T cells

3.0 X 10^5^ primary T cells per treatment were harvested by centrifugation at 1500 rpm for 3 min, washed with 1X PBS and resuspended in 30 μl 1X PBS. The cell suspension was placed onto poly-L-lysine-coated coverslips in a 24-well plate and allowed to adhere to the surface at 37°C for 15 min. Cells were fixed with 4% formaldehyde for 15 min, washed with 1X PBS and then permeabilized using 70% ethanol for 1 h at 4°C. After permeabilization, coverslips were placed in Wash Buffer A (2X SSC and 10% formamide in RNase-free ultrapure water) for 5 min at room temperature. Stellaris custom-synthesized fluorophore-conjugated oligonucleotide probe sets (**Supplementary Table 3**) were resuspended in hybridization buffer (10% dextran sulfate, 2X SSC and 10% formamide in RNase-free ultrapure water) to a final concentration of 125 nM. Coverslips were inverted onto 40 μl droplets of the hybridization mixture in a humidified chamber which was then tightly sealed and incubated in the dark at 37°C for 16 h. Afterwards, coverslips were transferred onto a fresh 24-well plate cells side up, incubated in the dark with Wash buffer A for 30 min and then counterstained for 15 min with 1 μg/ml DAPI that was prepared in Wash Buffer A. Coverslips were washed twice with Wash Buffer B (2X SSC in RNAase-free ultrapure H_2_O) prior to mounting onto glass slides using Prolong Antifade medium (Thermofisher).

### Combined immunofluorescence and RNA FISH staining

Primary T cells attached to poly-L-lysine-coated coverslips were fixed with 4% formaldehyde for 15 min, washed with 1X PBS and then permeabilized with a sequential treatment of 0.2% Triton-X-100 for 10 min and 1X Perm/Wash buffer for 30 min at room temperature. For the blocking step, cells were incubated in blocking solution for 30 min made up of 1% BSA (10 μg/ml) and 50 μg/ml Donkey IgG in 1X Perm/Wash. Thereafter, cells were incubated with primary antibody solutions prepared in blocking buffer for 2 h at room temperature, washed three times with 1X Perm/Wash buffer then incubated with 1:200 dilutions of Alexa Fluor-conjugated anti-rabbit secondary antibodies that were also prepared in blocking buffer. After immunostaining, the coverslips were washed twice with 1X PBS and re-fixed in 4% formaldehyde solution prior to initiation of the RNA FISH procedure. Coverslips were washed once with 1X PBS before incubation with Wash Buffer A (2X SSC and 10% formamide in RNase-free ultrapure water) for 5 min at room temperature. For RNA FISH, coverslips were inverted onto 40 μl droplets of the hybridization mixture containing fluorophore-conjugated oligonucleotide probe sets (**Supplementary Table 3**) and incubated in a dark humidified chamber at 37°C for 16 h. Afterwards, coverslips were transferred onto a fresh 24-well plate cells side up, incubated in the dark with Wash buffer A for 30 min and then counterstained for 15 min with 1 μg/ml DAPI that was prepared in Wash Buffer A. Coverslips were washed twice with Wash Buffer B (2X SSC in RNAase-free ultrapure H_2_O) prior to mounting onto glass slides using Prolong Antifade medium.

### Deconvolution fluorescence microscopy

Images were captured at high magnification (100X) by deconvolution microscopy using a DeltaVision epifluorescent microscope (Applied Precision). Raw *z* stack images were deconvolved and processed using the softWoRx analysis program (Applied Precision). For colocalization measurements, the softWoRx colocalization module was used to create correlation scatterplots and measure the Pearson’s coefficient. A region of interest for the colocalization analysis was defined as a single cell. Processed images were exported as JPEG files and micrographs were composed using Adobe Photoshop. The resulting TIFF files of the micrographs were subjected to fluorescence densitometry for each captured channel (FITC, Cy5, TRITC, or DAPI) using Quantity One (Bio-Rad).

### RNA-seq processing and transcript quantification

Raw RNA sequencing (RNA-seq) reads from publicly available bulk RNA-seq datasets of primary human memory CD4^+^ T cells (SRA accession SRP026389) ^33^ and QUECEL effector subsets (SRA accession SRP145508) ^54^, were subjected to quality control and adapter trimming using fastp (v1.4.0). Adapter sequences and low-quality bases were removed using the default filtering parameters, and only high-quality reads were retained for downstream analyses. For alignment-based analyses, trimmed reads were aligned to the GRCh38 primary human genome assembly using STAR (v2.7.9a) with the GENCODE release 50 reference annotation. Gene and transcript abundance estimates used in all downstream expression analyses were obtained using Salmon (v1.10.3) against a decoy-aware transcriptome index generated from the GENCODE release 50 transcriptome and the corresponding GRCh38 primary genome assembly. Transcript quantification was performed using automatic library type detection (-l A), selective alignment (--validateMappings), and sequence-specific (--seqBias) and GC-content (--gcBias) bias correction.

Transcript-level abundance estimates (quant.sf) were imported into R (v4.4) using tximport (v1.34.0). Transcript abundances were summarized to gene-level expression using transcript-to-gene mappings derived from the GENCODE release 50 annotation. Ensembl transcript and gene version suffixes were removed prior to transcript-to-gene mapping to ensure consistent identifier matching across all samples, and gene symbols were assigned using the corresponding GENCODE annotations. Gene-level transcripts per million (TPM), estimated counts, and effective transcript lengths were generated for downstream analyses, while transcript-level TPM matrices were retained for isoform-specific analyses where appropriate.

### Preparation of primary T-cell extracts for Western blotting

Experimentally treated memory CD4^+^ T cells were harvested and washed in ice-cold 1X PBS then lysed in cell lysis buffer A (150 mM NaCl, 10 mM KCl, 1.5 mM MgCl_2_, 0.5% Nonidet P-40, 1 mM DTT, 10 mM Hepes, pH 8.0) containing an EDTA-free protease and phosphatase inhibitor mixture (Roche Life Science). After clearing the whole cell extracts (WCE) by centrifugation at 5000 rpm for 5 min, an equivalent volume of sample across the experimental treatments were boiled in 1X LDS sample loading buffer containing 50 mM DTT. SDS-PAGE of the WCE was performed on 4-12% Bis Tris gels in MOPS or MES buffer at 150 V for 70 min, followed by transfer of resolved proteins to PVDF membranes. Thereafter, Western blotting was performed with primary antibodies against pThr202/Tyr204 ERK1/2, total ERK1/2, pThr389 p70S6K and pSer235/Ser236 rpS6 (Cell Signaling Technology).

### Statistics and data analysis

Box-and-whisker plots for fluorescence and flow cytometry data were prepared in Origin (OriginLab Corp). Microsoft Excel was used to generate bar graphs for the Western blotting and cell viability measurements. Error bars were generated from at least three independent experiments with findings presented as mean +/- standard error. Comparisons between two experimental groups were conducted using a two-tailed Student’s t-test, with p < 0.05 considered to be statistically significant.

## ACKNOWLEDGMENTS

This work used the CWRU/Pitt Rustbelt Center for AIDS Research flow cytometry, microscopy and clinical cores. This work was supported by National Institutes of Health grants R01 AI148083 to J.K. and U.M., R61 AI169629 to J.K. and a developmental grant to U.M. from the CWRU/Pitt Center for AIDS Research (P30 AI036219).

## AUTHOR CONTRIBUTIONS

U.M. and J.K conceived the study. U.M. designed the study, performed experiments, analyzed data, supervised some of the experiments, prepared the figures and wrote the manuscript. A.B., E.E., L.G.D., and M.J.C. synthesized all the DAG-indololactones and measured their in vitro binding affinities. M.Y. performed the RT-qPCR experiments to measure latent HIV reactivation in the QUECEL model (Figure 1E and Supplementary Fig. 1C), prepared the correlation plots presented in Figure 1E, and conducted the flow cytometry experiments shown in Supplementary Fig. 4D. A.A. analyzed the abundance of the mRNA species shown in Supplementary Fig. 5 using publicly available bulk RNA-seq datasets. A.D. conducted the HIV RNA FISH experiment shown in Fig. 1F and Supplementary Fig. 4B and assisted in the routine preparation of the QUECEL model. A.B., E.E., M.Y., A.D. and J.K. read the manuscript and provided comments. U.M. and J.K. edited the manuscript.

## DATA AVAILABILITY

All data supporting the findings of this study are available within the paper and its supplementary files. Source data is also provided with this paper.

## SUPPLEMENTARY FIGURES

**Supplementary Figure 1. Posttranscriptional induction of CycT1 and P-TEFb kinase activity are required for latent HIV reactivation and the synthesis of Tat.** Reactivation of latent HIV in the QUECEL primary Th17 model by TCR co-stimulation (A) or natural PKC agonists and HDAC inhibitors (B) is abrogated by the CDK9 kinase inhibitor flavopiridol (FVP at 100 nM). Proviral reactivation in (B) was measured by dual immunofluorescence detection of reporter EGFP and Tat (*Left graph*) or the dual detection of Tat and Nef (*Right graph*). Results shown in (A) and the left graph in (B) are from a flow experiment conducted with QUECEL primary Th17 cells prepared using naïve T cells from donor D18. Results shown in the right graph in (B) are from a flow experiment conducted with QUECEL primary Th17 cells prepared using naïve T cells from donor D25. (C) RT-qPCR measurements of latent HIV reactivation in QUECEL primary Th17 assessing the expression of unspliced and multiply spliced viral transcripts. P-TEFb inhibition with 100 nM FVP blunts the formation of these transcripts in response to TCR co-stimulation. These measurements were conducted with QUECEL primary Th17 cells prepared using naïve T cells from donor D26. (D) Western blot showing that Tat synthesis is completely blocked by 100 nM FVP in the Jurkat 2D10 clonal model. (E) Reactivation of latent HIV by natural PKC agonists is abrogated by 100 nM FVP in the Jurkat 3C9 clonal model. Graphed results are from four experiments. (F) Posttranscriptional induction of CycT1. RNA FISH and immunofluorescence staining of memory CD4^+^ T cells to examine the subcellular co-expression of CycT1 mRNA and CycT1 protein with or without TCR co-stimulation. This imaging experiment was conducted using memory CD4^+^ T cells belonging to donor D24. (G) Signal-dependent assembly of 7SK snRNP. RNA FISH and immunofluorescence staining of memory CD4^+^ T cells to examine the sub-cellular co-expression of 7SK snRNA and LARP7 protein with or without TCR co-stimulation. This imaging experiment was conducted using memory CD4^+^ T cells belonging to donor D16. (G) Combined 7SK RNA FISH and Mettl16 immunostaining intended to illustrate their differential nuclear localization patterns in unstimulated and TCR-activated memory CD4^+^ T cells. This imaging experiment was conducted using memory CD4^+^ T cells belonging to donor D16. (H) RNA FISH to examine the subcellular distribution of CycT1 and CD69 transcripts in unstimulated memory CD4^+^ T cells. Six representative micrographs are shown. This imaging experiment was conducted using memory CD4^+^ T cells belonging to donor D23.

**Supplementary Figure 2. Proteasomal inhibition modestly elevates CycT1 expression whereas translation inhibition abrogates TCR-induced CycT1 expression in memory T cells.** (A) Cell viability measurements showing that a 12-h treatment with 50 µg/ml cycloheximide (CHX) or 5 µM MG132 is minimally toxic to memory CD4^+^ T cells. Plotted data are from triplicate propidium iodide staining measurements using cells from a single healthy donor. (B) MG132 modestly induces CycT1 expression, whereas TCR-induced CycT1 expression is effectively blocked by CHX. Cells were treated for 8 h with MG132 (5 or 10 µM) or CHX (50 µg/ml) prior to measuring CycT1 or Cyclin D3 (CycD3) levels by intracellular immunofluorescence flow cytometry. In parallel, cells were activated via the TCR for 8 h in the absence or presence of 50 µg/ml CHX. Plotted data for CycT1 are representative of four independent experiments using primary memory CD4**^+^** T cells from two healthy donors (D27 and D30) and ex vivo polarized QUECEL primary Th17 cells. Plotted data for CycD3 are representative of three independent experiments using D27 and D30 memory CD4^+^ T cells. Statistical significance was calculated using a two-tailed Student’s t test. (C) Representative flow cytometry plots of the graphed data shown in (B).

**Supplementary Figure 3. Comparison of CycT1 and CycD3 protein stability in activated memory CD4^+^ T cells.** (A) TCR activation for 10 h followed by 50 µg/ml CHX treatment for the times shown. CycT1 and CycD3 protein decay was evaluated across a 2-hour chase window since the inhibition of translation post-activation revealed a limit for conducting CHX chase as between 2 and 4 h post-CHX treatment (indicated by the vertical blue dashed line in the bottom graphs). (B) Proteasomal inhibition prevents the loss of CycD3 but does not impact CycT1 expression. Cells activated via the TCR for 10 h were treated for 2 h with 50 µg/ml CHX in the presence or absence of 10 µM MG132. Representative flow plots of the graphed data are shown in (A) and (B). Plotted data are representative of at least four independent experiments using primary memory CD4**^+^** T cells from two healthy donors (D27 and D30) and ex vivo polarized QUECEL primary Th17 cells. Statistical significance was calculated using a two-tailed Student’s t test.

**Supplementary Figure 4. Formation of hyperphosphorylated RNAP II and latent HIV reactivation coincide with the biogenesis of P-TEFb.** (A) Flow plots representative of the data shown in Figure 1D. P-TEFb expression was measured by the dual detection of CycT1 and pSer175 CDK9. Hyperphosphorylated RNAP II was detected by immunofluorescence staining for pSer2 RNAP II CTD. The concentrations of stimuli used were as follows: 50 nM ingenol-3-angelate (ingenol); 1:1 bead-to-cell ratio anti-CD3/CD28 Dynabeads; and 1 µg/ml ionomycin. (B) *Top panel*, Representative micrographs of the HIV RNA FISH results shown in Figure 1F. Bottom panel, Representative histogram flow plots of the CycT1 immunofluorescence data are shown in Figure 1F. (C-E) Representative flow plots of the graphed data shown in Figure 1F showing measurement of Tat or Nef protein expression in QUECEL primary Th17 cells at varying times following TCR co-stimulation of the latently infected populations. Polarized Th17 cells were infected with a VSVG-pseudotyped pHR’-CD8a-EGFP-Nef+ single-round HIV construct, then subjected to a positive selection of acutely infected cells using the EasySep Mouse CD8a Positive Selection Kit II (StemCell) at Day 4 post-infection. Viral latency was established by enforcement of a quiescent phenotype through a 2-week culture of the positively selected cells. The CD8a-EGFP fusion protein is stably expressed on the cell surface and remains detectable in latently infected cells. (F) Peak co-expression of CycT1 and Tat 24 h after TCR co-stimulation of latently infected QUECEL primary Th17 cells. (G) Measurement of inducible HIV expression in QUECEL primary Th17 cells using a d2EGFP reporter. After infecting polarized Th17 cells with VSVG-pseudotyped packaged virus of the d2EGFP-carrying single-round HIV construct, viral latency was established by enforcement of a quiescent phenotype through a 2-week culture of the positively selected cells. Following TCR co-stimulation or treatment with either 10 nM bryostatin or 50 nM ingenol for the times shown, viral reactivation within the mixed population of cells was assessed by flow analysis of HIV LTR-driven d2EGFP expression.

**Supplementary Figure 5. Transcriptome analysis in memory CD4^+^ T cells and primary QUECEL subsets.** (A) Abundance of various mRNA species using RNA-seq reads derived from a publicly available bulk RNA-seq dataset (SRA accession SRP026389) generated using human memory CD4^+^ T cells that had been activated or not through TCR co-stimulation with anti-CD3/anti-CD28 beads for the times shown in two replicate experiments. The quantified transcripts shown are for cyclin T1 (CycT1), CD69, CD25, cyclin D3 (CycD3), cyclin D2 (CycD2), ELL2, FOS and GAPDH. (B) Examination of mRNA abundance from bulk RNA-seq datasets generated using human memory CD4^+^ T cells (Tm (Expt 1) and Tm (Expt 2); SRA accession SRP026389) and QUECEL polarized effector subsets (Th1, Th2, Treg and Th17; SRA accession SRP145508) that were activated or not for 24 h. Statistical significance was calculated using a two-tailed Student’s t test. Quantified measurements (A) and (B) are expressed as transcripts per million (TPM).

**Supplementary Figure 6. Differential regulation of the expression of the activation markers CD25 and CD69 in memory CD4^+^ T cells.** (A) Inhibition of CDK7-driven transcription substantially suppresses the inducible expression of CD25 but minimally affects the inducible expression of CD69. Memory CD4^+^ T cells were pre-treated or not with 50 nM THZ1, a CDK7 covalent inhibitor, for 30 min prior to challenge with 50 nM ingenol for 24 h. Thereafter, cells were fixed, permeabilized then subjected to immunofluorescence staining and flow cytometry to examine CD25 and CD69 protein expression. (B) Induction kinetics of CD69 and CD25 protein expression following treatment of memory CD4^+^ T cells with the PKC agonists (PKC Ag) ingenol (50 nM) or prostratin (1 µM). Included in the plotted data are results from two complete time course experiments conducted with memory T cells belonging to two donors (D25 and D27). (C) Transcriptional induction of CD25 expression in memory CD4^+^ T cells is sensitive to the CDK7 inhibitor THZ1 but refractory to inhibition by flavopiridol (FVP). Cells were pre-treated or not with 100 nM FVP or 50 nM THZ1 for 30 min prior to stimulation with the PKC agonists (PKC Ag) ingenol (50 nM) or prostratin (1 µM) for 24 h. Immunofluorescence flow cytometry data plotted in (B) and (C) are from at four independent experiments performed with memory T cells belonging to three donors (D25, D27 and D30). Statistical significance was calculated using a two-tailed Student’s t test. (D) Inhibition of transcription fails to suppress early induction of CD69 protein expression. Memory CD4^+^ T cells were pre-treated or not for 30 min with 50 nM THZ1, 1 μM actinomycin D or a combination of 10 μM U0126 and 100 nM sotrastaurin. Thereafter, cells were stimulated with 50 nM ingenol or 1 μM prostratin for 2 h prior to immunofluorescence staining and flow cytometry to examine CD69 protein expression.

**Supplementary Figure 7. CycT1 protein synthesis in primary T cells induced by natural PKC agonists is mediated by MEK-ERK signaling and not PKC enzymes.** (A) Inhibition of MEK with U0126 substantially blocks ingenol- or prostratin-induced translation of CycT1 but modestly represses TCR-induced CycT1 protein expression. Health donor memory CD4^+^ T cells (obtained from donors D16 and D17) and uninfected QUECEL primary Th17 cells (derived from donors D15 and D16) were pre-treated or not with the MEK inhibitor U0126 (at 10 and 20 µM) for 30 min prior to 24 h stimulation through the TCR or with 50 nM ingenol or 1 µM prostratin. Plotted data are from five independent experiments. (B) Inhibition of PKC with sotrastaurin (100 nM) or Ro-31-8220 (100 nM) minimally affects CycT1 protein synthesis induced by TCR co-stimulation, 50 nM ingenol or 1 µM prostratin. Plotted data represent eight independent experiments. Seven experiments used memory CD4^+^ T cells from four donors (D9, D10, D15 and D16): five tested the PKC-θ inhibitor sotrastaurin (at 100 nM), and two tested the pan-PKC inhibitor Ro-31-8220 (at 100 nM). One experiment used uninfected QUECEL primary Th17 cells derived from donor D14 and tested the PKC-θ inhibitor sotrastaurin (at 100 nM). (C and D) Effect of the inhibition of MEK on the expression of CD69 and CD25 induced by TCR co-stimulation, 50 nM ingenol or 1 µM prostratin. Data in (C) are from 11 independent experiments with each experimental treatment performed in two or more biological replicates. Data in (D) are from seven independent experiments with each experimental treatment performed in two or more biological replicates. The donors and primary cell types used in (C) and (D) are shown in the source data file. (E) Inhibition of MEK suppresses bryostatin-induced CD69 expression. Health donor memory CD4^+^ T cells (obtained from donor D16) and uninfected QUECEL primary Th17 cells (derived from donor D9) were pre-treated or not with U0126 (at 10 and 20 µM) for 30 min prior to 24 h stimulation with 10 nM bryostatin. Plotted data are from three independent experiments. (F) Inhibition of PKC with sotrastaurin modestly suppresses prostratin-induced CD69 expression. Health donor memory CD4^+^ T cells (obtained from donors D16 and D25) were pre-treated or not with 100 nM sotrastaurin for 30 min prior to 24 h stimulation with 1 µM prostratin. Plotted data are from three independent experiments. (G) Combined inhibition of MEK and PKC-θ abrogates ingenol-induced CD69 expression. D16 memory CD4^+^ T cells were pre-treated or not with 10 µM U0126 or 100 nM sotrastaurin or a combination of both inhibitors for 30 min prior to 24 h TCR stimulation or challenge with 50 nM ingenol. In (A) through (G), CycT1, CD25 and CD69 protein expression was examined by immunofluorescence flow cytometry. In (A) through (F) statistical significance was calculated using a two-tailed Student’s t test.

**Supplementary Figure 8. Measurement of latent HIV reactivation by DAG-lactones in Jurkat cell line models.** (A) Representative histogram plots showing the extent of latent HIV reactivation in the Jurkat 3C9 wildtype HIV clone upon 24 h treatment with increasing concentrations of the parent DAG lactone AJH836 or the DAG-indololactone derivatives. (B) Graphs of composite data from five independent experiments showing the extent of latent HIV reactivation in the Jurkat 3C9 clone upon 24 h treatment with increasing concentrations of DAG-lactones. (C) Examination of latent HIV reactivation in Jurkat clonal models with a wildtype proviral promoter (2D10 and 3C9) or a defective proviral promoter (2B5) following treatment with increasing concentrations of DAG-lactones. Each data point represents an average calculated from at least three independent experiments. (D) Effects of 24 h DAG-lactone treatments on the viability of Jurkat cells. Plotted data shows % cell viability relative to the untreated condition and is a compilation of eight independent experiments.

**Supplementary Figure 9. Synthetic DAG-lactones stimulate CycT1 translation and induce active P-TEFb with minimal elevation of T-cell activation markers.** Memory CD4^+^ T cells derived from two donors, an adult male (D23) and adult female (D24), were treated with varying concentrations of AJH836 (A), 2C76 (B), or 2D168 (C). Afterwards, cells were immunofluorescently co-stained for transcriptionally active P-TEFb (dual staining for CycT1 and pSer175 CDK9), CD69 and CD25. Cyan and brown data points are from two independent experiments conducted with donor D24 cells; the other data points are from independent experiments carried out with donor D23 cells. Statistical significance was calculated for each treatment relative to the untreated condition (No stim) using a two-tailed Student’s t test. *N.S.* (not statistically significant) indicates a *p* value > 0.05. (D) Representative 2D dot plots of the graphed flow data presented in Figure 3.

**Supplementary Figure 10. DAG-indololactones and their combinations with HDAC inhibitors are well tolerated by memory CD4^+^ cells.** (A) Viability assessments of 24 h treatments with DAG-indololactones or natural PKC agonists in memory CD4^+^ T cells. Graphed data are from at least three independent experiments conducted using cells belonging to donors D23 and D24. (B) Effect of DAG-indololactone and HDACi combination treatments on the viability of memory CD4^+^ T cells (*Top graph*) or QUECEL primary Th17 cells (*Bottom graph*). Cell viability in (A) and (B) was measured after propidium iodide staining.

**Supplementary Figure 11. Latent HIV reactivation profiles as assessed by dual Tat and Nef immunofluorescence flow cytometry.** Uninfected, latently infected, and reactivated QUECEL primary Th17 cells were immunostained for Tat and Nef following treatment with various latency-reversing stimuli (A) or treatment with varying concentrations of the DAG-lactone AJH836 (B). The two-dimensional plots show proviral gene expression as monitored by CD8a-EGFP versus Tat *(Top)*, CD8a-EGFP versus Nef *(Middle)*, and Tat versus Nef *(Bottom)*.

**Supplementary Figure 12. DAG-indololactones synergize with HDAC inhibitors to reactivate latent HIV in the QUECEL primary Th17 cell model.** (A) Effect of combination treatments of 2A127 and the HDACi romidepsin (RMD) on latent HIV reactivation in the QUECEL model. Plotted flow data are from four independent experiments conducted with QUECEL primary Th17 cells generated using naïve T cells derived from two donors. (B) Effect of combination treatments of 2C78 and the HDAC inhibitors SAHA, panobinostat (PNB), or romidepsin (RMD) on latent HIV reactivation in the QUECEL model. Plotted flow data are from four or more independent experiments conducted with QUECEL primary Th17 cells generated using naïve T cells derived from two donors. In both (A) and (B) cells were treated as shown for 24 h prior to immunostaining and examination of dual Tat and Nef expression by flow cytometry. (C and D) Individual treatments of memory CD4^+^ T cells with 500 nM 2A127 or 500 nM 2C78 or in combination with HDACi minimally induce the expression of CD25 and CD69. Cells obtained from donors D24 and D29 were treated as shown for 24 h prior to immunostaining and examination of CD25 and CD69 expression by flow cytometry.

**Supplementary Figure 13. Unlike 2A127, natural PKC agonists and the synthetic DAG-lactone AJH836 selectively induce a drastic loss of the CD4 receptor in memory CD4^+^ T cells.** (A) Natural PKC agonists and the synthetic DAG-lactone AJH836 induce a drastic loss of the CD4 receptor in memory T cells from donor D25, in contrast to the DAG indololactone 2A127 which modestly decreases CD4 expression. (B and C) Treatment with ingenol downregulates the expression of CD4, but not CD28, CD3 or CD45-Ro in memory T cells from donor D23. (D) Combination treatments of 2A127 and HDACi minimally impact CD4 expression in D25 memory CD4^+^ T cells. *Left panel*, Effects of 2A127 + SAHA or ingenol treatment on CD4 expression. *Right panel*, Effects of 2A127+ SAHA or ingenol treatment on CD3 expression. In (A-D) cells were treated as shown for 24 h prior to immunofluorescence staining of fixed and permeabilized cells.

**Supplementary Figure 14. PKC agonists, but not DAG-indololactones, induce CD69 elevation and the selective downregulation of CD4 in primary T cells.** Memory CD4^+^ T cells belonging to donor D30 were treated for 24 h as shown then fixed, permeabilized and fluorescently co-stained for CD3, CD4, CD69 and CCR5.

**Supplementary Figure 15. Combination of 2A127 and panobinostat robustly reactivates latent HIV in the QUECEL model with minimal CD4 downregulation and expression of T cell activation markers.** Uninfected QUECEL primary Th17 cells were treated as shown for 24 h prior to immunostaining and examination of CD4, CD25, and CD69 expression. Latently infected QUECEL primary Th17 cells were treated as shown for 24 h prior to examination of proviral HIV expression by Tat/Nef dual immunostaining.

**Supplementary Figure 16.** (A) Representative 2D dot plots of the graphed flow data presented in Figure 6A. (B and C) Inhibition of mTOR kinase suppresses the formation of transcriptionally active P-TEFb in response to challenge with 2A127 or prostratin. Memory CD4^+^ T cells belonging to donor D24 were pre-treated or not with the inhibitors shown for 30 min prior to treatment for 24 h with either 10 μM 2A127 or 1 μM prostratin. Thereafter, fixed and permeabilized cells were fluorescently co-stained for CycT1 and pSer175 CDK9.

**Supplementary Figure 17. DAG analogs stimulate the phosphorylation of S6K and rpS6 through the ERK1/2 signaling pathway.** (A) Memory CD4^+^ T cells belonging to donor D23 were treated with 50 nM ingenol or 10 nM bryostatin for the times shown then subjected to Western blotting of whole cell extracts to examine the phosphorylation of ERK1/2, S6K and rpS6. Total ERK1/2 was also analyzed as a normalization control. (B) Densitometry quantification of the effect of TCR co-stimulation on the expression of pT202/Y204 ERK1/2, pT389 p70S6K and pS235/S236 rpS6, with detected levels normalized to corresponding levels of ERK1/2. (C) Densitometry quantification of the effect of 50 nM ingenol on the expression of pT202/Y204 ERK1/2, pT389 p70S6K and pS235/S236 rpS6, with detected levels normalized to corresponding levels of ERK1/2. (D) Densitometry quantification of the effect of 10 nM bryostatin on the expression of pT202/Y204 ERK1/2, pT389 p70S6K and pS235/S236 rpS6, with detected levels normalized to corresponding levels of ERK1/2. The plotted average densitometry values in (B), (C) and (D) are from at least three independent Western blotting experiments performed using memory CD4^+^ T cells belonging to donors D23 and D24. Statistical significance values shown were calculated using a two-tailed Student’s t test. (E) Densitometry quantification of the Western blotting results shown in **Figure 6F** with levels normalized to total ERK1/2. (F) Densitometry quantification of the Western blotting results shown in **Figure 6G** with levels normalized to total ERK1/2.

**Supplementary Figure 18. Inhibition of mTOR and RSK suppresses the phosphorylation of rpS6 induced by DAG analogs.** (A) QUECEL primary Th17 cells were pre-treated or not with the following inhibitors for 30 min: 10 μM U0126, 50 nM Torin1, 500 nM rapamycin, 10 μM FMK and 10 μM BID1970. Thereafter, the cells were treated with 1 μM AJH836 or 50 nM ingenol for 24 h prior to preparation of whole cell extracts for Western blotting to examine the phosphorylation of rpS6. Total ERK1/2 was also analyzed as a normalization control. (B) Densitometry quantification of the Western blotting results shown in (A) for pS235/S236 rpS6 with levels normalized to total ERK1/2. (C) Healthy memory CD4^+^ T cells belonging to donor D27 were pre-treated or not with the following inhibitors for 30 min: 10 μM U0126, 50 nM Torin1, 500 nM rapamycin, 10 μM FMK and 10 μM BID1970. Thereafter, the cells were treated with 1 μM 2A127 or 1 μM 2C78 for 4 h prior to preparation of whole cell extracts for Western blotting to examine the phosphorylation of rpS6. Total ERK1/2 was also analyzed as a normalization control. (D) Densitometry quantification of the Western blotting results shown in (C) for pS235/S236 rpS6 with levels normalized to total ERK1/2.

**Supplementary Figure 19. Inhibition of mTOR but not RSK effectively retards the reactivation of latent HIV by DAG-indololactones in QUECEL primary Th17 cells.** (A) Two-dimensional plots of Tat/Nef dual immunofluorescence staining to examine the effect of various inhibitor treatments on latent HIV reactivation by 1 μM 2A127, 1 μM 2C78 and 1 μM AJH836. (B and C) Effect of individual or combination inhibitor treatments on latent HIV reactivation by 1 μM 2A127 as measured by Tat immunofluorescence staining. The results shown in (B) and (C) are from five independent experiments conducted with QUECEL primary Th17 cells that were prepared using naïve T cells from two donors (D23 and D27). Statistical significance was calculated using a two-tailed Student’s t test. (D) Effect of individual inhibitor treatments on latent HIV reactivation by 1 μM AJH836. The results shown are from two independent experiments conducted with QUECEL primary Th17 cells that were prepared using naïve T cells from donor D27. In (A) through (D) the inhibitor treatments were as follows: 10 μM U0126, 50 nM Torin1, 500 nM Rapamycin, 10 μM FMK, and 10 μM BID1970. Cells were pre-treated with these inhibitors for 30 min prior to a 24-h challenge with the DAG-lactones.

**Supplementary Figure 20. DAG analog-induced expression of CD69 and CD25 are refractory to the inhibition of mTOR in both memory CD4^+^ and QUECEL primary Th17 cells.** (A) Representative 2D dot plots of the graphed flow data shown in Figure 9. Cells were pre-treated with inhibitors at the indicated concentrations for 30 min prior to a 24-h challenge with1 µM prostratin. (B-D) Effect of individual or combination inhibitor treatments on the inducible expression of CD69 or CD25 in memory CD4^+^ or primary Th17 cells. Cells were pre-treated with inhibitors for 30 min prior to a 24-h challenge with1 µM AJH836 (B), 1 µM prostratin (C), or 50 nM ingenol (D). Inhibitor concentrations were as follows: 10 µM or 20 µM U0126, 50 nM Torin1, 500 nM Rapamycin, 10 µM FMK, 10 µM SL 0101 and 10 µM BID1970. Fixed and permeabilized cells were then subjected to immunofluorescent staining for CD69 and CD25. The experiment in (B) was performed with QUECEL primary Th17 cells prepared using naïve T cells from donor D27. The results shown in (C) are from four independent experiments conducted using memory CD4^+^ T cells from donors D9, D10, D24 and D25. The results shown in (D) are from three independent experiments performed with memory CD4^+^ T cells from donors D9, D10 and D24.

**Supplementary Figure 21. Viability measurements in uninfected QUECEL primary Th17 and memory CD4^+^ T cells.** Cells were challenged for 24 h with the DAG analogs as shown with or without inhibitor treatment. Cell viability in (A) was measured after propidium iodide staining of uninfected QUECEL primary Th17 cells prepared with naïve T cells belonging to donor D27. Cell viability in (B) was measured following eFluor 450 fixable dye staining of memory CD4^+^ T cells belonging to donor D25.

**Supplementary Figure 22. Flow gating strategies employed in the current study.** (A) Gating strategy for assessment of Tat/Nef dual immunofluorescence in QUECEL primary Th17 cells. Monoclonal antibodies towards Tat and Nef were conjugated with Pacific Blue (PB) and Alexa Fluor 647 (AF647) fluorophores, respectively. (B) Gating strategy for examination of inducible P-TEFb following staining of primary T cells with eFluor 450 fixable viability dye (FVD) and subsequent immunofluorescence staining of fixed and permeabilized cells for CycT1 and pSer175 CDK9. Antibodies towards CycT1 and pSer175 were conjugated with AF647 and AF488 fluorophores, respectively. (C) Gating strategy following staining of primary T cells with eFluor 450 FVD and subsequent immunofluorescence staining of fixed and permeabilized cells for P-TEFb, CD25 and CD69. (D) Gating strategy for the examination of inducible P-TEFb, CD25 and CD69 in primary T cells without eFluor 450 FVD staining.

