## Supplementary figures and images for "RasGRP1 agonism stimulates P-TEFb biogenesis via MEK-ERK-mTORC1 signaling to reverse HIV latency with minimal CD4 downregulation"

### Supplementary Figure 1

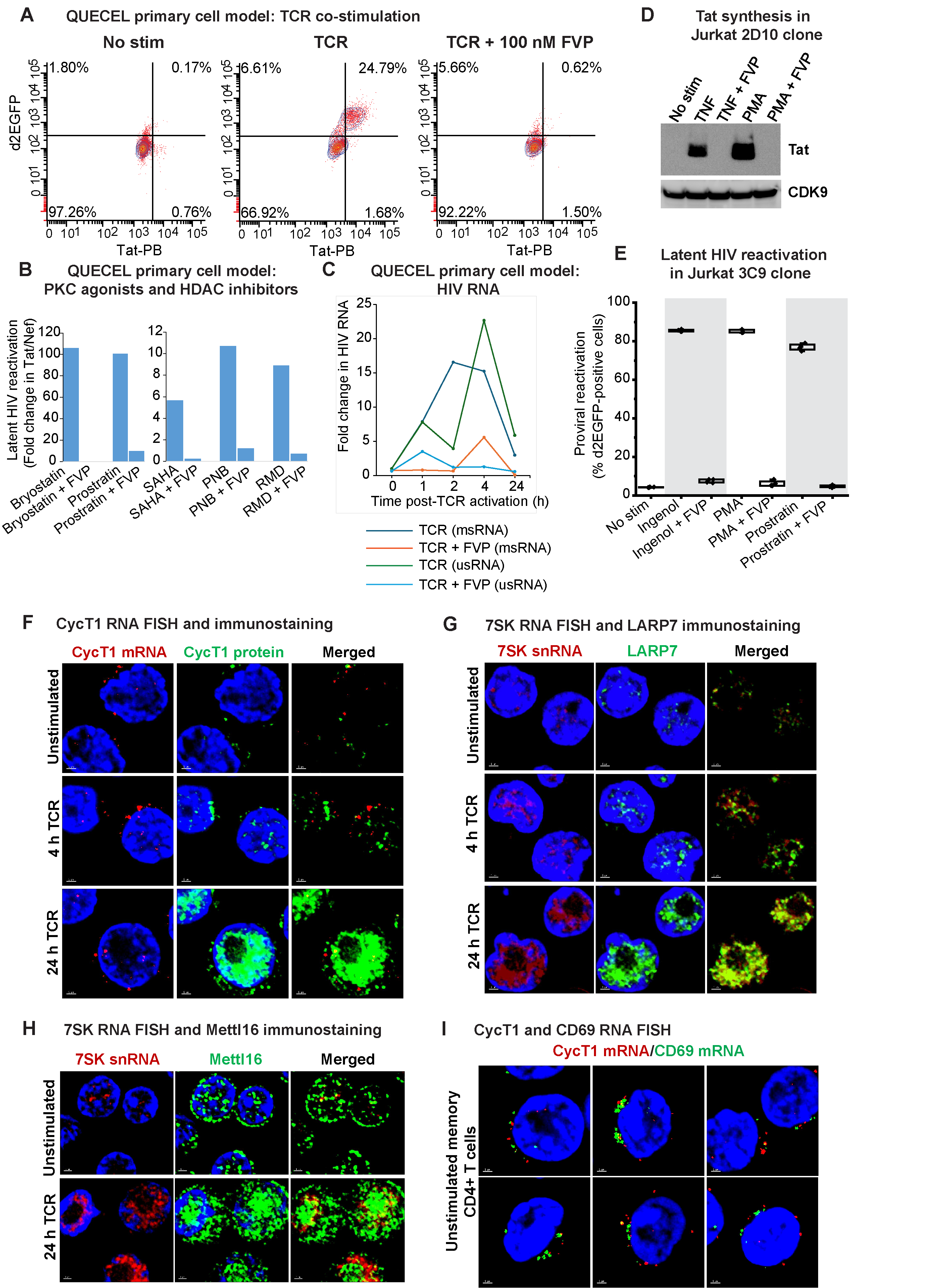

### Supplementary Figure 2

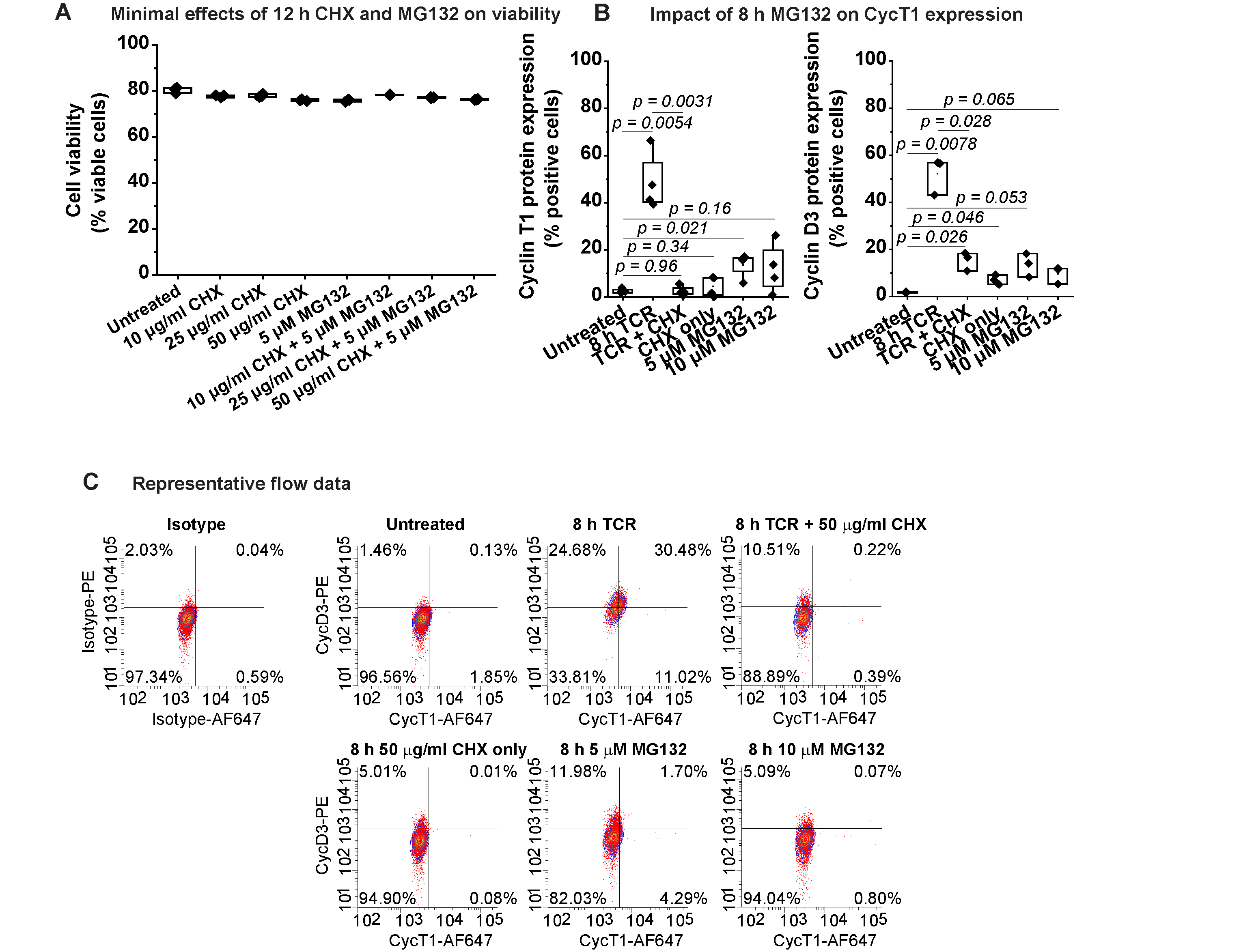

### Supplementary Figure 3

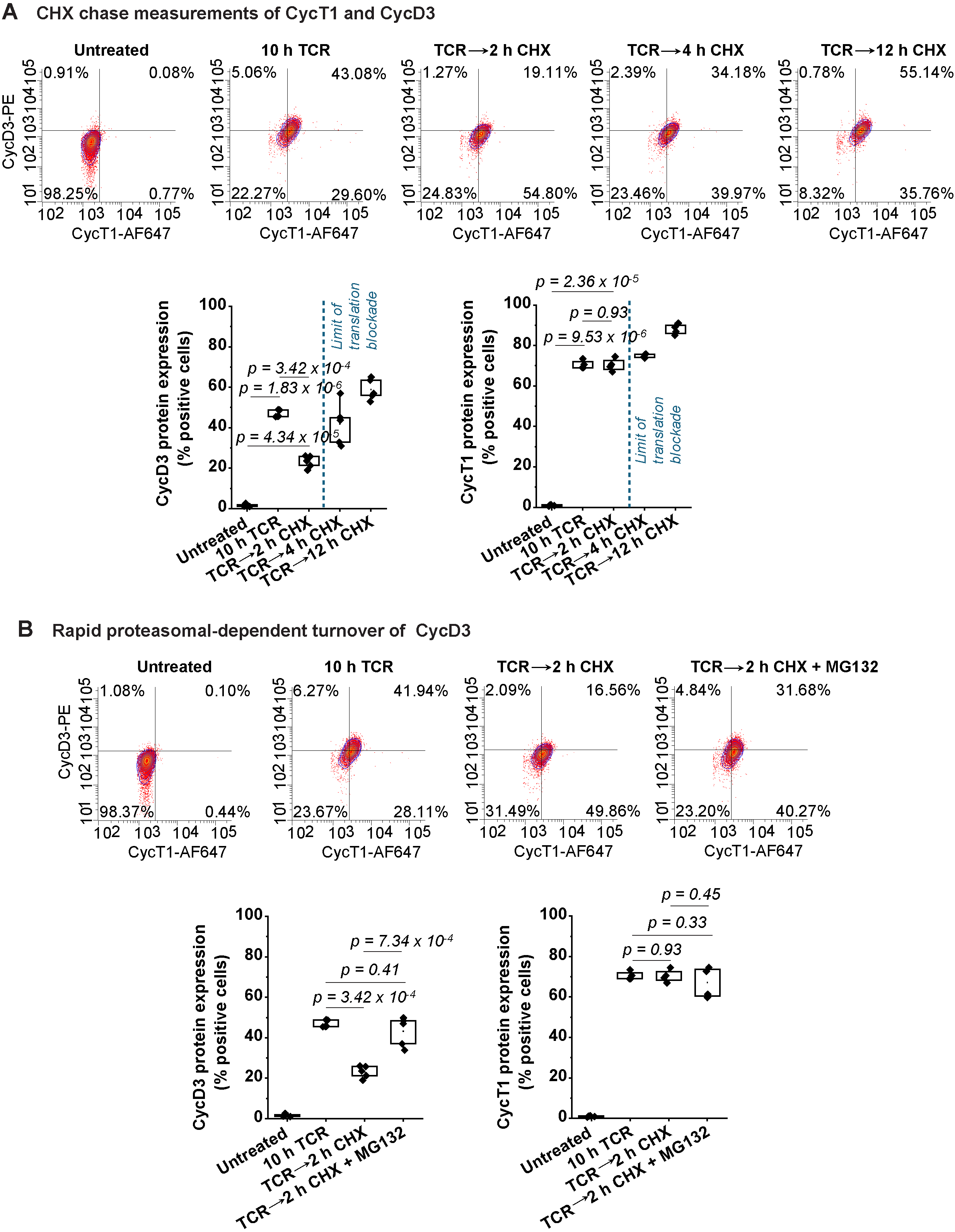

### Supplementary Figure 4A-B

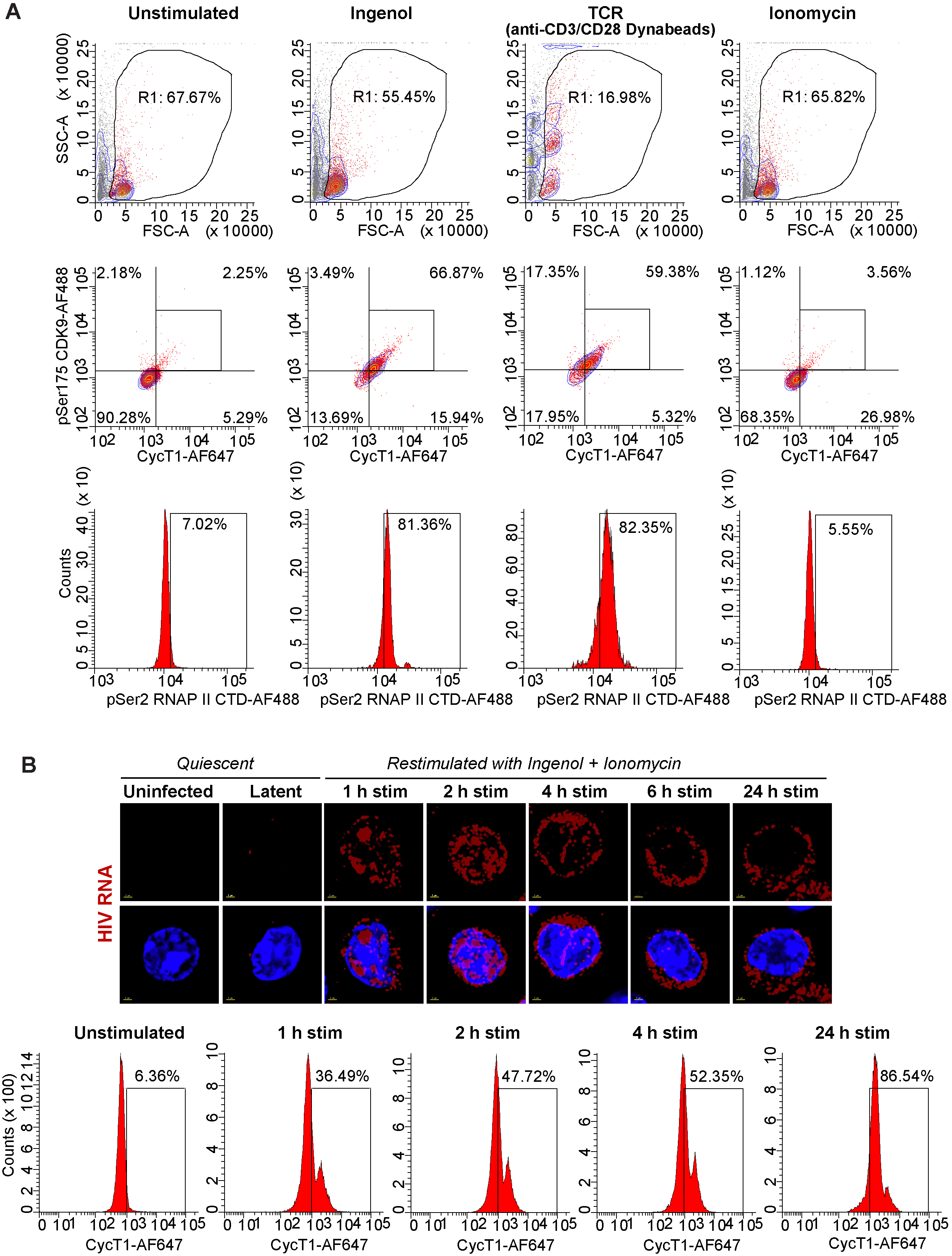

### Supplementary Figure 4C-F

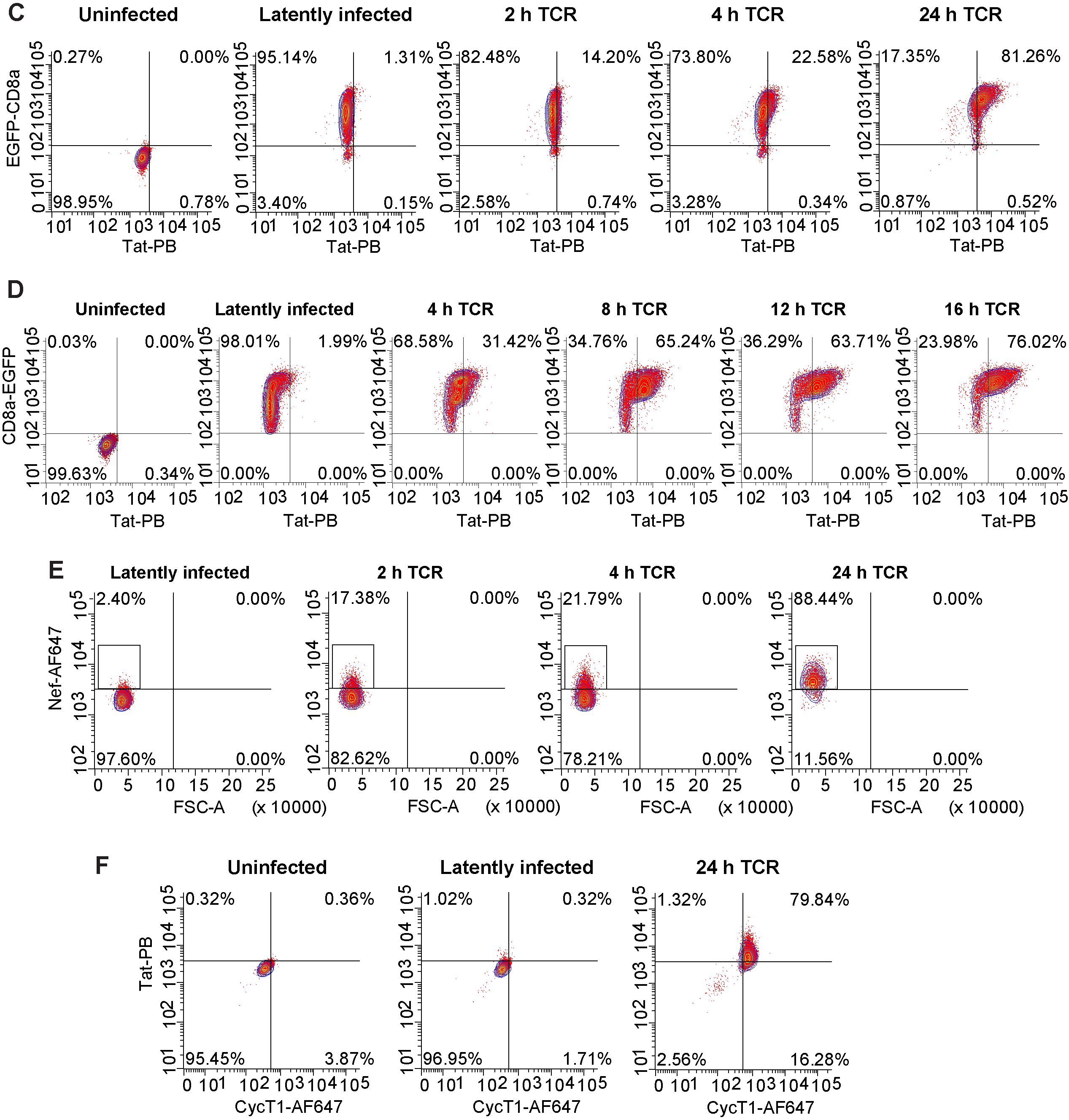

### Supplementary Figure 4G

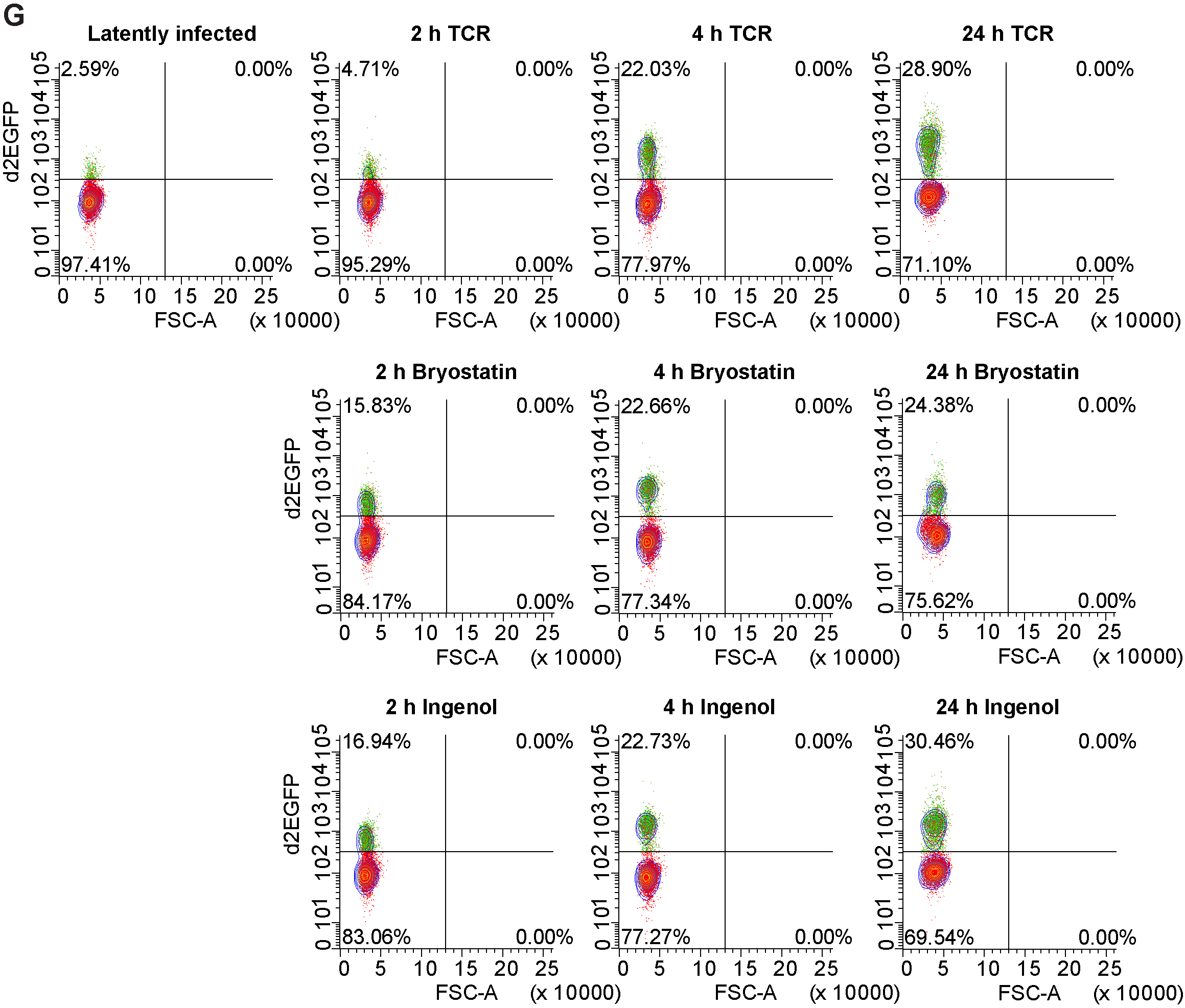

### Supplementary Figure 5

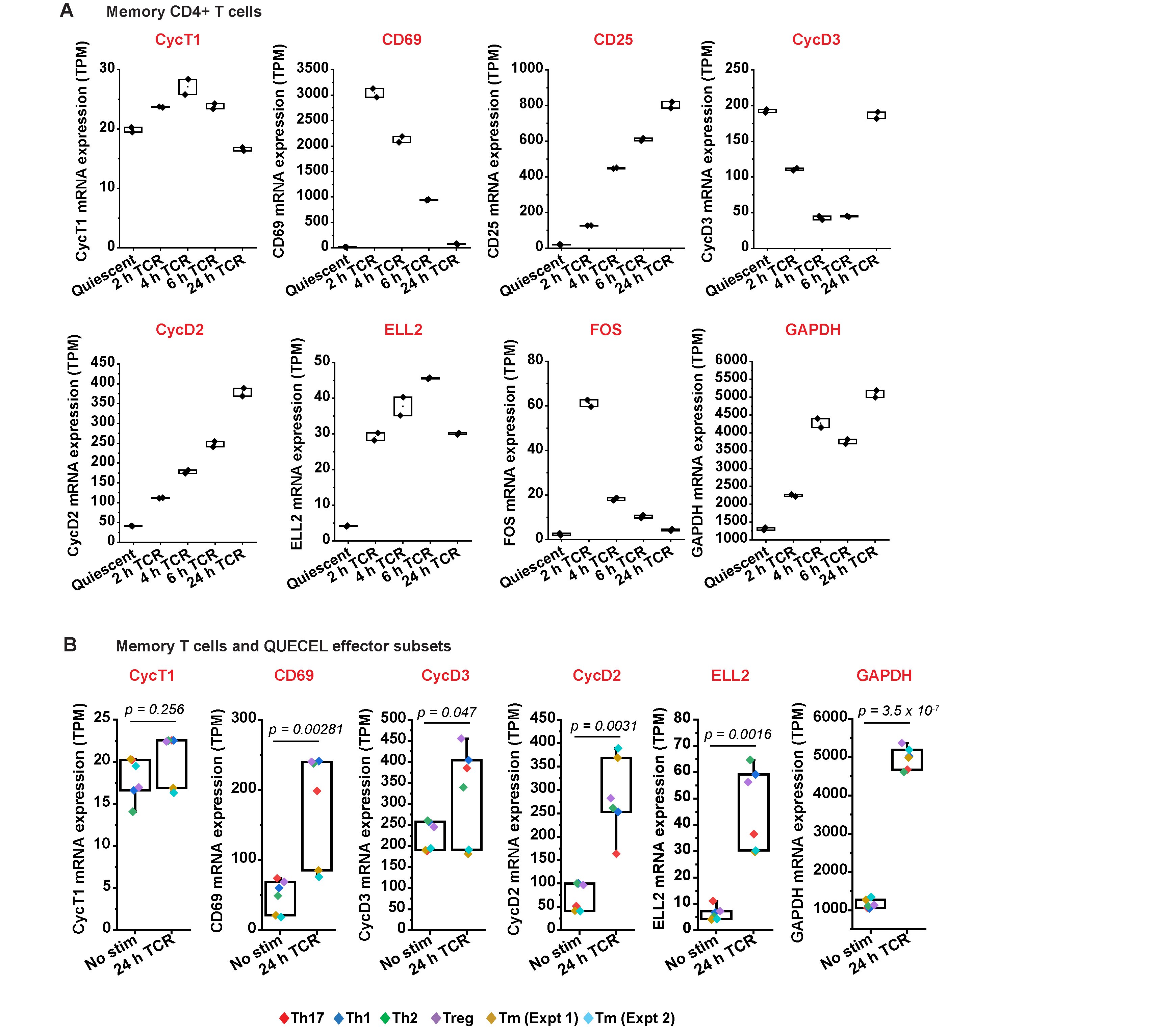

### Supplementary Figure 6

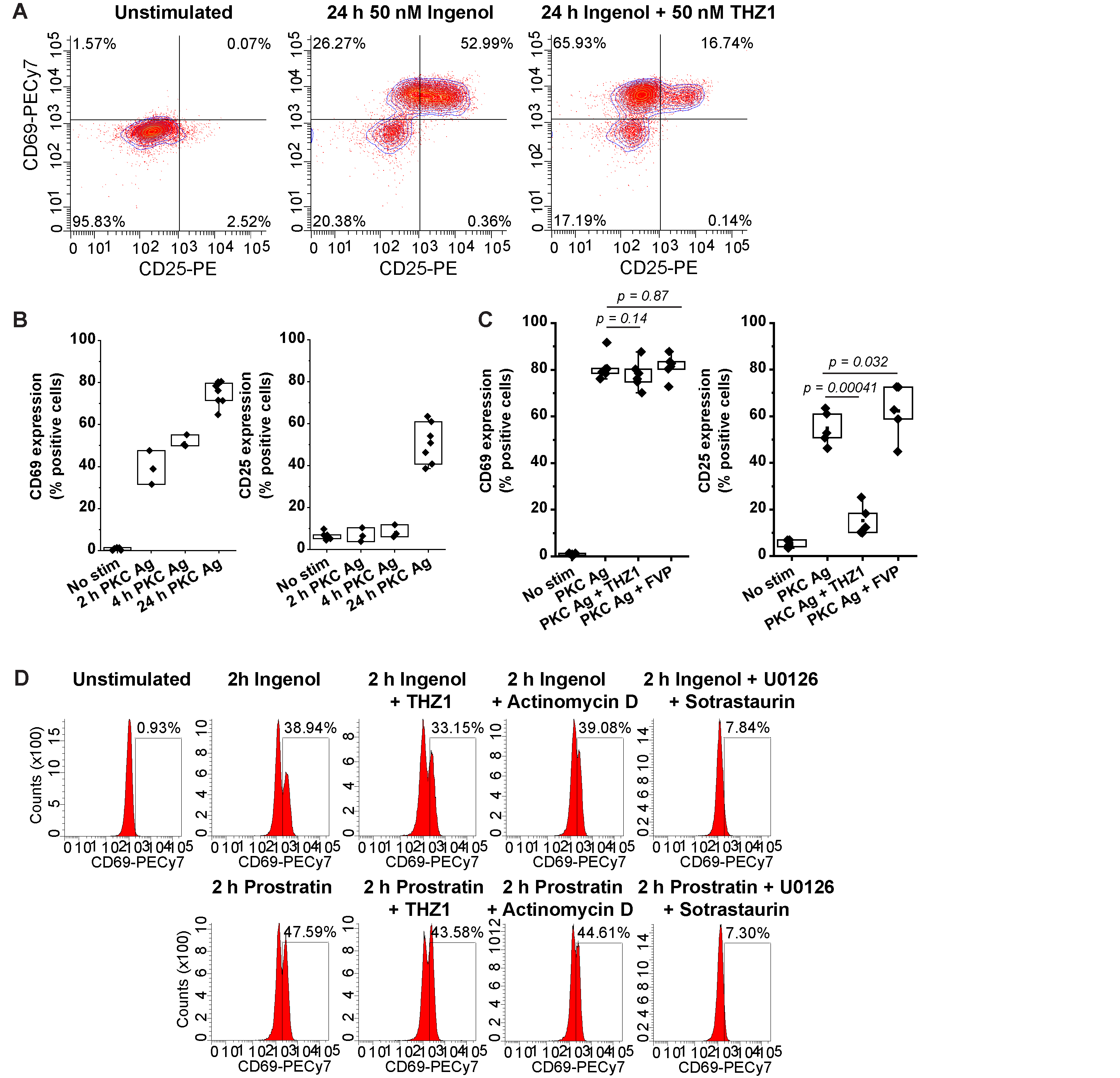

### Supplementary Figure 7

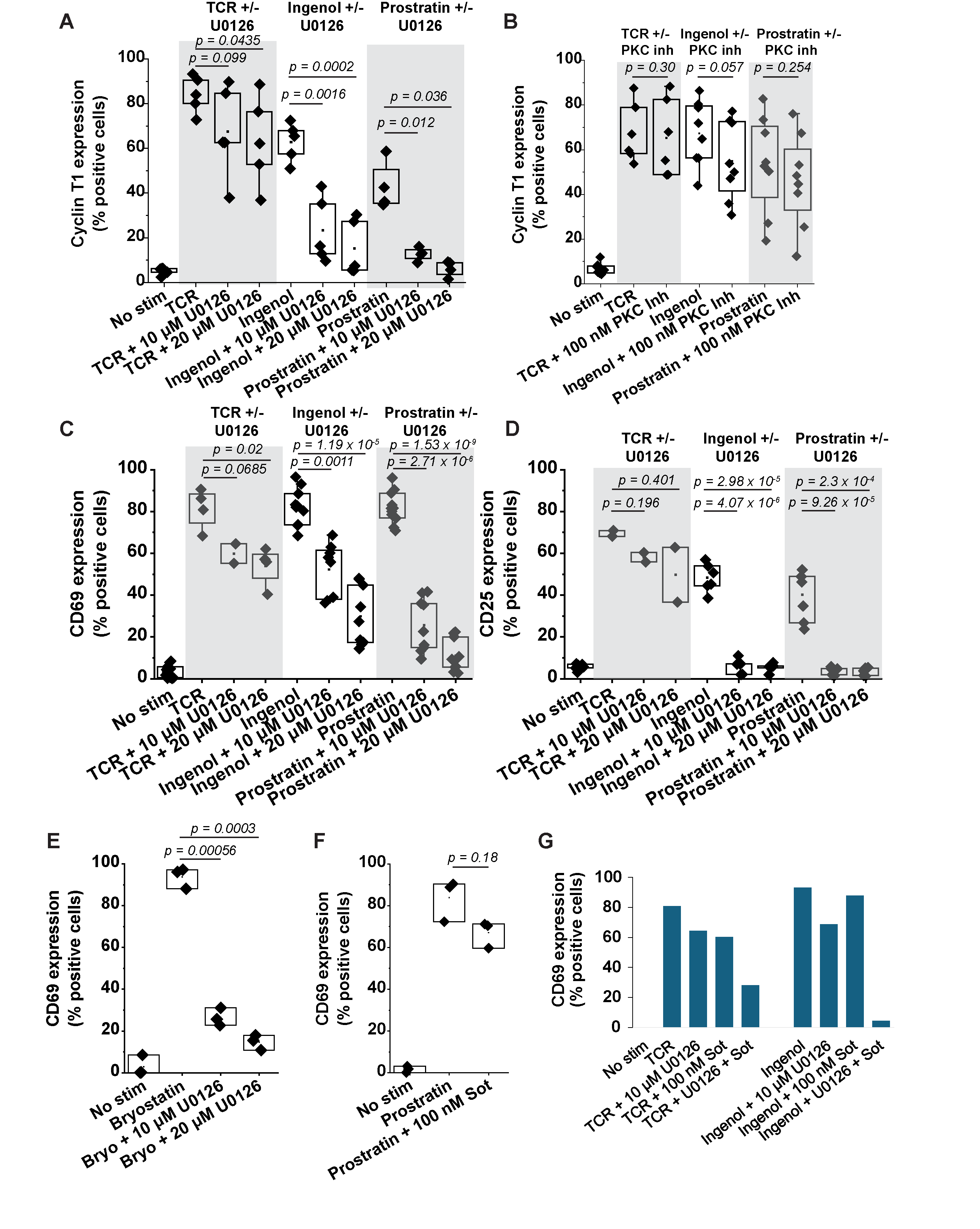

### Supplementary Figure 8

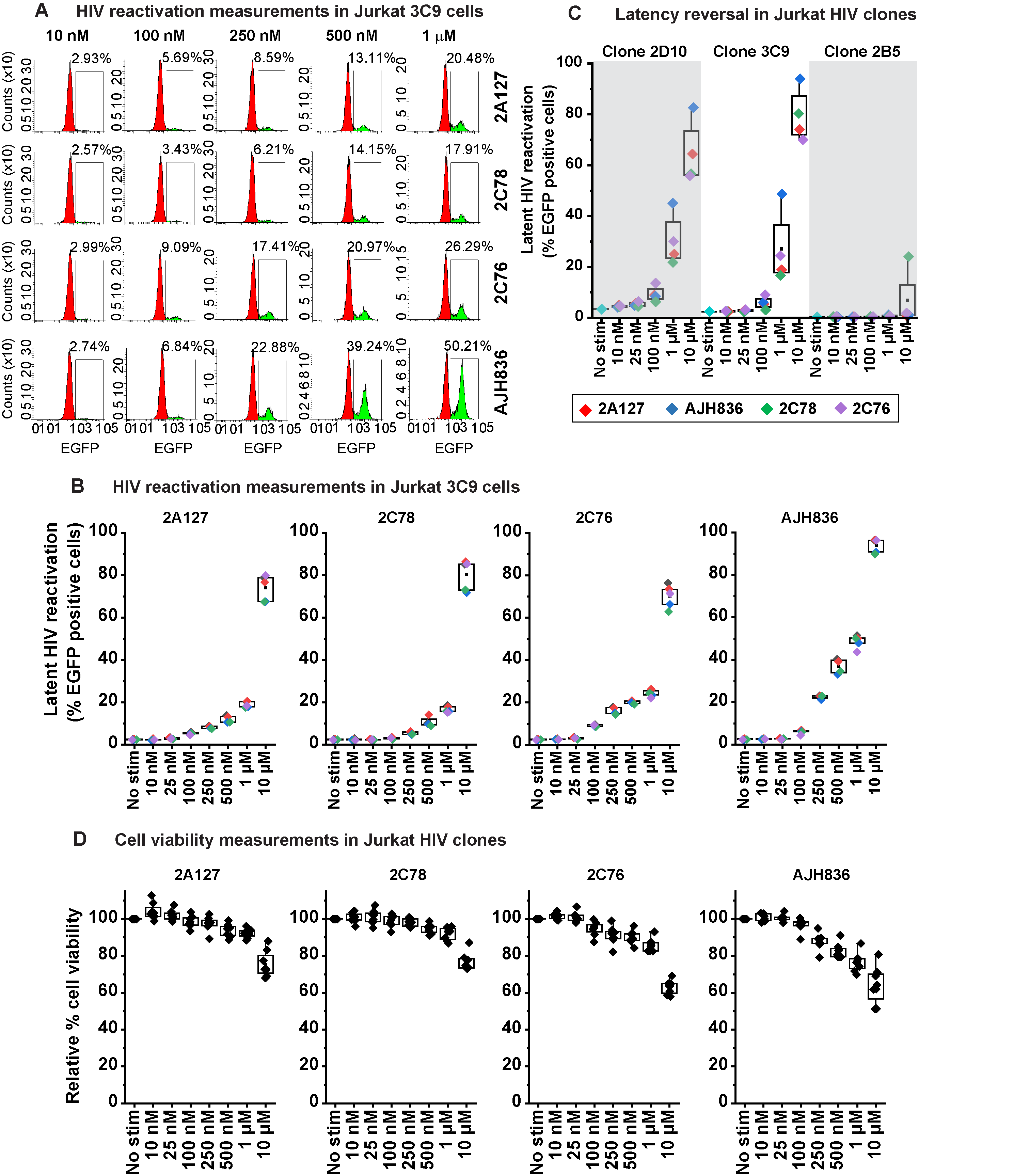

### Supplementary Figure 9

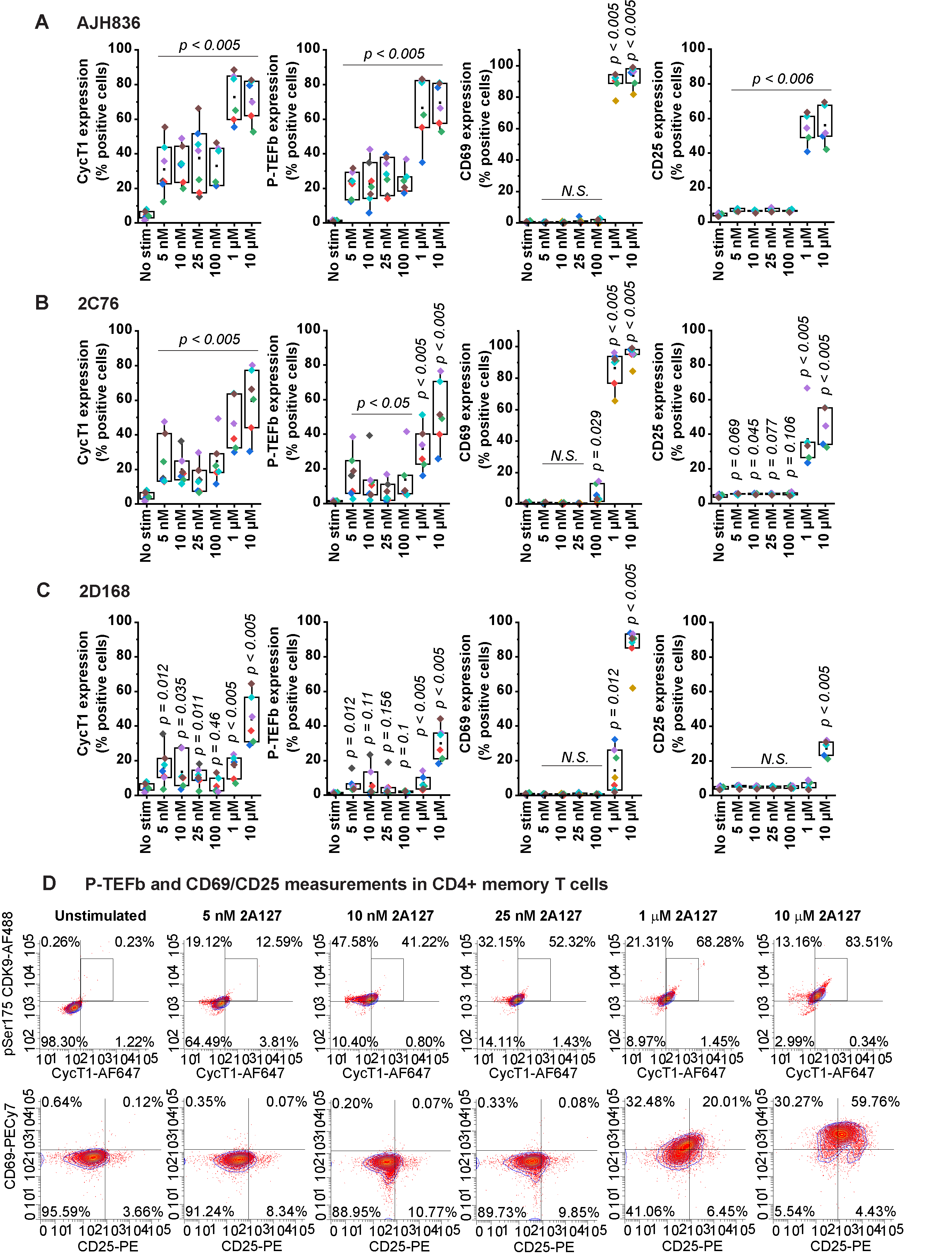

### Supplementary Figure 10

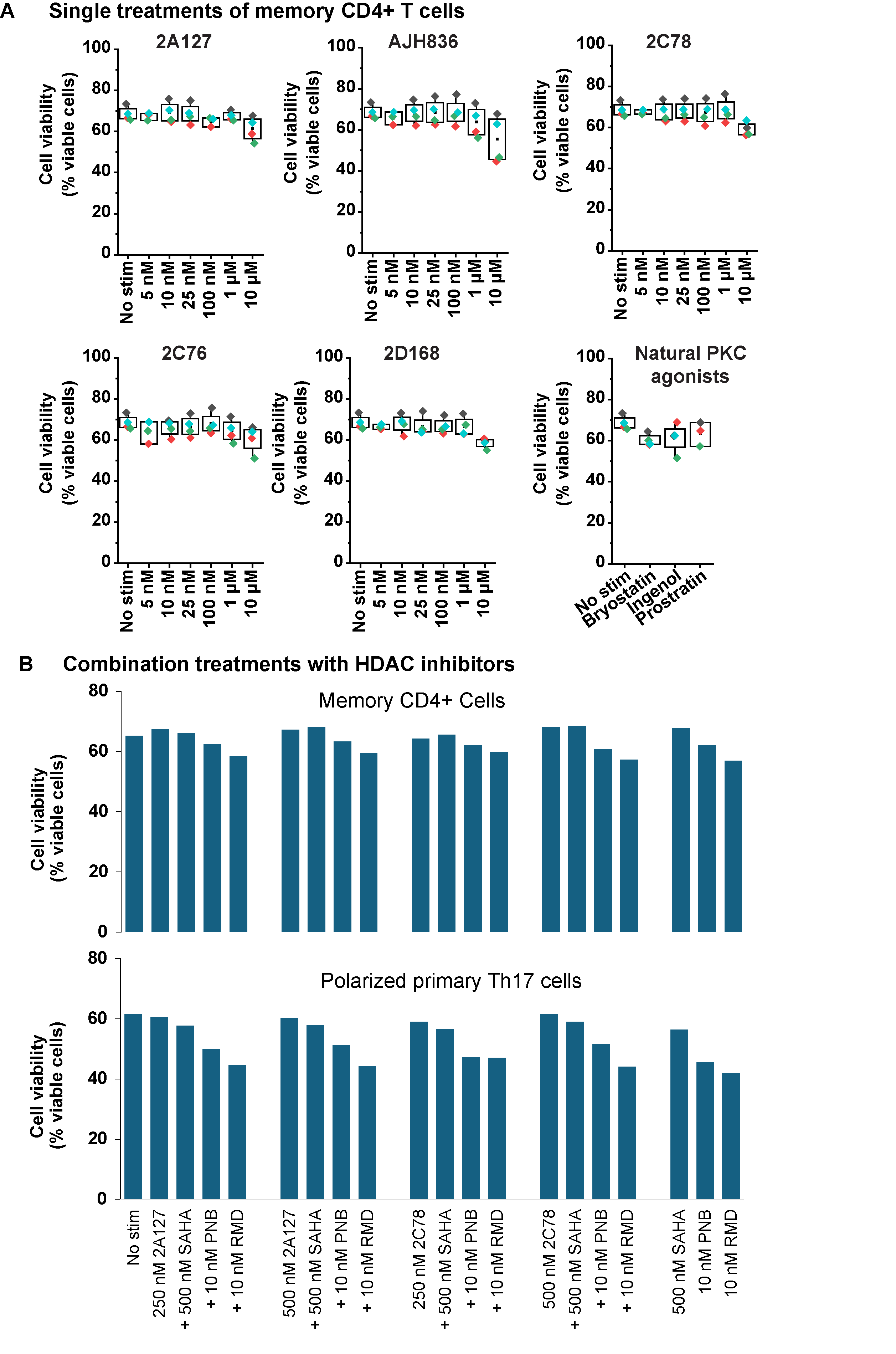

### Supplementary Figure 11A

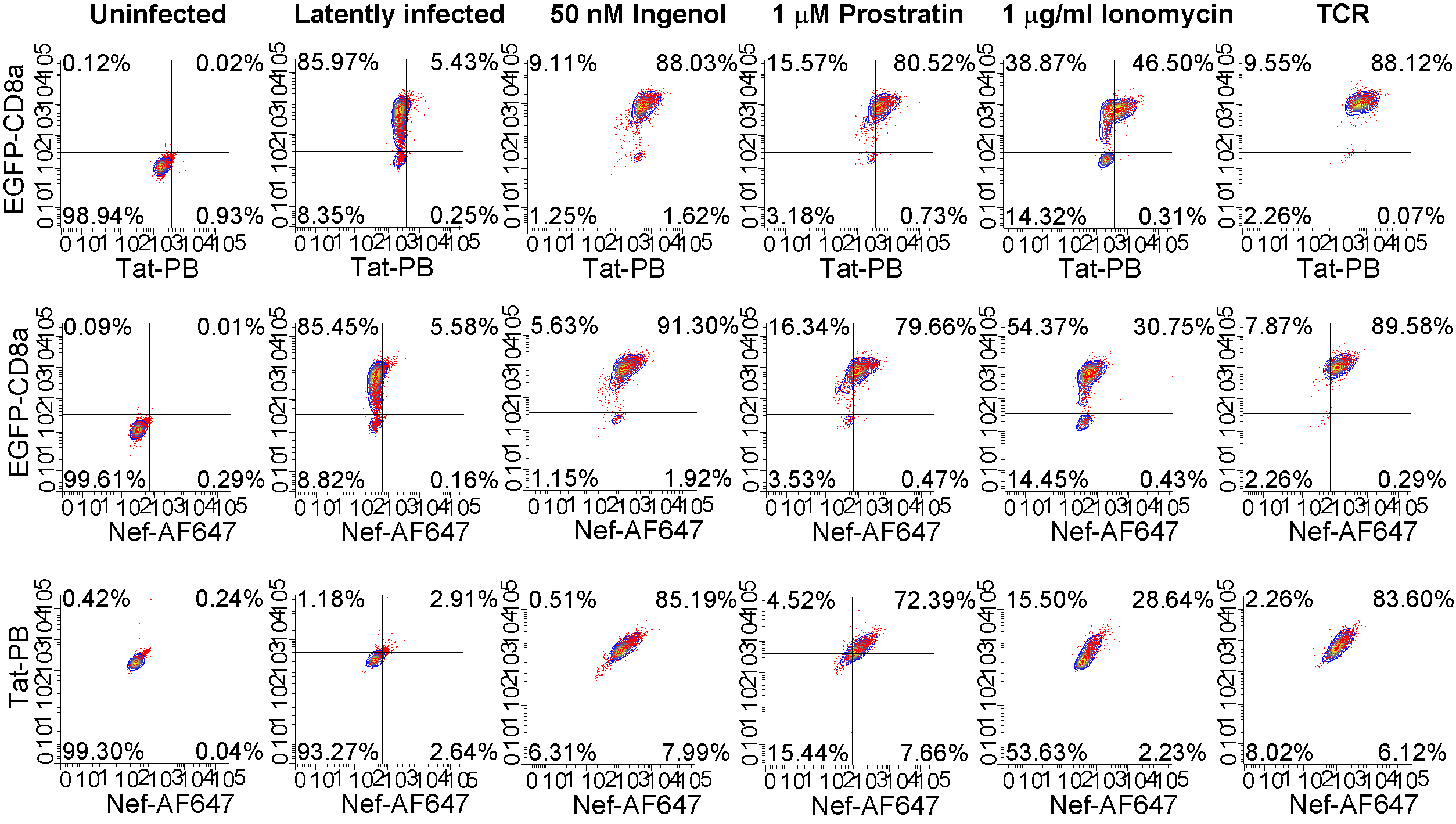

### Supplementary Figure 11B

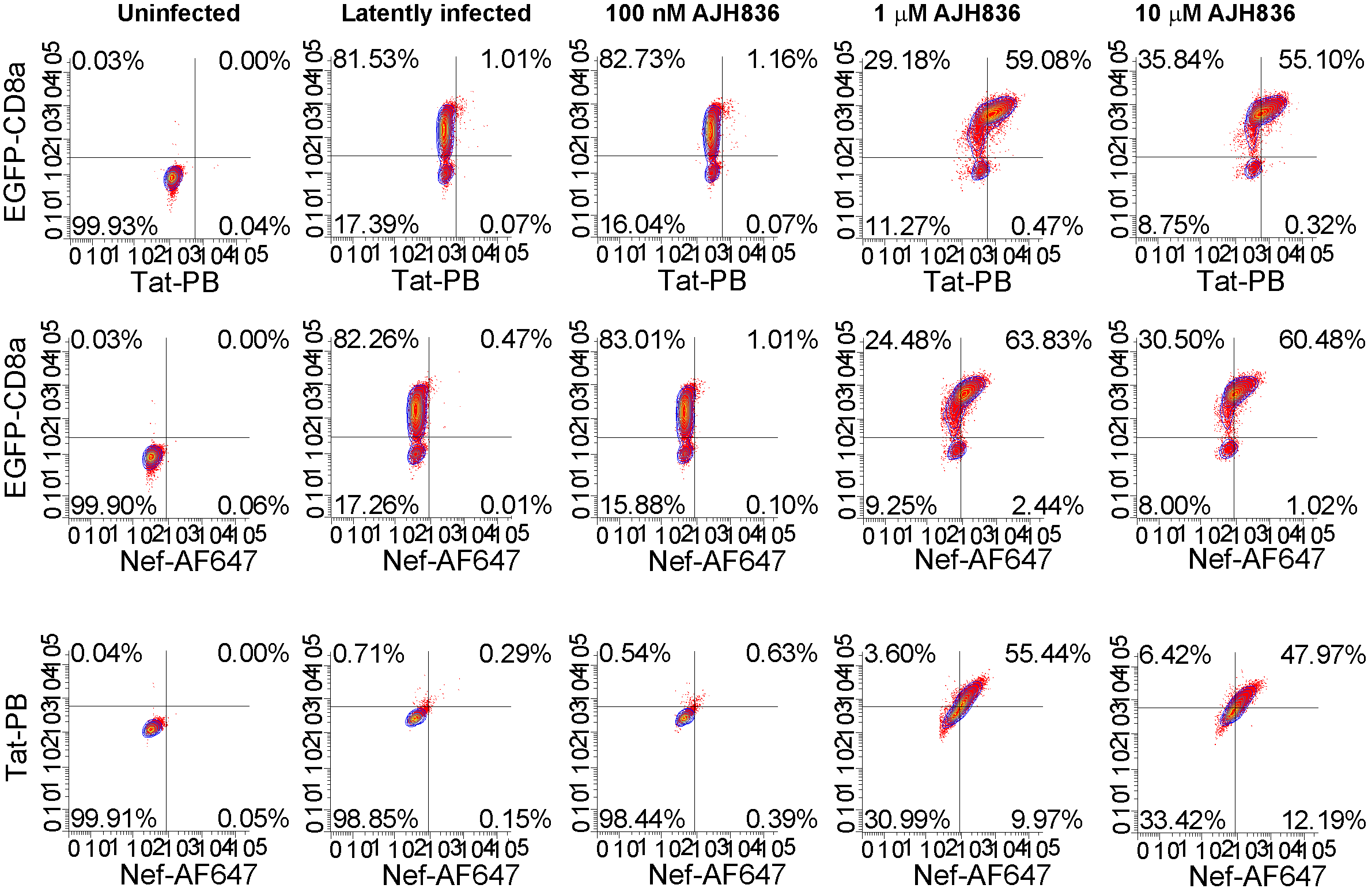

### Supplementary Figure 12

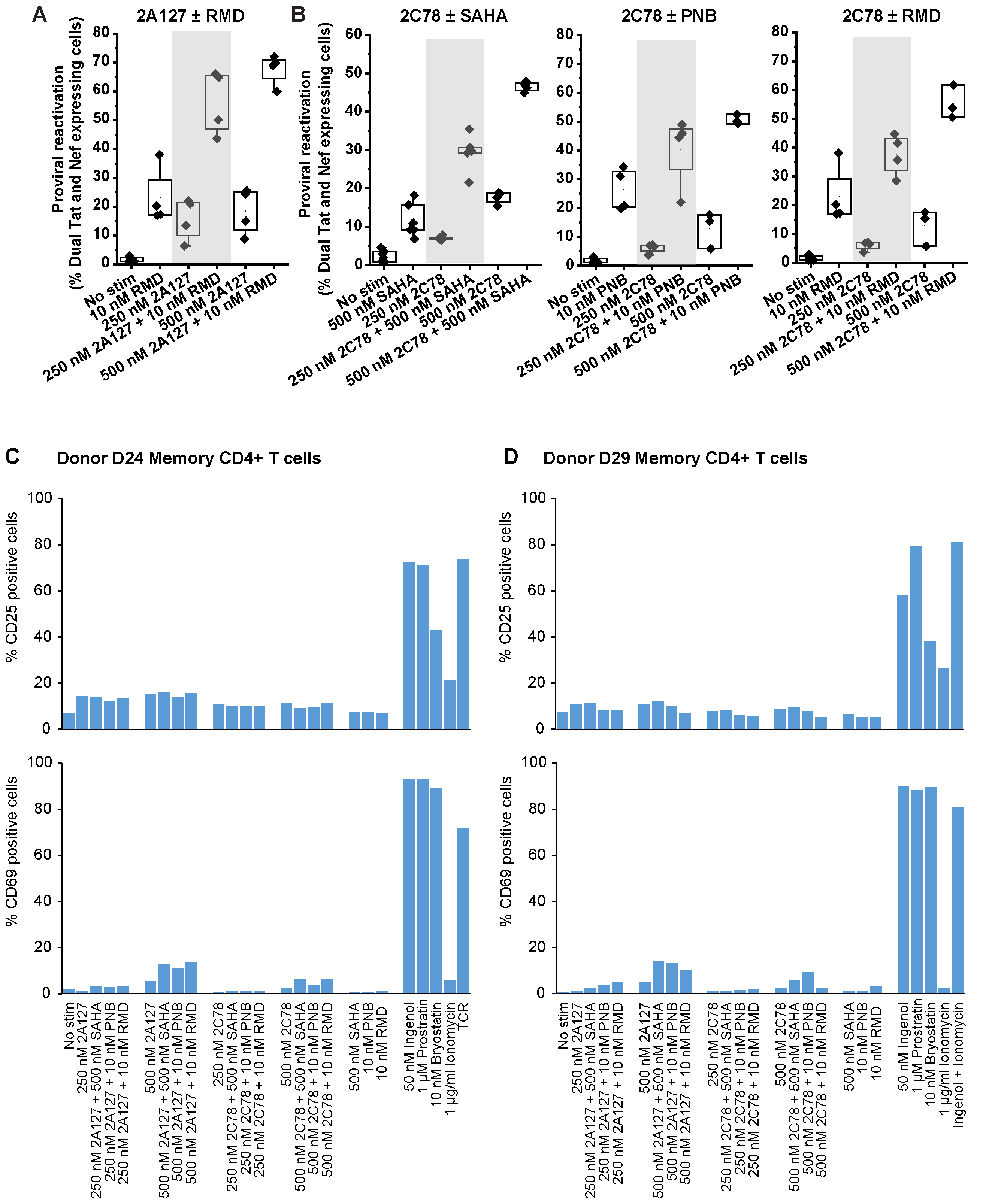

### Supplementary Figure 13

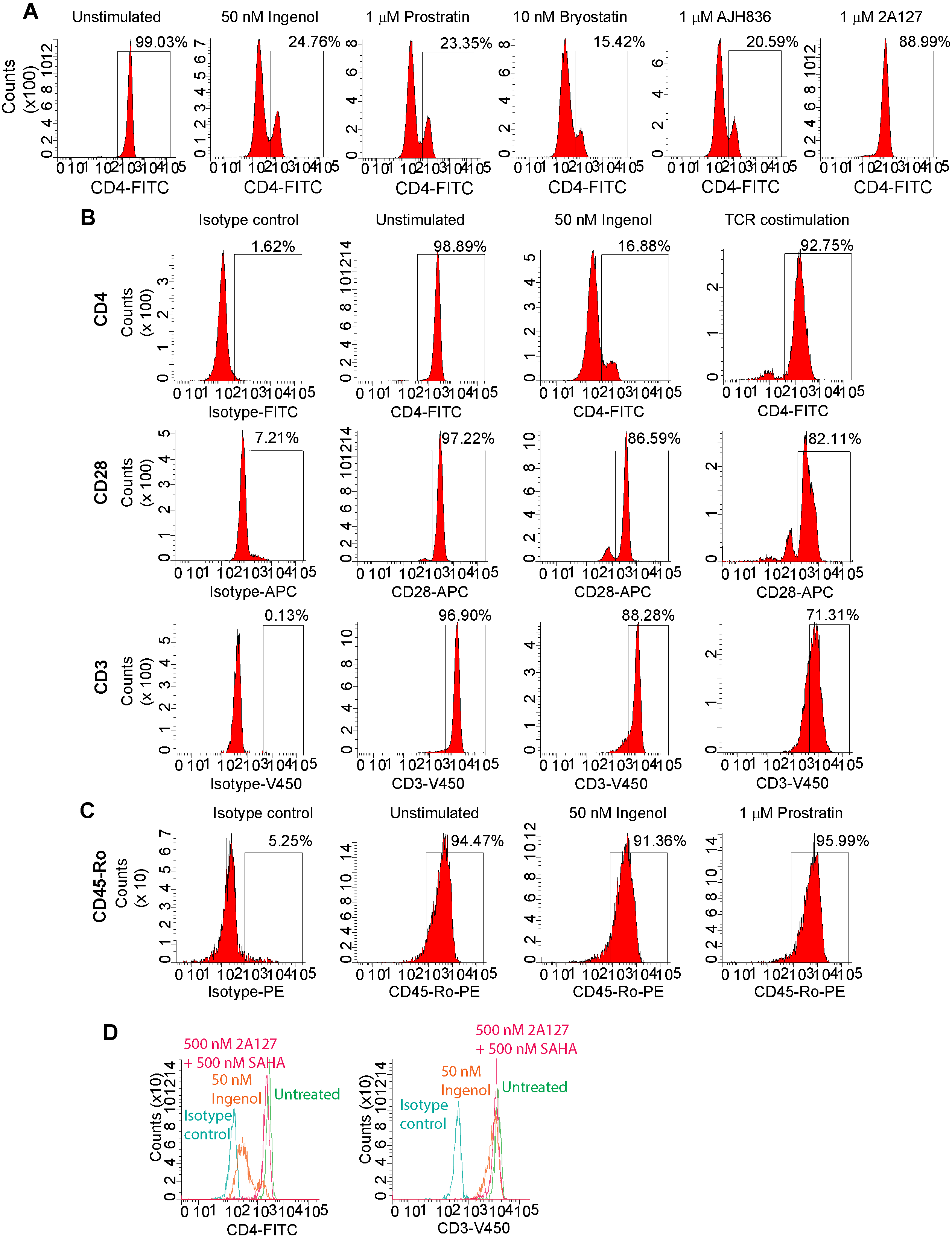

### Supplementary Figure 14

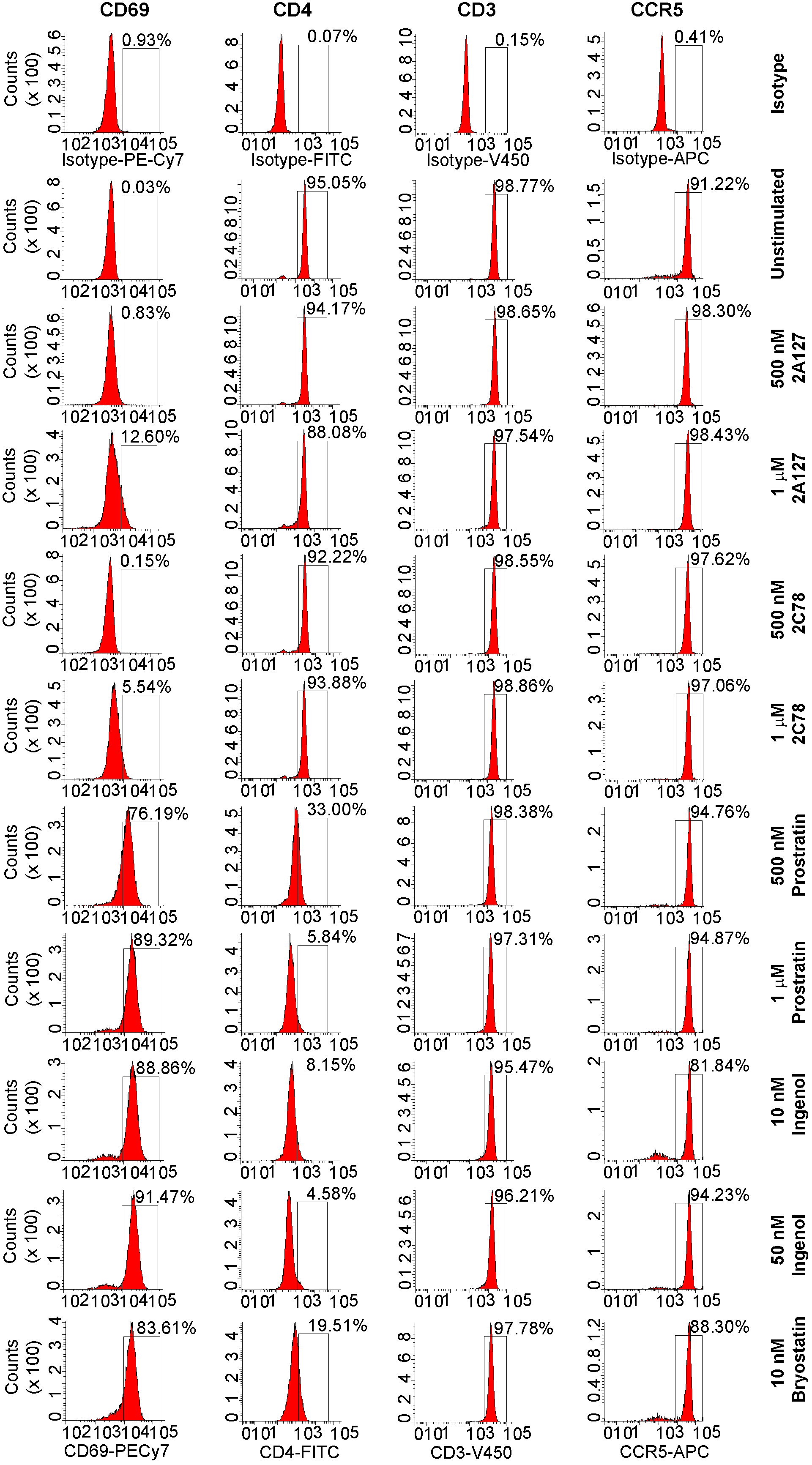

### Supplementary Figure 15

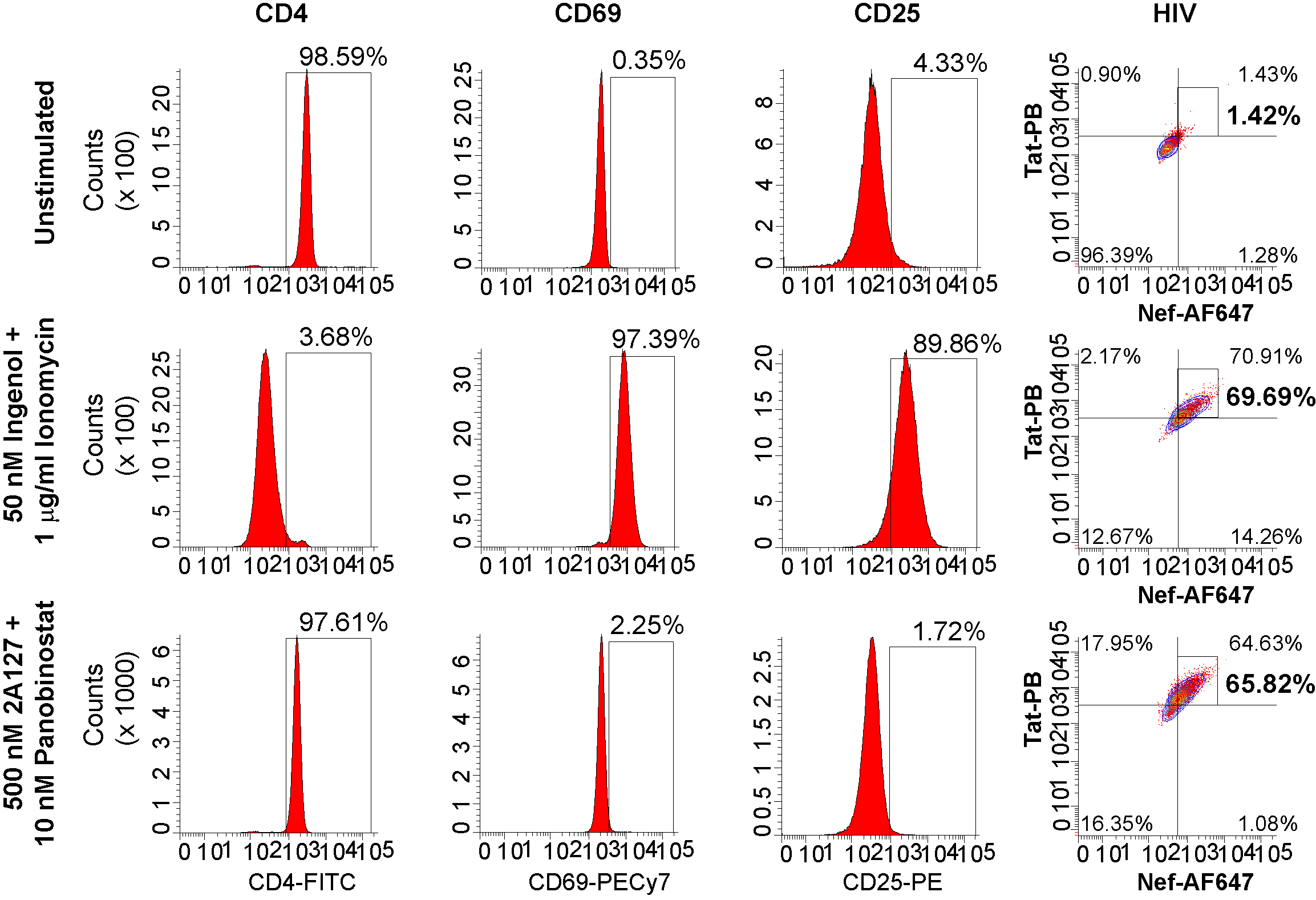

### Supplementary Figure 16

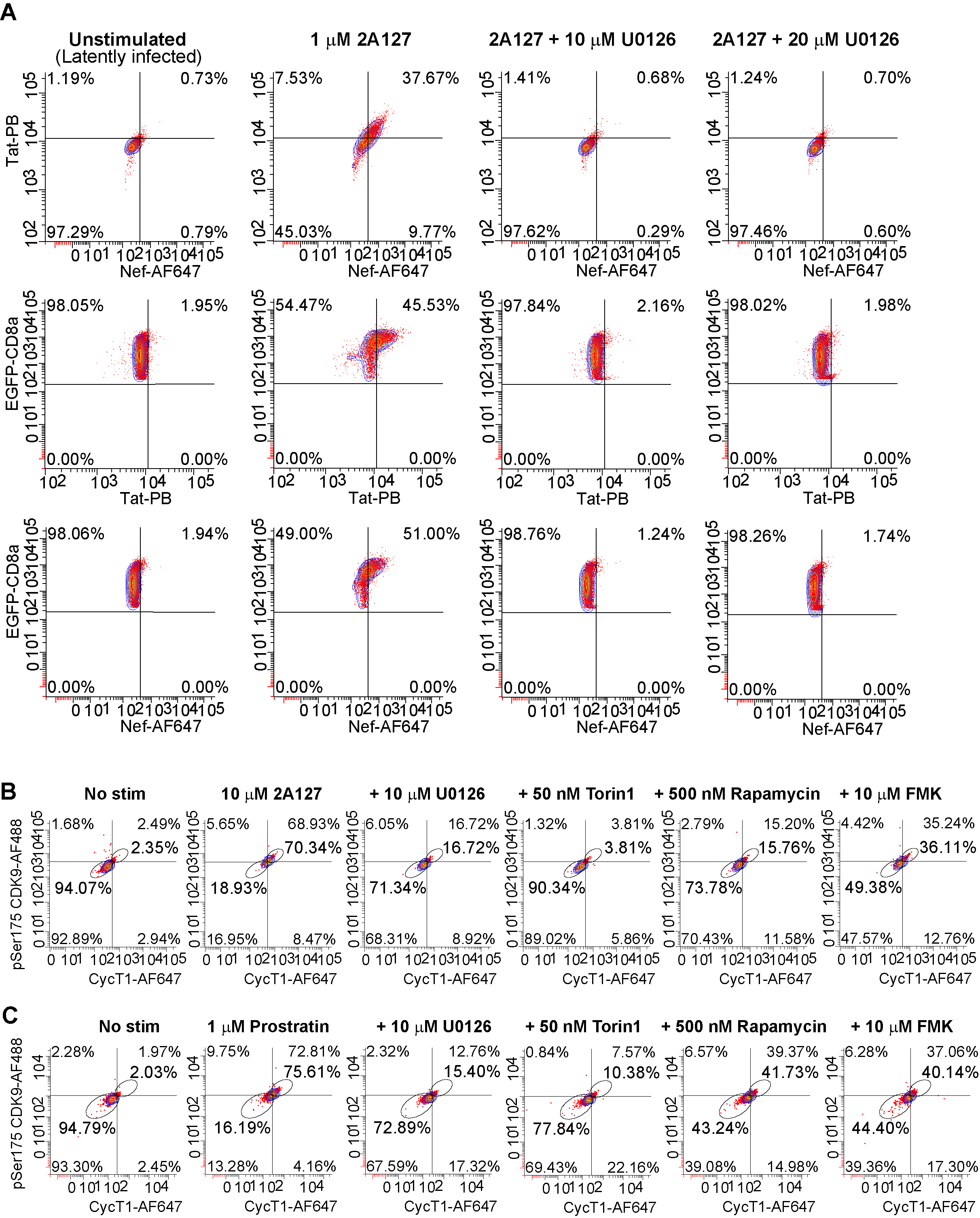

### Supplementary Figure 17

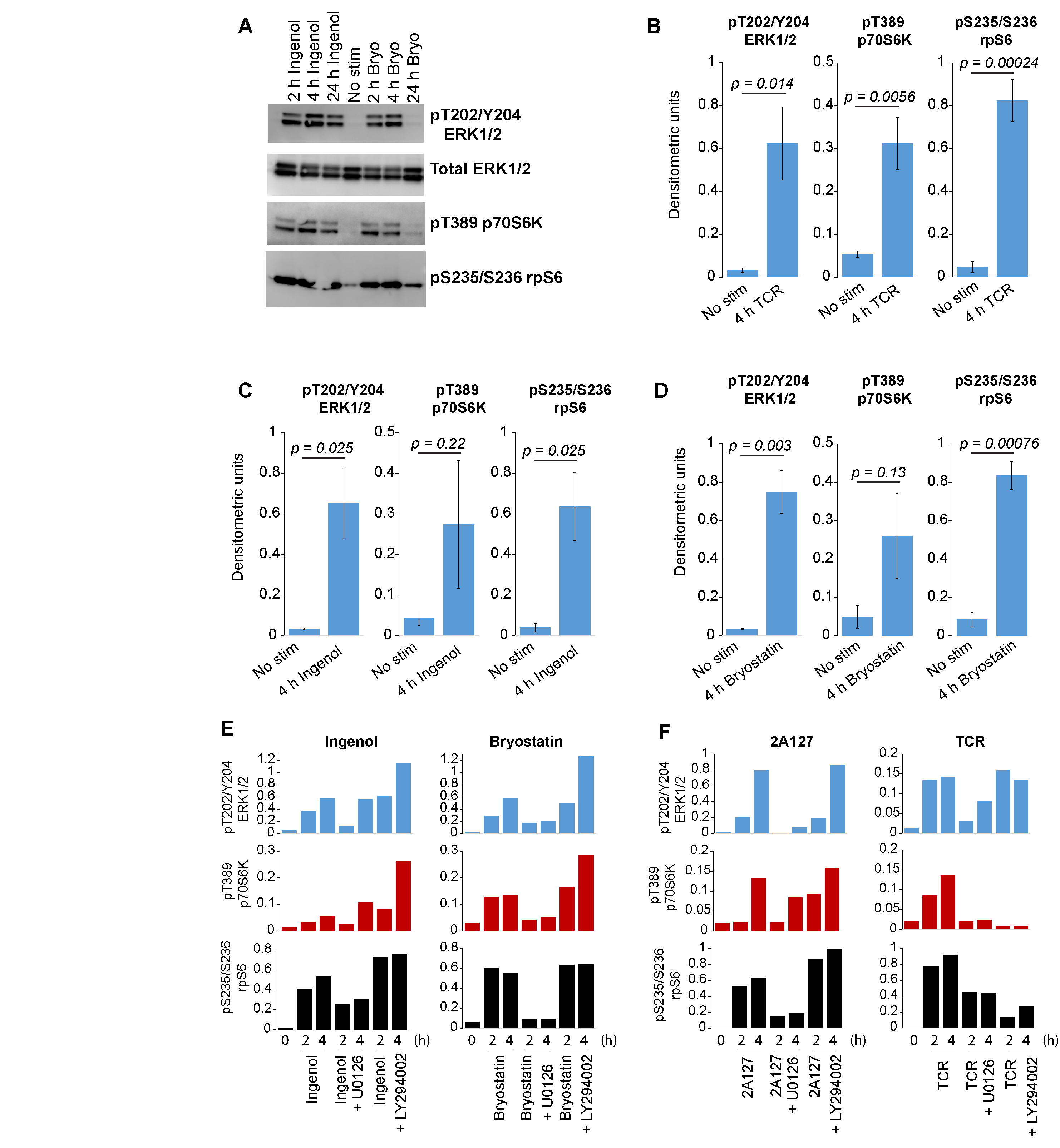

### Supplementary Figure 18

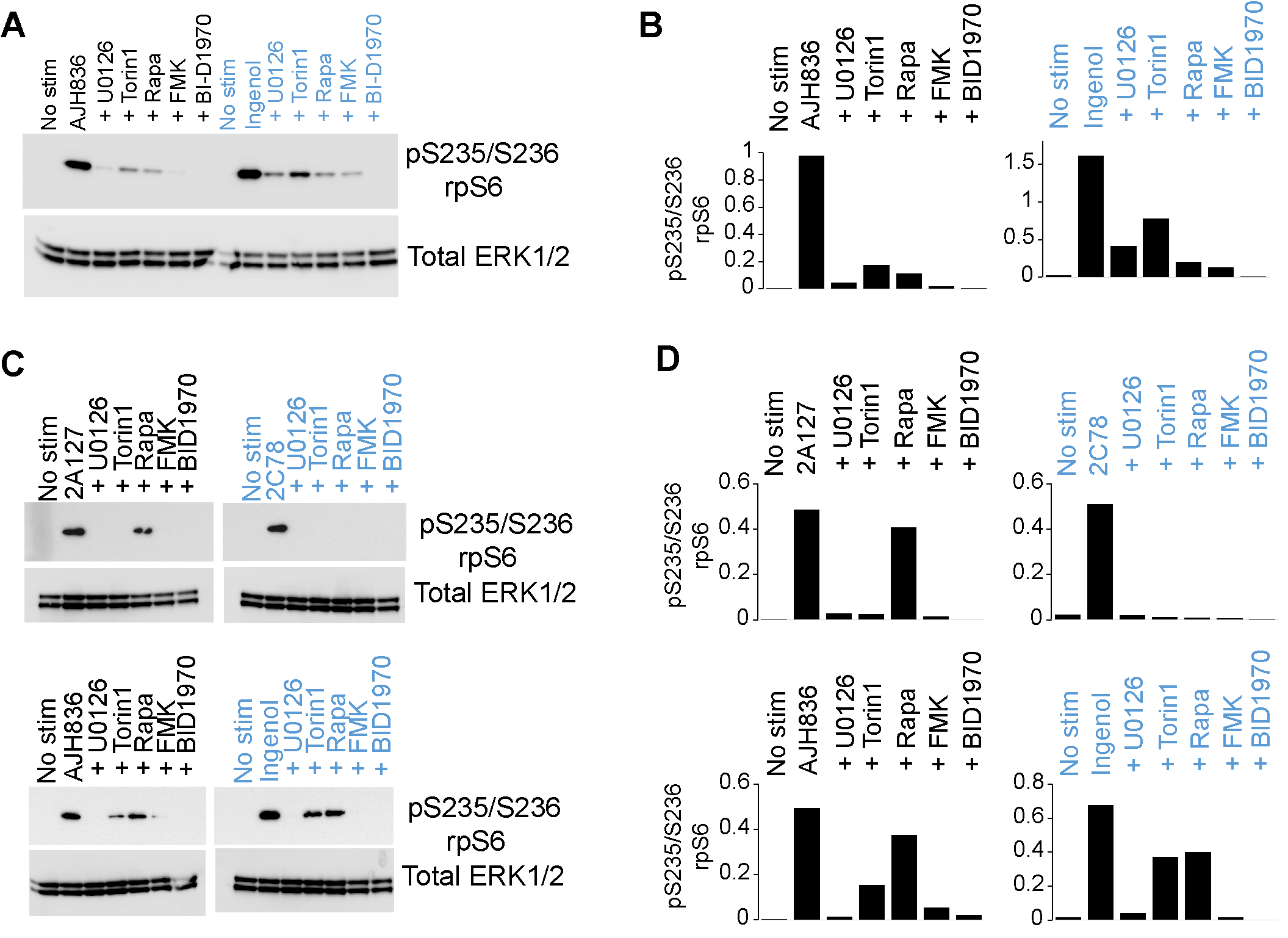

### Supplementary Figure 19

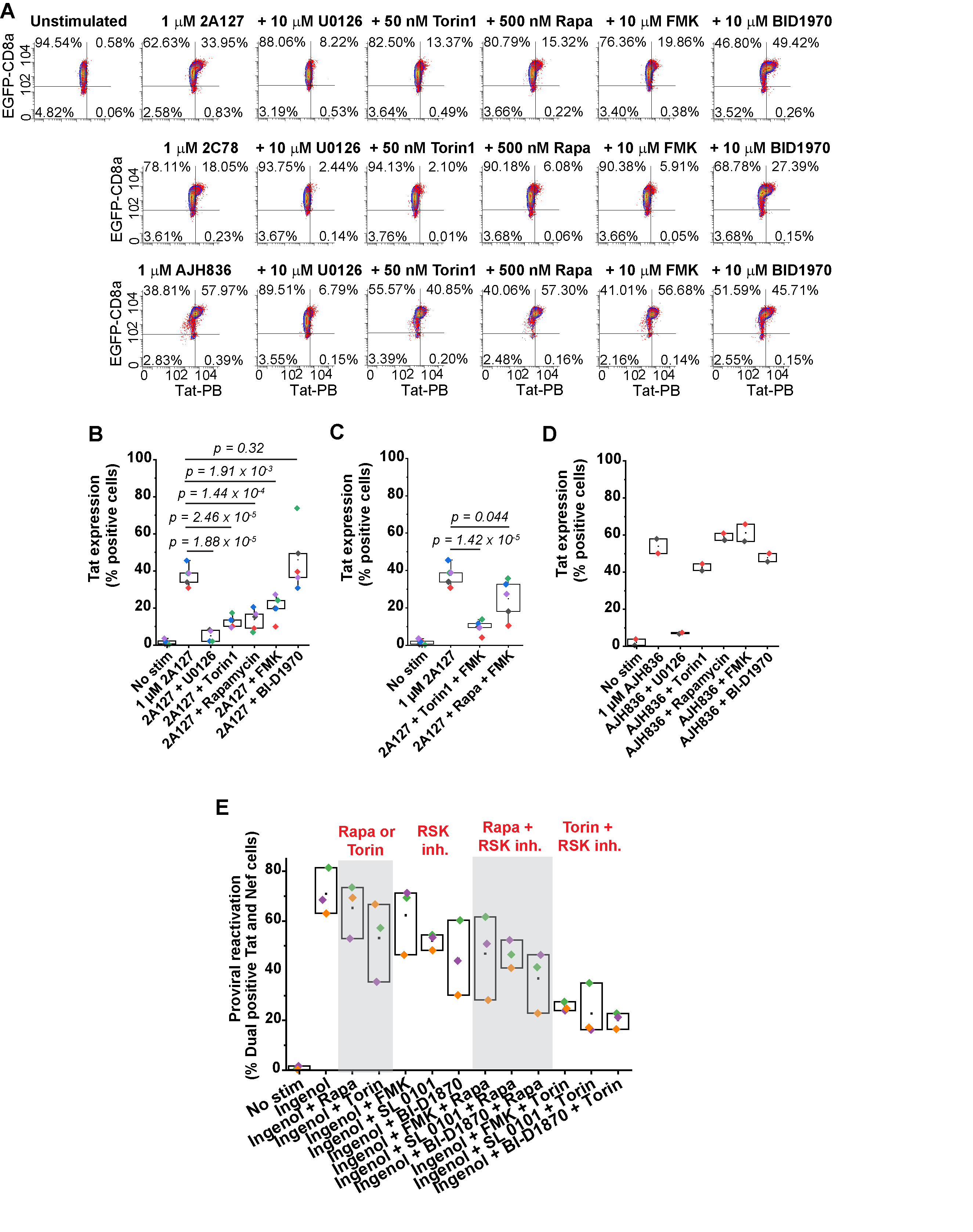

### Supplementary Figure 20

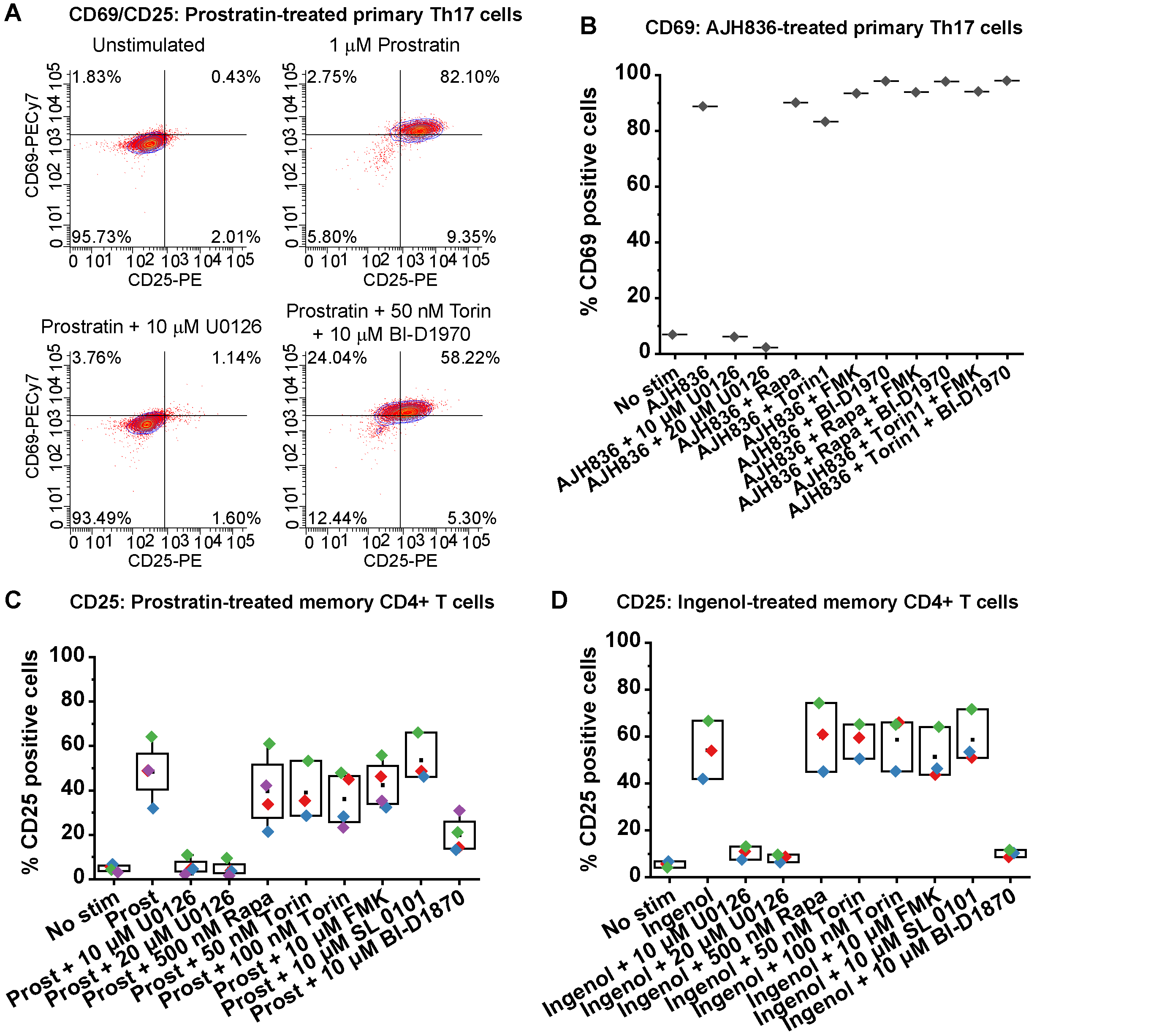

### Supplementary Figure 21

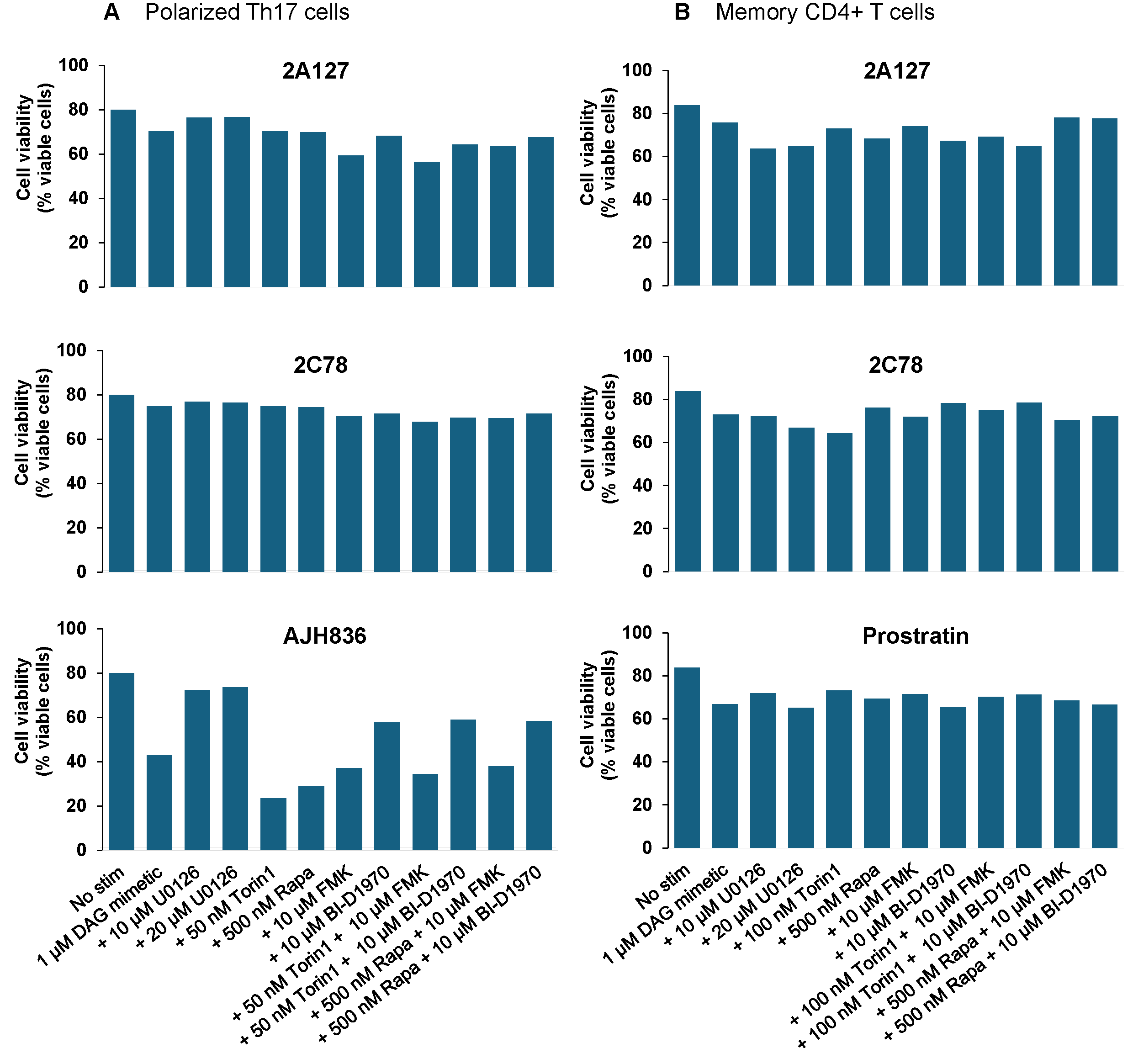

### Supplementary Figure 22A

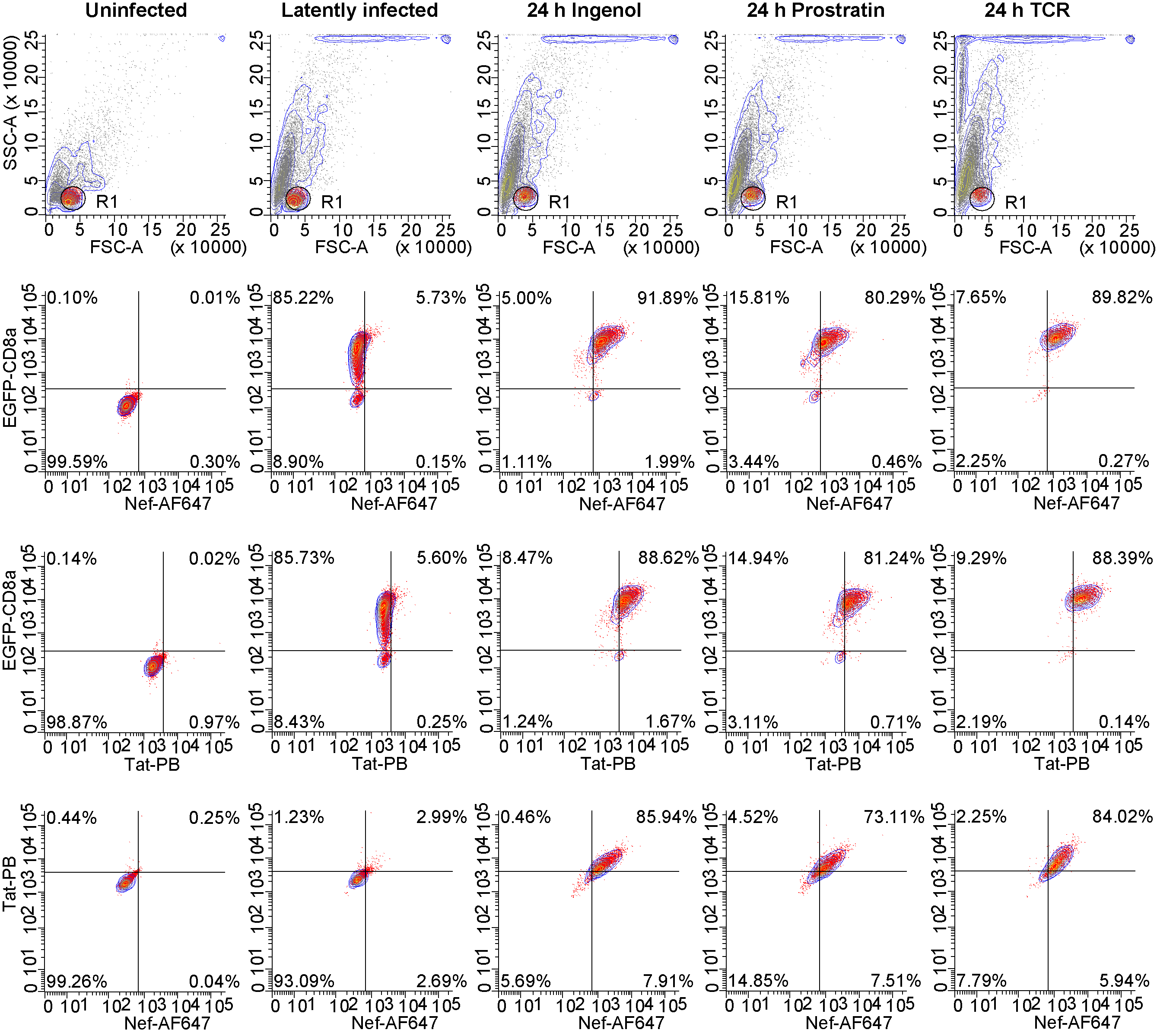

### Supplementary Figure 22B

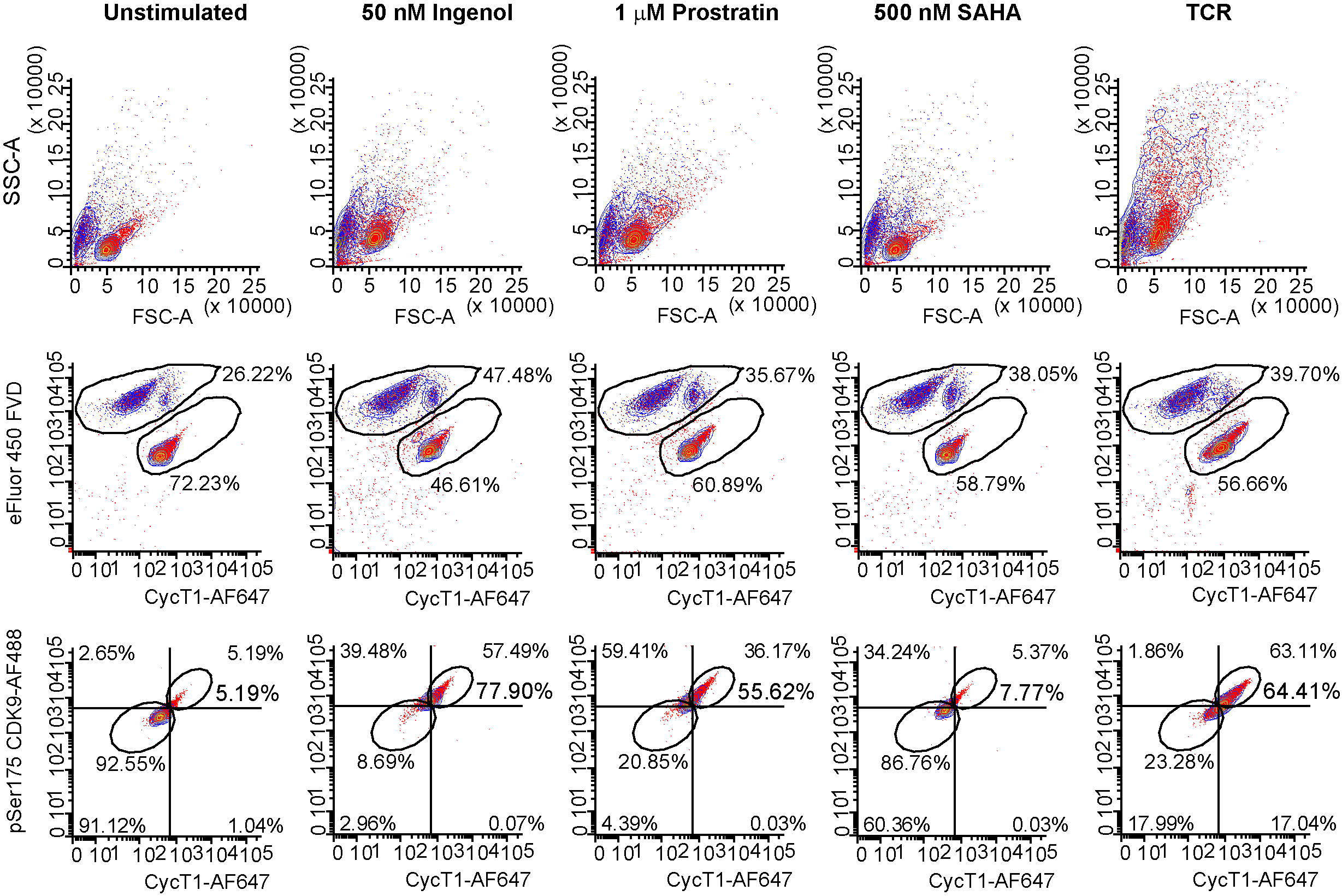

### Supplementary Figure 22C

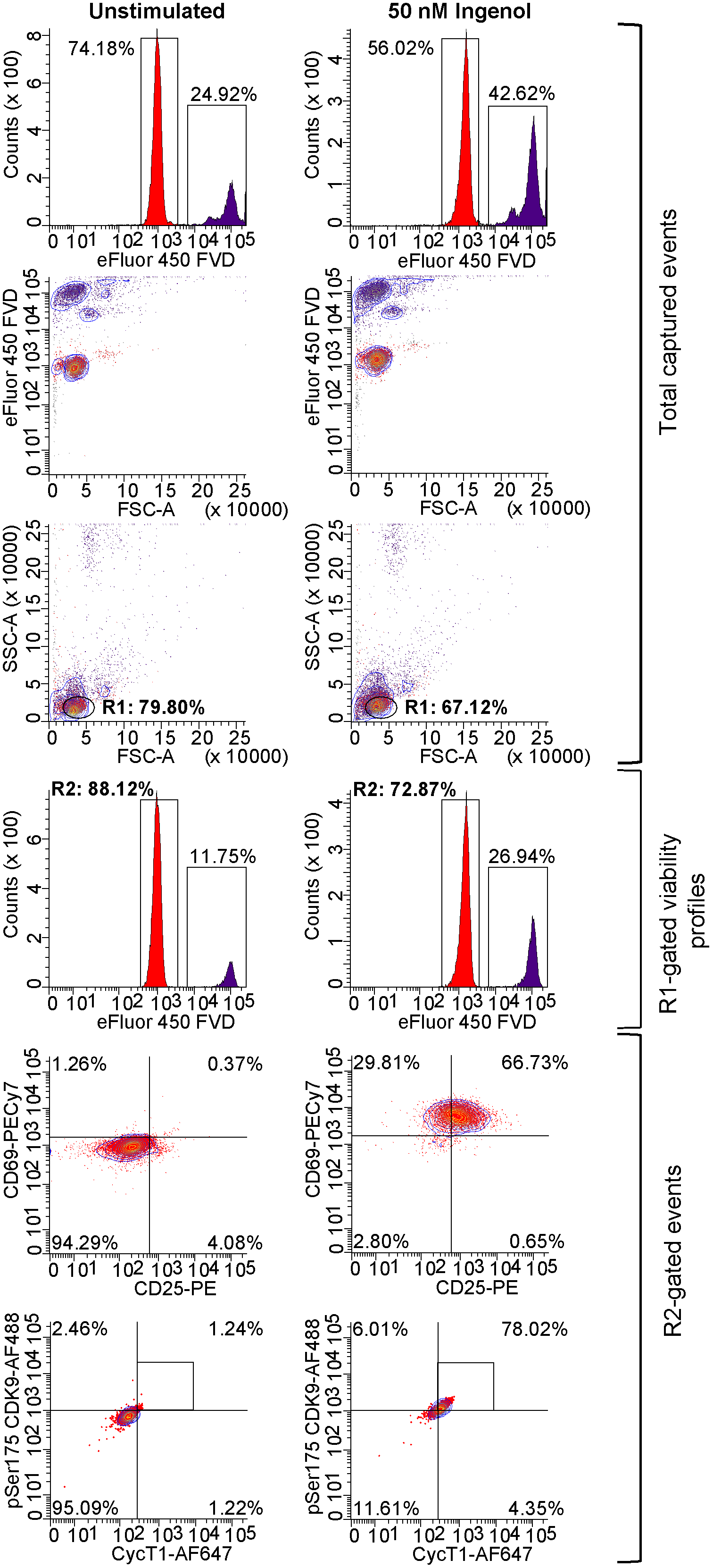

### Supplementary Figure 22D

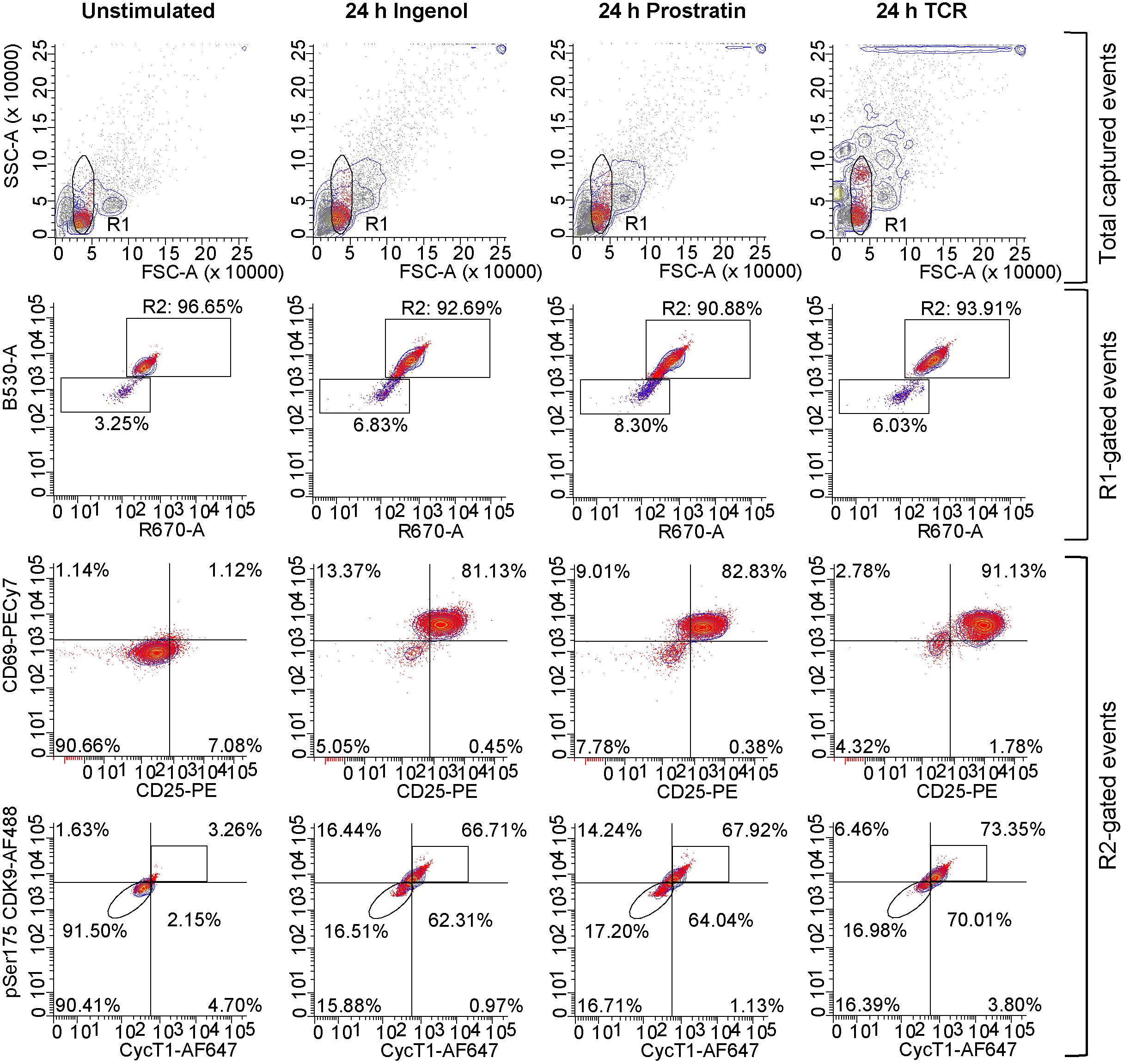

### Supplementary Table 1

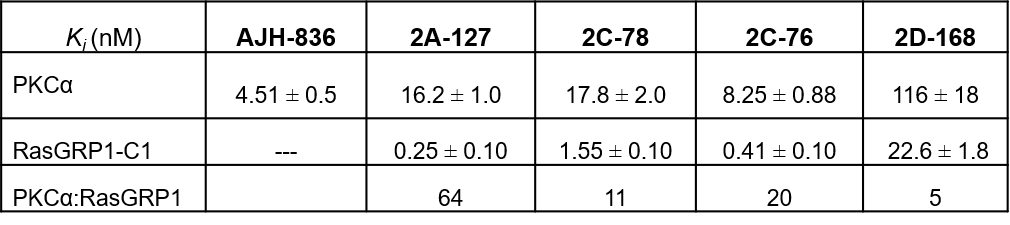

**Supplementary Table 1. Binding selectivity of DAG-indololactones for PKCα vs RasGRP1**

### Supplementary Table 3

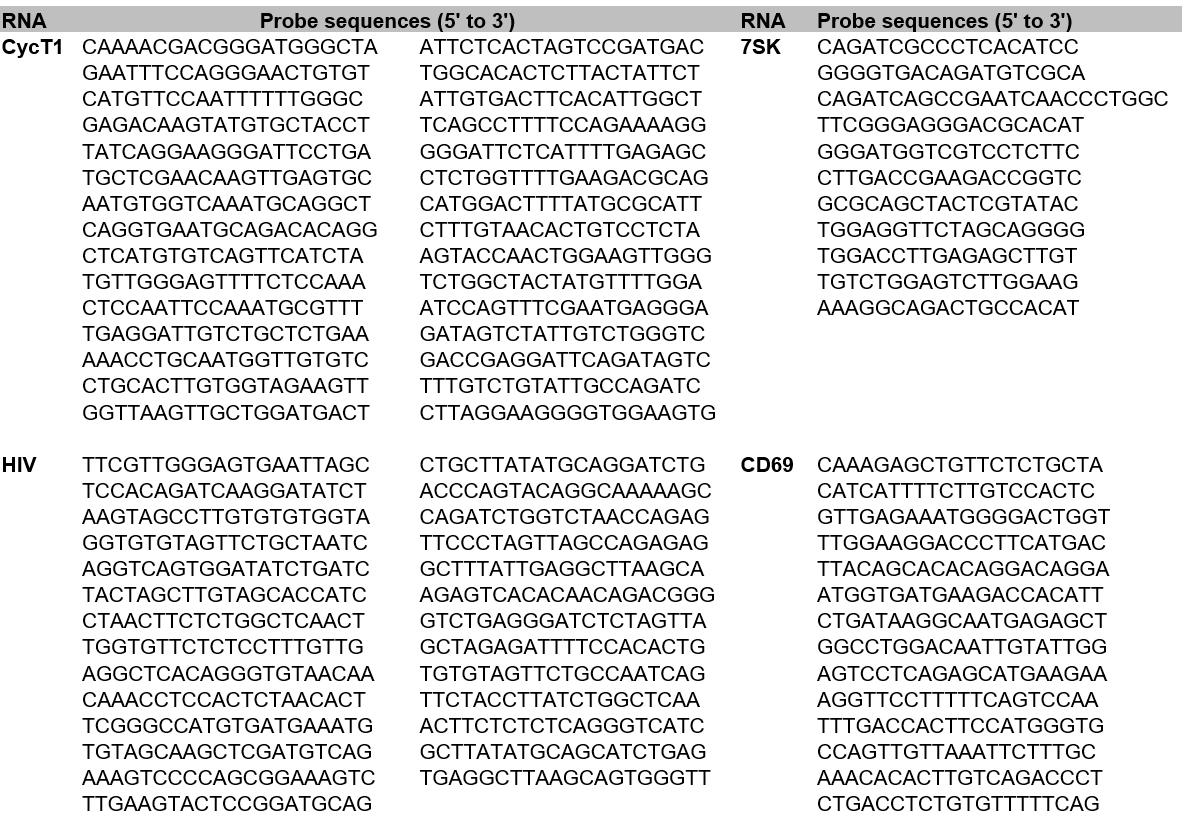
**Supplementary Table 3. Stellaris DNA oligonucleotide RNA FISH probes**
