## Supplementary Table 2 for "RasGRP1 agonism stimulates P-TEFb biogenesis via MEK-ERK-mTORC1 signaling to reverse HIV latency with minimal CD4 downregulation"

**Supplementary Table 2. Key reagents used in the current study**

| **REAGENT** | **SOURCE** | **CATALOG # or Donor #** |
| --- | --- | --- |
| **Primary cells** |  |  |
| Healthy donor PBMCs | AllCells | D9, D10, D14, D15, D16, D17, D18, D23, D24, D25, D26 |
| Healthy donor PBMCs | StemCell Technologies | D27, D29, D30 |
| PBMCs from treated PWH | Rusbelt CFAR Clinical Core | D3214, D6447, D6133, D6404, D6475 |
| **LRA reagents** |  |  |
| Ingenol-3-angelate (ingenol) | Cayman Chemical | 16207 |
| Prostratin | Millipore-Sigma | P0077 |
| Bryostatin 1 | Millipore-Sigma | B7431 |
| BRD4 PROTAC ZXH-3-26 | MedChemExpress | HY-122826 |
| SMAC MIMETIC AZD5582 | Selleckchem | S7362 |
| JQ1 | Millipore-Sigma | SML0974 |
| SAHA | Millipore-Sigma | SML0061 |
| Panobinostat | APEXBIO | A8173 |
| Romidepsin | APEXBIO | A8178 |
| Recombinant human IL-15 | Peprotech | 200-15 |
| AJH836 DAG-lactone | Supplied by INTI-Argentina | N/A |
| 2A127 DAG-indololactone | Synthesized in INTI-Argentina | N/A |
| 2C78 DAG-indololactone | Synthesized in INTI-Argentina | N/A |
| 2C76 DAG-indololactone | Synthesized in INTI-Argentina | N/A |
| 2D168 DAG-indololactone | Synthesized in INTI-Argentina | N/A |
| Ionomycin | Millipore-Sigma | I9657 |
| **Inhibitors** |  |  |
| Rapamycin | Fisher Scientific | AAJ62473MF |
| Torin1 | Selleckchem | S2827 |
| FMK | MedChemExpress | HY-52101A |
| SL 0101 | MedChemExpress | HY-15237 |
| BI-D1870 | MedChemExpress | HY-10510 |
| THZ1 | Millipore-Sigma | 532372001 |
| Flavopiridol | Millipore-Sigma | F3055 |
| Ro-31-8220 | Millipore-Sigma | 557520 |
| Sotrastaurin | Selleckchem | S2791 |
| U0126 | Millipore-Sigma | 662005 |
| LY294002 | Selleckchem | S1105 |
| Actinomycin D | Selleckchem | S8964 |
| MG132 | Selleckchem | S2619 |
| Cycloheximide | Millipore-Sigma | 239763-M |
| **Antibodies** |  |  |
| Cyclin T1 | ThermoFisher | PA5-77892 |
| pSer175 CDK9 | Custom-synthesized | N/A |
| pSer2 RNAP II CTD | Abcam | 5095 |
| pThr202/pTyr204 ERK1/2 | Cell Signaling Tech | 4370 |
| ERK1/2 | Cell Signaling Tech | 9102 |
| pThr389 P70S6K | Cell Signaling Tech | 9234 |
| pSer235/p236 S6 | Cell Signaling Tech | 4858 |
| Nef | Abcam | ab42355 |
| Tat | Custom-synthesized | NT3 2D1.1 |
| Mouse IgG blocking antibody | ThermoFisher | 10400C |
| **Antibody conjugates** |  |  |
| CD69-PECy7 | BD Biosciences | 557745 |
| CD69-APC | BD Biosciences | 555533 |
| CD25-PE | BD Biosciences | 555432 |
| CCR5-APC | BD Biosciences | 560748 |
| CD28-V450 | BD Biosciences | 561397 |
| CD45-RO-PE | BD Biosciences | 347967 |
| CD3-V450 | BD Biosciences | 560365 |
| CD4-Pacific Blue | BD Biosciences | 558116 |
| CD4-FITC | BD Biosciences | 550628 |
| Cyclin D3-PE | Biolegend | 684904 |
| anti-Rabbit IgG AF488 | ThermoFisher | A11008 |
| anti-Mouse IgG Cy5 | Abcam | AB97037 |
| Mouse Isotype IgG PE | Biolegend | 400112 |
| Mouse Isotype IgG FITC | BD Biosciences | 554679 |
| Mouse Isotype IgG PECy7 | Biolegend | 400126 |
| Mouse Isotype IgG V450 | BD Biosciences | 560373 |
| Mouse Isotype IgG Pacific Blue | Biolegend | 400331 |
| Mouse Isotype IgG APC | Biolegend | 400120 |
| **Antibody conjugation kits** |  |  |
| Pacific Blue antibody labeling kit | ThermoFisher | P30013 |
| Alexa Fluor 488 antibody labeling kit | ThermoFisher | A20181 |
| Alexa Fluor 647 antibody labeling kit | ThermoFisher | A20186 |
| **Primary cell isolation kits** |  |  |
| Memory CD4+ T cell enrichment kit | StemCell Technologies | 19157 |
| Naïve CD4 T cell isolation kit | Biolegend | 480042 |
| Mouse CD8a positive selection kit | StemCell Technologies | 18953 |
| **Other reagents** |  |  |
| PhosSTOP | Roche | 4906837001 |
| EDTA-free protease inhibitors | Roche | 11836170001 |
| Stellaris RNA FISH probes | LGC Biosearch Technologies | Custom-synthesized with Quasar 670 or TAMRA dye |
| Alexa Fluor 594 Phalloidin | ThermoFisher | A12381 |
| eFluor 450 Fixable Viability Dye | ThermoFisher | 65-0863-18 |
| Propidium iodide | Cayman Chemical | 10008351 |
